# Organizing principles of the cellular electron economy

**DOI:** 10.64898/2026.10.02.755995

**Authors:** Pierre Millard, Jean-Charles Portais

**Affiliations:** TBI, Université de Toulouse, CNRS, INRAE, INSA, Toulouse, France; MetaboHUB-MetaToul, National Infrastructure of Metabolomics and Fluxomics, Toulouse, France; University Toulouse, CNRS, INSERM, EFS, ENVT, RESTORE Institute, Toulouse, France

## Abstract

Life runs on electron flow. Cells extract electrons from nutrients and transfer them through metabolism, coupling carbon use to energy conservation, biosynthesis and redox homeostasis. Yet how and even whether electron flow is organized at the system-level across metabolism remain unknown. Here we formalize the concept of the cellular electron economy: a quantitative, genome-wide, reaction-resolved representation of electron generation, transfer and disposal. This electron-centric perspective reveals a quantitative relationship governed by two distinct organizing factors: the carbon source sets the baseline and maximal rate of electron flow, whereas the capacity to dispose of electrons to terminal sinks determines where within this range the system operates. Carbon uptake can even increase as net electron flux falls. This system-level organization is shown to apply to five microorganisms with different intracellular electron routing pathways and predicts graded physiological responses to oxygen supply across carbon sources and respiratory architectures. The cellular electron economy thus links reaction-level redox chemistry to whole-cell metabolic organization and reveals how the same organizing principles govern electron flow across diverse metabolic architectures.

## Introduction

Nobel laureate Albert Szent-Györgyi’s famous observation that “*life is nothing but an electron looking for a place to rest*” does not exaggerate the fundamental role of controlled electron flows in biology. Nutrient oxidation releases electrons that are relayed through intracellular redox reactions and ultimately delivered to terminal sinks, including terminal electron acceptors and reduced metabolic products, coupling carbon use to energy conservation, biosynthesis and redox homeostasis. Electron transfer is therefore a fundamental component of cellular metabolism, shaping how nutrients are oxidized, how energy is conserved and how redox balance is maintained. Individual redox reactions and carriers are well characterized, and whole-cell electron balances quantify how substrate-derived electrons are partitioned among biomass, products and terminal acceptors^1–3^. What remains unclear is whether any system-level organization of electron flow emerges from these reaction-level transfers, and if it does, how this organization relates to the whole-cell electron balance.

Existing approaches consider individual reactions, electron carriers or whole-cell electron balances, without connecting these levels into a unified metabolism-wide description of electron flow. Metabolic flux and isotope-tracing methods resolve reaction rates and atom routing^4–7^, whereas redox analyses typically focus on specific carriers such as NADH and NADPH^1,8^. While network-level descriptions treating metabolism explicitly as an electron-flow system have begun to emerge,^8,9^ whole-cell electron balances typically collapse the intracellular network to net inputs and outputs^2,3^. Here, we define the *cellular electron economy* as the system-level organization of electron generation, intracellular transfer and disposal across metabolism. Note that this differs from recent uses of the term “electron economy” to simply describe the allocation of electrons^10,11^. Testing whether the cellular electron economy follows general principles that transcend conventional pathway boundaries and recur across metabolic states and organisms requires a representation that connects wholecell electron conservation to the intracellular routes through which electrons are transferred.

We formalize the cellular electron economy through an electron-centric representation of metabolism that makes electron generation, transfer and disposal quantitatively accessible throughout the metabolic network. By combining reaction chemistry with metabolic fluxes, we derive genome-wide, reaction-resolved electron fluxes. This connects local donor–acceptor transfers to whole-cell electron flow and makes its system-level organization quantitatively testable.

We use this framework to identify the organizing principles of the cellular electron economy. Our results show that carbon uptake and electron flow can become decoupled, revealing two distinct organizing roles: within a given metabolic architecture, the carbon source sets the boundaries of the electron flow rate, whereas the disposal capacity—the capacity of the metabolic network to route electrons to available terminal sinks—determines where between the boundaries the system operates. This system-level organization recurs across microorganisms with different intracellular routing structures and predicts graded physiological responses to oxygen supply across both carbon sources and respiratory architectures. More broadly, this electron-centric view reveals a level of metabolic organization at which shared system-level principles can be separated from the organism-specific biochemical routes that realize them.

## Results

### Formalizing the electron economy

We represent metabolic states through their electron economies: where electrons are generated, how they are transferred through the network and where they are ultimately disposed of (Fig. 1a,b). By reconstructing electron flow across the metabolic network, this representation exposes a system-level organization that is obscured in conventional representations of metabolism.

**Fig. 1.**
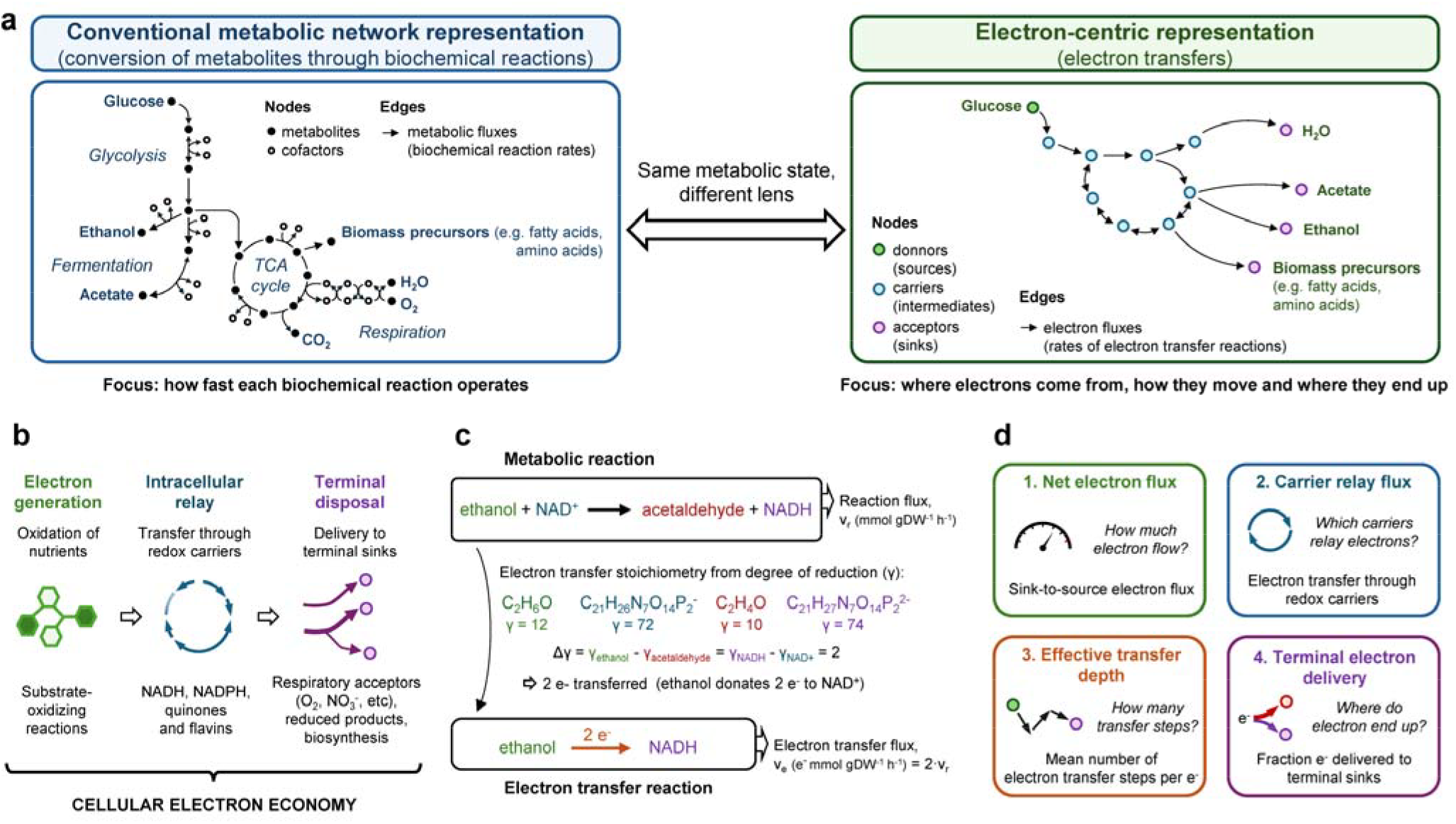
Formalizing the cellular electron economy exposes electron flow throughout metabolism. **a**, A metabolic state can be represented both by carbon fluxes (left) and in electron-transfer space (right), where redox reactions define how electrons are generated, transferred and ultimately disposed of. **b**, The cellular electron economy links three system-level processes: electron generation from nutrients, intracellular relay through redox carriers and reactions, and electron disposal to terminal sinks. **c**, Reaction chemistry converts metabolic flux into electron flux. Degree-of-reduction changes identify donor–acceptor transfers within individual reactions, e.g. in alcohol dehydrogenase, oxidation of ethanol transfers two electrons to NAD^+^. Combining this transfer stoichiometry with reaction flux yields reaction-resolved electron fluxes across the metabolic network. **d**, *Net electron flux* quantifies the magnitude of source-to-sink electron flow, and three complementary descriptors characterize its routing: *carrier relay flux* measures transfer through electron carriers, *effective transfer depth* measures the average number of steps in electron flux pathways, and *terminal electron delivery* partitions electron flow among terminal sinks.

We quantify the cellular electron economy using the degree of reduction (γ), which measures the electron equivalents associated with a compound from its elemental composition and charge^2,3^. Whereas classical electron equivalent balances are typically applied at the whole-cell scale, we apply the same principle locally to individual reactions across the metabolic network. For each reaction, changes in γ between chemically corresponding reactants and products reveal how many electrons are transferred and in which direction. We then identify the corresponding electron donors and acceptors from reaction formulae, charge and stoichiometry (Fig. 1c; Methods; Supplementary Note 1).

Combining reaction flux with electron-transfer stoichiometry yields a genome-wide map of reaction-resolved electron flux, with quantitative rates of electron transfer for each metabolic reaction (Fig. 1c). The aggregate electron economy is then characterized by distinguishing the overall magnitude of electron flows from electron flow pathways (Fig. 1d). The net electron flux (J_e_) measures the overall source-to-sink electron flow from nutrients to terminal sinks. Three complementary descriptors characterize its routing: carrier relay flux describes electron transfer through redox carriers, effective transfer depth measures the average number of transfer steps traversed by electrons, and terminal electron delivery records electron destinations. Thus, metabolic states with similar carbon uptake or overall electron flow can still differ in how electrons are relayed and where they end up.

Applied to the iML1515 metabolic reconstruction of *Escherichia coli*^12^, this framework confidently classifies 86.5% of 2,373 chemical reactions as redox or non-redox (Supplementary Tables 1–3 and Supplementary Fig. S1). The resulting electron transfer network contains 151 electron carriers and 263 directed edges, with 97% of nodes belonging to a single connected network (Supplementary Fig. S2). Thus, electron transfer defines a connected network that spans conventional pathway boundaries (Supplementary Table 4) rather than fragmenting into isolated redox subnetworks.

### Decoupling of electron and carbon flows

The aerobic-to-anaerobic transition in *E. coli* provides a familiar physiological benchmark for the proposed electron-centric representation: removing oxygen eliminates respiratory electron disposal and forces greater reliance on fermentation. We examined whether the increase in glucose uptake under anaerobiosis corresponds to greater electron flow, and how the loss of respiration changes electron routing. We analysed experimentally constrained flux distributions from aerobically and anaerobically grown *E. coli*^*13*^ and constructed the corresponding electron flux maps. As oxygen uptake decreases from 11.9 to 0 mmol gDW^−1^ h^−1^, glucose uptake increases from 8.7 to 14.9 mmol gDW^−1^ h^−1^. Meanwhile, net electron flux decreases from 68.8 to 61.3 e^−^ mmol gDW^−1^ h^−1^, corresponding to a decrease from 7.91 to 4.12 e^−^ per glucose (Fig. 2a). *E. coli* cells can thus consume more glucose while sustaining less electron flow, showing that electron flow rates are not dictated by carbon uptake rates.

**Fig. 2.**
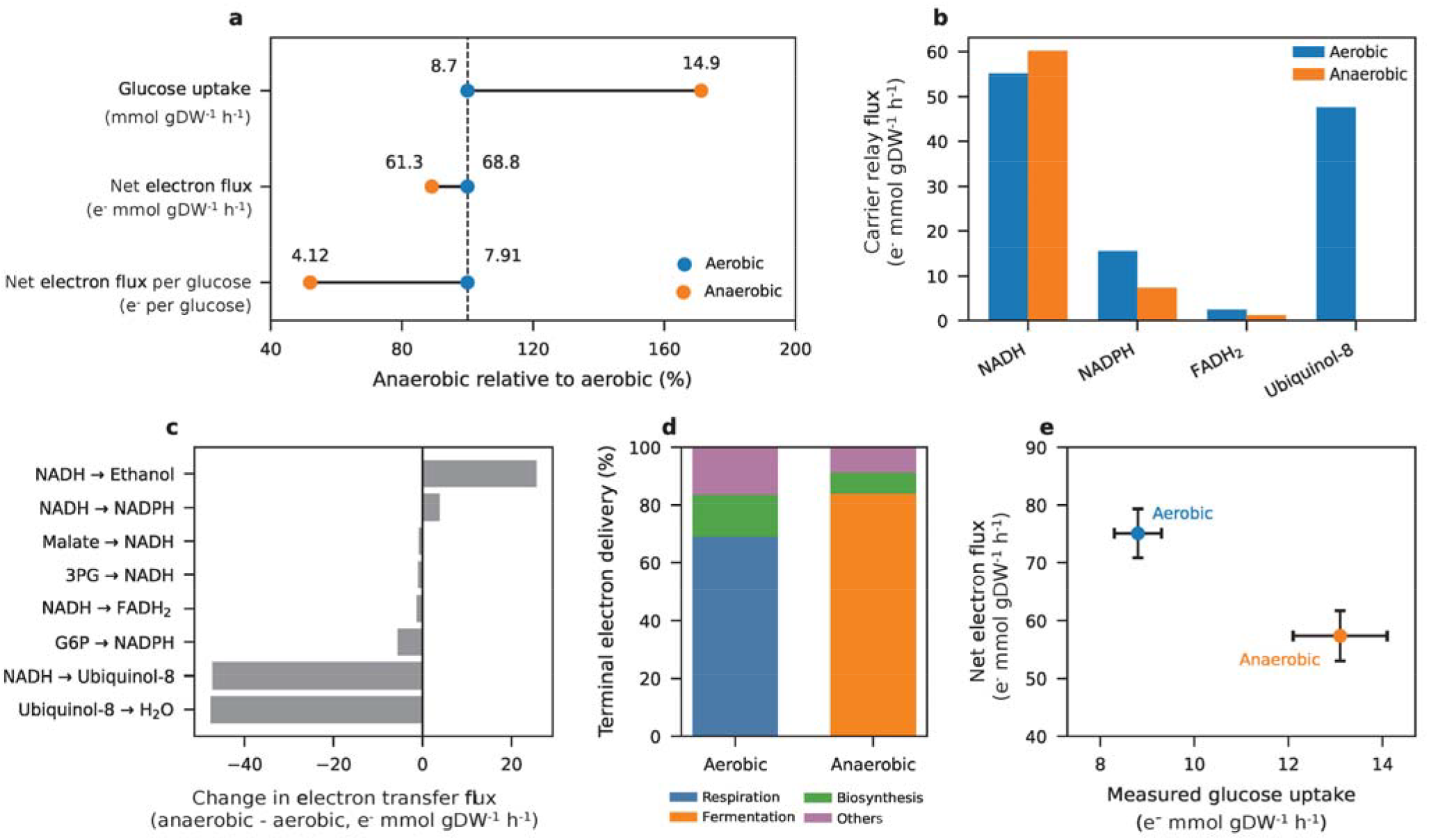
Decoupling of carbon and electron flows in glucose-grown *E. coli*. **a**, Glucose uptake increases between aerobic and anaerobic growth while net electron flux and net electron flux per glucose decrease, revealing a divergence between carbon uptake and electron flow. **b**, Relay through major redox carriers shows that this decrease is accompanied by a redistribution of carrier use. **c**, Reaction-level electron-transfer changes illustrate the extensive rerouting associated with anaerobiosis. **d**, Terminal electron delivery shifts from predominantly respiratory to fermentative sinks. **e**, Measured glucose uptake increases as net electron flux decreases. Horizontal error bars are substrate uptake rate standard deviations, and vertical error bars are conservative envelopes obtained by independent propagation of the reported marginal flux limits.

The decrease in electron flow under anaerobiosis is accompanied by extensive rerouting of electrons. NADH relay flux changes little but relay through the respiratory quinone ubiquinol-8 collapses (Fig. 2b). Respiratory electron transfer reactions disappear while NADH-to-ethanol transfer becomes a major anaerobic route (Fig. 2c). As a result, terminal electron delivery shifts from predominantly respiratory sinks (69%) to fermentative sinks (84%; Fig. 2d). This electron-centric representation thus reveals two features that carbon uptake alone does not, namely how much electron flow occurs and how that flow is routed through the network (Supplementary Fig. S3).

The electron flux maps obtained through the same approach from Gonzalez et al.’s ^13^C-MFA maps^14^ reveal the same decoupling. On glucose, from aerobic to anaerobic growth, net electron flux decreases from 8.53 to 4.38 e^−^ per glucose (Supplementary Fig. S4), while measured glucose uptake increases from 8.8 to 13.1 mmol gDW^−1^ h^−1^ and net electron flux decreases from 75.1 to 57.3 e^−^ mmol gDW^−1^ h^−1^ (Fig. 2e). The same decoupling can therefore be evidenced from experimentally measured intracellular fluxes (Supplementary Fig. S4).

We next investigated whether the lower electron flux under anaerobiosis is due to the absence of oxygen itself or to the cell’s use of oxygen for electron disposal. If the limiting factor is oxygen-linked electron disposal, the anaerobic level of electron flux should be retrievable in the presence of oxygen by blocking respiratory electron transfer. We analysed the aerobic growth of the ECOM4LA *E. coli* strain, in which terminal cytochrome oxidases are deleted and respiratory O_2_ use is nearly abolished^15^. When constrained by the measured exchange rates, the iML1515 electron transfer network yielded 10.33 e^−^ per glucose for the wild type and 4.13 for the ECOM4LA strain, closely matching the 4.10 e^−^ obtained when the respiratory capacity was set to zero (Supplementary Fig. S5). Loss of respiratory electron transfer therefore reduces electron flow even in the presence of oxygen. The electron-centric representation thus separates oxygen availability from the respiratory disposal capacity—the amount of respiratory electron transfer that the metabolic network can actually support.

As a complementary test, we varied the respiratory architecture—here, the set of terminal oxidases available to the cell—in the presence of oxygen. Across *E. coli* strains with different terminal-oxidase configurations^16^, acetate secretion showed architecture-specific responses to increasing oxygen supply and persisted despite measurable O_2_ consumption (Supplementary Fig. S6). Thus, oxygen availability alone is insufficient: respiratory architecture determines how much of oxygen’s electron-accepting capacity the cell can actually use.

### A carbon source–disposal relation

To test whether similar distinctions apply to the availability and disposal capacity of terminal acceptors other than oxygen, we investigated whether the same imposed electron-accepting capacity produces the same electron flow when supplied by different terminal acceptors. Holding glucose uptake constant, we compared O_2_, nitrate and trimethylamine N-oxide (TMAO) as terminal acceptors over a common range of imposed electron-accepting capacities. In the absence of any external respiratory terminal acceptor, the network carries 4.10 e^−^ per glucose. Increasing the acceptor capacity increases the net electron flux by an amount that depends on the acceptor (Fig. 3a; Supplementary Fig. S7). Fumarate and dimethyl sulfoxide (DMSO) likewise enhance growth and electron transfer activity but produce smaller increases in net electron flux (Supplementary Figs. S8–S9). Similarly, based on Toya et al.’s experimental ^13^C-flux data^17^, nitrate increases electron generation (the production of electron equivalents in the measured redox fluxes) from 4.27 to 6.38 e^−^ per glucose relative to strict anaerobiosis (Supplementary Fig. S10).

**Fig. 3.**
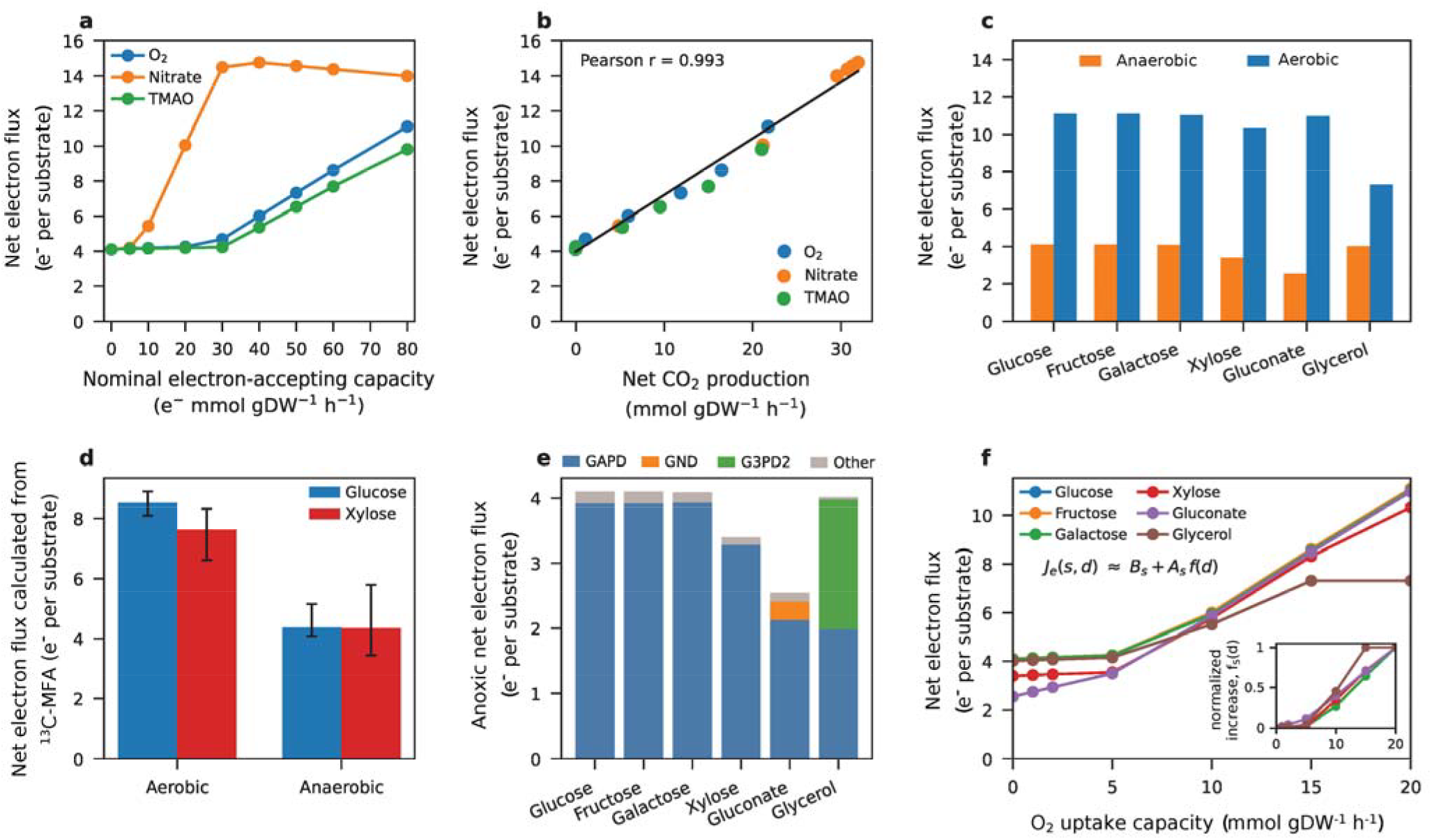
A quantitative relationship defines the separate roles of carbon source and disposal capacity in the electron economy. **a**, At fixed glucose uptake, O_2_, nitrate and TMAO support different increases in net electron flux when compared over a common range of electron-accepting capacities. **b**, The increase in net electron flux closely tracks additional carbon oxidation measured as net CO_2_ production (Pearson r=0.993). **c**, Across six carbon sources at equal uptake rates, reducing respiratory capacity to zero shifts net electron flux from the substrate-specific maximal rate towards its baseline. **d**, Electron flux maps calculated from independent ^13^C-flux data^14^ reproduce the complementary effects of carbon source and disposal capacity on glucose and xylose, where error bars are conservative marginal-limit envelopes. **e**, Reaction-level decomposition shows that the substrate-specific baseline fluxes are overwhelmingly accounted for by a small set of electron-generating reactions. **f**, Across the complete six-substrate × seven-O_2_-capacity response surface, the separate responses converge onto a common trajectory after subtracting substrate-specific baselines and scaling by the corresponding response amplitude. The inset shows the scaled curves from the empirical relationship J_e_(S,d) ≈ B_S_ + A_S_f(d).

As expected from redox stoichiometry, net electron flux closely tracks additional carbon oxidation across O_2_, nitrate and TMAO titrations (Pearson r = 0.993; Fig. 3b). The key question is therefore what determines how extensively carbon can be oxidized. What matters is not only how much electron-accepting capacity is available, but how effectively the metabolic network can use it to support carbon oxidation. We refer to this effective network-level capacity as disposal capacity.

We next investigated the dependence of electron flow on carbon source at zero and high O_2_ uptake capacity. At matched substrate uptake rates, net electron flux is 45–77% lower at zero than at high respiratory capacity across glucose, fructose, galactose, xylose, gluconate and glycerol (Fig. 3c). However, the baseline electron flux varies substantially between substrates, ranging from 2.54 e^−^ per gluconate to 4.10 e^−^ per glucose or fructose. The ~4 e^−^ value for glucose is therefore one of several substrate-specific baseline electron fluxes (Supplementary Table 5).

These complementary effects of carbon source and disposal capacity were also observed in the results of electron flux calculations based on experimental data^14^: net electron flux per substrate decreased from 8.53 to 4.38 e^−^ on glucose and from 7.63 to 4.35 e^−^ on xylose between aerobic and anaerobic growth (Fig. 3d), despite increased substrate uptake in both cases. Normalization by substrate carbon number or degree of reduction yielded similar responses for glucose and xylose (Supplementary Fig. S4). Thus, electron-flux maps calculated from independently measured intracellular fluxes reproduced this relationship, which persisted after normalization by substrate carbon number or degree of reduction.

The substrate-specific baseline arises from a small set of carbon-source-specific electron-generating reactions. Glucose, fructose, galactose and xylose catabolism is dominated by the same lower-glycolytic electron-generating reaction, catalysed by glyceraldehyde-3-phosphate dehydrogenase (GAPD), whereas gluconate and glycerol catabolism involve additional substrate-specific electron-generating reactions. Nearly all (94.9–99.1%) of the net electron flux predicted by the model is accounted for by one or two central metabolic reactions (Fig. 3e). The electron flux maps under anaerobiosis calculated from Gonzalez et al.’s flux maps^14^ likewise show GAPD accounting for 83.8% and 74.5% of net electron flux on glucose and xylose, respectively (Supplementary Tables 5-6 and Supplementary Fig. S11). The baseline electron flux thus depends on the carbon source because different substrates channel carbon through different electron-generating reactions as they enter central metabolism.

Together, these results highlight the separate roles of the carbon source and disposal capacity in the electron economy: while the carbon source sets the baseline and maximum rate of electron flow, the disposal capacity determines how close to the maximum rate the metabolism operates. To test whether these contributions are quantitatively separable, we generated 42 metabolic states by combining each of the six substrates with seven O_2_-uptake capacities. After subtracting the substrate-specific baselines and scaling the data by the corresponding response amplitude in electron flux, the six nearly overlapping response curves follow a common trajectory (Fig. 3f). This near overlap indicates that substrates differ mainly in terms of the associated baseline and maximum achievable electron flows, whereas the system’s response to increasing disposal capacity is largely shared between substrates. A rank-one decomposition, which tests how much of the response matrix can be represented by a single shared pattern, captured 99.5% of the total variance in baseline-subtracted electron-flux increase. Additive carbon-source and O_2_-capacity effects explained 95.4% of the variance, with only 4.6% attributable to the interaction. The two effects are therefore approximately separable: the carbon source determines the baseline and potential increase, whereas the response to increased disposal capacity is largely substrate-independent. Accordingly, net electron flux can be empirically approximated as:

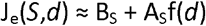

where *S* denotes the carbon source and *d* the disposal capacity. This relationship decomposes the net electron flux into a substrate-specific baseline term, B_S_, and a variable response term, A_S_f(*d*). A_S_ is the maximum response amplitude, defined as the difference between the maximal and baseline net electron fluxes for carbon source *S*. The maximal net electron flux is therefore B_S_+A_S_. f(*d*) is a normalized response function, with f(0)=0 and f(*d*_max_)=1, which describes where between the baseline and maximal rates the system operates as the disposal capacity varies. Here, *d* is parametrized by the imposed O_2_-uptake capacity. Thus, once substrate-specific differences in baseline and response amplitude are accounted for, heterogeneous metabolic states respond similarly to variations in disposal capacity. The *E. coli* electron flux response surface can therefore be characterized in terms of these two organizing variables (Supplementary Fig. S12).

### Shared principles across microorganisms

To investigate whether the organization identified in *E. coli* applies to other organisms, we applied the same chemistry-driven reconstruction, without organism-specific redox curation, to *S. cerevisiae*^18^, *B. subtilis*^19^, *S. enterica*^20^ and *K. phaffii*^21^, with 81.5–96.1% of reactions confidently classified (Supplementary Table 7).

At matched glucose uptake rates, oxygen limitation reduces the net electron flux per glucose by 48–67% across the five organisms (Fig. 4a). The oxygen-limited baseline is similar in all five, at 4.10 ± 0.09 e^−^ per glucose, whereas the response amplitude varies from 3.79 to 8.49 e^−^ per glucose (Supplementary Table 8). The intracellular routes carrying electron flow differ markedly, however. Transfer depth decreases strongly in *E. coli* and *S. enterica*, varies little in *B. subtilis*, and increases in both yeasts, whereas NADH relay flux is maintained in *E. coli* but decreases by >50% in the other four organisms (Fig. 4b and Supplementary Fig. S13). Reaction-resolved electron flux is extensively redistributed across the network in all these organisms (Supplementary Table 9). What the studied organisms share, therefore, is the system-level organization of electron flow rather than the specific pathways through which it is implemented.

**Fig. 4.**
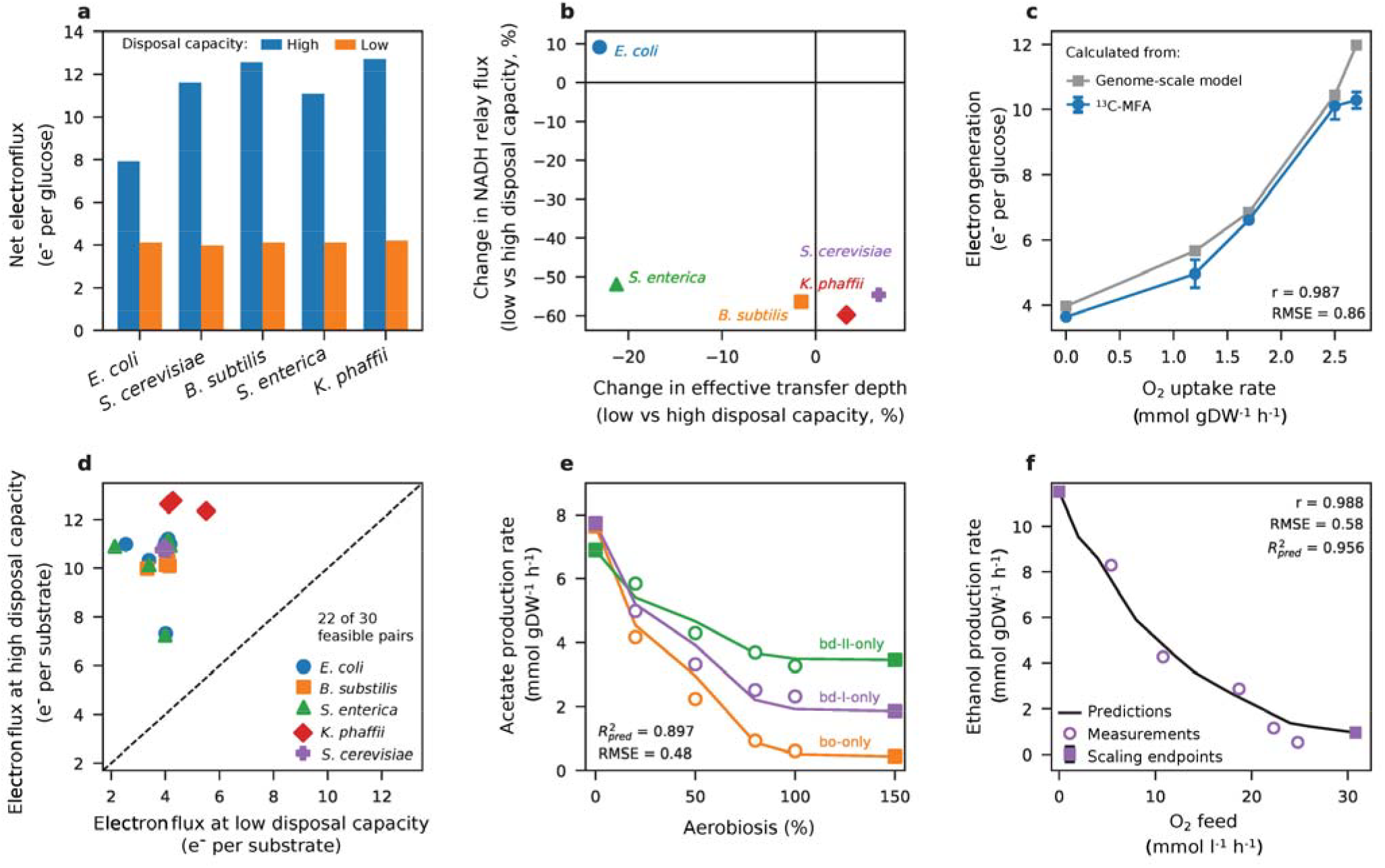
Shared organization of the cellular electron economy across microorganisms. **a**, Oxygen restriction reduces net electron flux per glucose across five microorganisms, despite differences in metabolic architecture. **b**, Accompanying changes in effective transfer depth and NADH relay differ between organisms, showing that similar system-level responses can arise through different intracellular routes. **c**, Across five oxygen conditions in *S. cerevisiae*, electron generation calculated from experimental ^13^C-flux data^22^ closely follows matched yeast-GEM predictions; data-derived points are mean ± s.d. of two cultures. **d**, Across 22 feasible organism–carbon-source pairs, higher respiratory capacity increases net electron flux above a carbon- and organism-dependent baseline. Unfeasible states are listed in Supplementary Table 10. **e**, The acetate response to aerobiosis inferred from the TBE029 *E. coli* parental background with all three terminal oxidases predicts the responses of three single-terminal-oxidase architectures^16^ after architecture-specific scaling. Filled squares denote the 0% and 150% aerobiosis states used to define each target-specific baseline and amplitude; the open circles are the 12 intermediate states used to test the predictions (predictive R^2^ = 0.897, RMSE = 0.48 mmol gDW^−1^ h^−1^; Supplementary Fig. S19). **f**, The response shape inferred from the glucose oxygen series in Weusthuis et al.’s data^25^ predicts the progressive reduction of ethanol production with increasing O_2_ supply during maltose growth. Only the zero- and highest-O_2_ maltose conditions were used to define the maltose-specific scaling; the five intermediate states were only used to test the prediction.

To test this cross-species model prediction against measured intracellular fluxes, we compared electron generation calculated from published yeast ^13^C-flux maps with the same quantity calculated from the genome-scale models. Across five oxygen levels in *S. cerevisiae*^*22*^, electron generation decreases from 10.28 to 3.64 e^−^ per glucose as oxygen availability decreases, and closely follows the matched genome-scale prediction (r = 0.987; Fig. 4c). For *K. phaffii*^*23*^, oxygen restriction decreases electron generation by 38–40% despite 79–89% increases in glucose uptake. The divergence between glucose uptake and electron generation is therefore not specific to *E. coli* (Supplementary Fig. S14).

Neither is the disposal capacity effect specific to oxygen. Across 17 feasible terminal-acceptor conditions in the five reconstructions, the increase in net electron flux correlates strongly with additional carbon oxidation (r = 0.858), but only weakly with the provided electron-accepting capacity (r = 0.350). After scaling responses within each organism, 15 conditions across the three bacteria capable of using multiple acceptors converge on a common relationship between electron-flux increase and carbon oxidation (r = 0.995; Supplementary Fig. S15). Fundamentally therefore, how much a terminal acceptor increases electron flow is related more closely to how effectively the metabolic network couples use of that acceptor to carbon oxidation than to acceptor identity or the electron-accepting capacity made available.

Does the division of labour between the carbon source and the disposal capacity persist across organisms and carbon sources? Of the 30 investigated organism–carbon-source combinations, 22 yield feasible metabolic states under both oxygen-limited and high respiratory capacity conditions, and net electron flux increases with respiratory capacity in every case (Fig. 4d). The baseline electron flux spans 2.14–5.51 e^−^ per substrate, and the response amplitude ranges from 3.24 to 8.76 e^−^ per substrate. Thus, both the baseline and maximum electron flows depend on the carbon source and the organism. Differences in substrate carbon number or degree of reduction do not generally account for this variation (Supplementary Fig. S16). This relationship also extends to acetate, pyruvate and succinate (Supplementary Fig. S17). Applying our electron-balance analysis to published whole-cell *C. shehatae* carbon-yield data^24^ highlights the same qualitative separation: the fraction of electrons delivered to O_2_ increases with oxygen availability, whereas under oxygen limitation, the nature of the carbon source determines how electrons are partitioned between reduced products (Supplementary Fig. S18).

To investigate the predictive value of this shared response structure, we first attempted to transfer the response shape across respiratory architectures with different magnitudes of acetate overflow. Using independently collected *E. coli* data covering six levels of oxygen supply^16^, we inferred the shape of the acetate response to oxygen from the parental strain after normalization between low- and high-oxygen endpoints. For each of three strains with a single terminal oxidase, the parental response predicts acetate production response across all three respiratory architectures (predictive R^2^ = 0.897, RMSE = 0.48 mmol gDW^−1^ h^−1^; Fig. 4e). Changes in the respiratory architectures therefore alter the magnitude of overflow metabolism but the transferable oxygen response shape remains the same (Supplementary Fig. S19).

We then attempted to transfer the response shape across carbon sources. If the shared response reflects the system’s disposal capacity rather than the nature of the carbon source, a response inferred on one substrate should be transferrable to another after substrate-specific scaling. In Weusthuis et al.’s chemostat data obtained with varying levels of O_2_ supply^25^ (and using ethanol production as a measure of the shift from fermentative towards oxidative metabolism) the normalized response to O_2_ supply inferred from the glucose series alone predicts the graded decrease in ethanol production with increasing O_2_ supply during maltose growth (r=0.988, predictive R^2^=0.956, RMSE = 0.58 mmol gDW^−1^ h^−1^; Fig. 4f), using only the zero- and highest-O_2_ maltose conditions as constraints. Thus, a response inferred on glucose predicts the progressive suppression of fermentative ethanol production with increasing oxygen supply on maltose (Supplementary Fig. S20).

Together, these analyses show that a system’s electron disposal capacity constrains how far electron flow can rise above a baseline set by the carbon source and metabolic architecture. This organization recurs across diverse microorganisms despite substantial differences in intracellular routing and terminal acceptors, and the shared response predicts graded physiological responses across both respiratory architectures and carbon sources. The electron-centric representation therefore separates shared system-level organization from the distinct intracellular routes and respiratory architectures that realize it.

## Discussion

An electron-centric view of metabolism reveals a simple system-level organization of electron flow that emerges from complex reaction-level behaviour. By linking whole-cell electron balance to reaction-level electron transfer, genome-scale electron-flux analysis makes this organization quantitatively testable. Because the electron-transfer network is reconstructed directly from reaction chemistry rather than from predefined pathways, the same framework can be applied across distinct metabolic architectures without tailoring the redox reconstruction to each organism. This perspective therefore separates the shared system-level structure from the various biochemical routes that instantiate it in specific organisms.

The electron-centric view also shows that electron fluxes are not dictated by the carbon uptake rate. Conventional carbon-flux descriptions emphasize how carbon is routed through metabolism, whereas the electron-centric view exposes how far that carbon is oxidized and how the associated reducing power is transferred and ultimately disposed of. This perspective provides a common explanation for apparently disparate metabolic responses: carbon uptake can increase when oxidation decreases if the electron disposal capacity is limited, and the respiratory architecture determines how extensively available terminal acceptors can support carbon oxidation. In *E. coli*, oxygen removal increases substrate uptake while reducing electron flow, and electron fluxes calculated from ^13^C-flux maps show the same decoupling. Applying the same analysis to published *K. phaffii* ^13^C-flux maps reveals a similar divergence between increased glucose uptake and reduced electron generation. Because greater carbon oxidation necessarily increases electron transfer by redox stoichiometry, the unresolved question is what determines how extensively incoming carbon can be oxidized. The electron-centric representation separates terminal-acceptor availability from the network’s ability to use that acceptor for electron disposal. Our ECOM4LA analysis shows that electron flows can remain at levels close to those calculated in the absence of respiratory capacity, even when oxygen is available. Analyses of respiratory-chain variants likewise show that oxygen availability alone does not determine the balance between oxidative and overflow metabolism. Together, these results identify disposal capacity as a system-level constraint on the level of carbon oxidation and therefore on the electron flux that the network can sustain. This reframes overflow metabolism in electron-centric terms: limited disposal capacity favours the accumulation of incompletely oxidized products, a widespread phenotype across bacteria, yeasts and mammalian cells^26^.

A second organizing principle is that carbon source and disposal capacity impose largely independent constraints on electron flow. For a given metabolic architecture, the carbon source sets the baseline and maximal electron flow rate, whereas the disposal capacity determines where within this range the system operates. Across organisms, the baseline and maximal rates also depend on the metabolic architecture and are therefore not intrinsic properties of the carbon source alone. Electron flow data across a wide range of metabolic states are well modelled by the empirical relationship J_e_(S,d) ≈ B_S_ + A_S_f(d). Furthermore, the fact that the glucose-inferred response accurately predicts the change in ethanol production with increasing oxygen supply in *S. cerevisiae* grown on maltose suggests that this response structure is predictive across carbon sources.

A third principle is that the same system-level organization can be realized through different intracellular routes. Oxygen limitation reduced net electron flux across all five studied organisms despite organism-specific changes in carrier relay and transfer depth. Consistent with this separation between organization and implementation, the oxygen-response shape inferred from the parental *E. coli* strain predicts acetate responses in three single-oxidase strains after scaling to architecture-specific endpoints. Across the studied organisms, carbon sources and terminal acceptors, increases in electron flux are more closely associated with the level of carbon oxidation than with the nature or electron-accepting capacity of the acceptor. Together, these results identify carbon source and electron disposal capacity as organizing variables applying the same constraints across various terminal acceptors and respiratory architectures. The disposal capacity reflects how effectively the metabolic network can route electrons to terminal sinks and thereby support carbon oxidation.

The concept of a cellular electron economy therefore highlights a shared level of metabolic organization. By separating the system-level organization of electron flows from their specific biochemical routes, this perspective provides a common basis for comparing metabolic states and architectures that differ extensively at the reaction level. How broadly these principles extend can now be tested across wider ranges of substrates, respiratory chemistries and biological systems, including photosynthetic, fermentative, chemolithotrophic and multicellular metabolism. Departures from the shared relationships may be especially informative, revealing where regulation, energetics, enzyme capacity or redox control impose additional constraints on electron flow. The cellular electron economy reveals not only where Szent-Györgyi’s electron comes to rest, but what determines the route and extent of its journey.

## Methods

### Metabolic models and degree of reduction

The primary model was the *E. coli* iML1515 reconstruction^12^. Yeast9/yeast-GEM v9.1.0^18^, iBB1018^19^, iYS1720^20^ and iMT1026^21^ were used for organism-specific analyses (Supplementary Note 2). Degree of reduction was calculated from model formulae and charges as γ = 4nC + nH − 2nO − 3nN + 5nP + 6nS − Q^2,3^. Compounds with undefined compositions or containing unsupported elements or transition metals were left unscored (Supplementary Notes 1 and 3).

### Chemistry-driven reconstruction of electron transfers

Reactant–product correspondence was inferred globally within each reaction from formula, charge and stoichiometry (Supplementary Note 1). Exact transport matches and 16 universal redox anchors were resolved first. Remaining occurrences were paired by maximum-cardinality, minimum-cost bipartite matching with elemental-conservation constraints. Ambiguous residual mappings were accepted only when independently supported by anchored balanced reactions, otherwise they were left explicitly ambiguous. Pairwise γ changes were used to identify donor and acceptor correspondences and reaction-level electron balance. Neither a reaction-specific blacklist nor organism-specific redox curation was used.

### Electron transfer network and electron flux quantities

Static electron-transfer topologies were based on reactions confidently classified as redox and electron-balanced. Forward flux maps a reduced donor to the reduced acceptor, and negative flux reverses the edge. Multi-pair reactions were allocated by pair-specific electron counts, while partially resolved reactions were retained in flux accounting through explicit unresolved nodes. Reaction identities were preserved in a directed multigraph. Network analyses were performed with NetworkX^27^ and, where indicated, Louvain clustering^28^, SciPy^29^ and scikit-learn^30^. Subsystem enrichment *p*-values were corrected for multiple testing using the Benjamini–Hochberg method ^31^ (Supplementary Note 4).

For reaction r with metabolic flux v_r_ and electron transfer stoichiometry n_e,r_, the magnitude of the reaction-resolved electron flux is |v_r_|n_e,r_ and direction-aware activities were assigned to donor–acceptor edges. The net electron flux, J_e_, was defined as the total positive source injection (equal to the total sink delivery within tolerance); the carrier relay at a node was calculated as min(inflow,outflow); the effective transfer depth was determined from the cumulative edge activity/J_e_; and the terminal delivery partitions sink flux between terminal fates. Fluxes are reported as e^−^ mmol gDW^−1^ h^−1^ or normalized per substrate (Supplementary Notes 1 and 5).

### *E. coli* aerobic/anaerobic analysis

Published aerobic and anaerobic *E. coli* glucose states^13^ were constrained to glucose uptake rates of 8.7 and 14.9 mmol gDW^−1^ h^−1^ and O_2_ uptake rates of 11.9 and 0 mmol gDW^−1^ h^−1^, respectively. Reported aerobic non-detected products were fixed to zero. Biomass maximization was followed by pFBA^32^ in COBRApy^33^. Experimental exchange constraints are inputs, and growth, unconstrained exchanges and internal fluxes are predictions (Supplementary Note 5 and Supplementary Table 11).

### *E. coli* carbon source and respiratory capacity perturbations

To test substrate generality across carbon sources differing in carbon number, degree of reduction and entry into central metabolism, uptake of glucose, fructose, galactose, xylose, gluconate or glycerol was fixed to 10 mmol gDW^−1^ h^−1^ and O_2_-uptake capacity was set to 0 or 20 mmol gDW^−1^ h^−1^. Percentage decreases in Fig. 3c were calculated relative to the 20 mmol gDW^−1^ h^−1^ capacity state. These endpoints are the 0 and 20 mmol gDW^−1^ h^−1^ states of the factorial response surface shown in Fig. 3f (Supplementary Note 6 and Supplementary Table 5).

For ECOM4LA, phenotypic means for oxic wild-type MG1655 and ECOM4LA were taken from Portnoy et al.^15^. iML1515 was constrained to the reported glucose, oxygen, D-lactate and acetate exchange rates, biomass was maximized and parsimonious flux balance analysis (pFBA) applied before electron flux calculation. The analysis quantifies electron flow in the experimentally observed wild-type and ECOM4LA physiological states. The multi-deletion, adaptively evolved genotype was not reconstructed reaction by reaction (Supplementary Note 7, Supplementary Tables 12–13 and Supplementary Fig. S5).

For the respiratory-architecture comparison, mean acetate secretion rates for TBE029 and the bo-only, bd-II-only and bd-I-only terminal-oxidase strains across 0, 20, 50, 80, 100 and 150% aerobiosis were taken from Steinsiek *et al*.^*16*^. These measurements served two complementary analyses: comparison of oxidative/overflow states across respiratory architectures and predictive transfer of the oxygen-response shape across architectures (Supplementary Notes 7–8, Supplementary Tables 14–16 and Supplementary Figs. S6 and S19).

### Cross-species analyses

Cross-species analyses were performed on *S. cerevisiae*/Yeast9, *B. subtilis*/iBB1018, *S. enterica*/iYS1720 and *K. phaffii*/iMT1026 with the same chemistry rules as for *E. coli*. For each organism, paired glucose states were compared at the same substrate-uptake rate. The physiological oxygen-limited endpoint for *K. phaffii* was an upper bound on O_2_ uptake of 0.5 mmol gDW^−1^ h^−1^. All six substrates in all five reconstructions were screened (30 combinations) and unavailable or infeasible states were explicitly retained, yielding 22 supported paired states. Robustness analyses included O_2_ sensitivity, targeted FVA, equal-parsimony alternatives and iJO1366^34^ (Supplementary Notes 2 and 6 and Supplementary Tables 7–8, 10 and 17–18). Network-wide electron flux redistribution was additionally quantified between paired glucose states as Jensen–Shannon and total-variation distances between normalized reaction-resolved donor–acceptor electron flux distributions (Supplementary Note 9 and Supplementary Table 9).

### Terminal acceptor and carbon source analyses

For the main acceptor titrations, glucose uptake was fixed at 10 mmol gDW^−1^ h^−1^ and O_2_, nitrate and TMAO were compared on a common imposed electron-accepting-capacity grid of 0, 5, 10, 20, 30, 40, 50, 60 and 80 e^−^ mmol gDW^−1^ h^−1^. These imposed capacities were converted to acceptor-specific uptake caps using 4 e^−^ per mmol O_2_ and 2 e^−^ per mmol nitrate or TMAO before pFBA. Fumarate and DMSO were tested at representative endpoints, with net CO_2_ extracted from the same solutions. Independent non-O_2_ constraints (obtained from Denby^35^, Toya^17^ and Perrenoud and Sauer^36^) were used for validation, with the quantities reported in each study kept distinct from genome-scale source-to-sink fluxes. Electron generation was calculated from reported ^13^C-derived redox fluxes^17^ using two electrons per NAD(P)H or FADH_2_ equivalent (Supplementary Note 10). A separate five-reconstruction capability screen was performed to evaluate 24 candidate organism–acceptor capabilities at fixed glucose, with absent and zero-uptake cases explicitly retained. For non-carbon terminal acceptor conditions supporting uptake, the resulting increase in net electron flux was related to both the imposed electron-accepting capacity and the associated increase in CO_2_ production. For the three multi-acceptor bacteria, expansion and oxidation were scaled by within-organism maxima before pooling 15 states for comparative analysis (Supplementary Note 11, Supplementary Tables 19–20 and Supplementary Fig. S15).

For carbon source analysis, glucose, fructose, xylose, galactose, gluconate and glycerol uptake rates were fixed at 10 mmol gDW^−1^ h^−1^ under the organism-specific oxygen-limited and high-respiratory-capacity conditions defined above. Dominant electron sources in the oxygen-limited state were identified directly from pFBA source decomposition (Supplementary Note 6 and Supplementary Tables 5 and 17). All five reconstructions were also attempted with acetate, pyruvate and succinate to extend entry-point coverage. Feasibility and oxygen-limited/high-respiratory-capacity growth ratios are reported in Supplementary Note 11, Supplementary Table 21 and Supplementary Fig. S17.

### Carbon source × O_2_-capacity factorial

Glucose, fructose, xylose, galactose, gluconate and glycerol were also crossed in a factorial design with seven O_2_-uptake capacities (0, 1, 2, 5, 10, 15 and 20 mmol gDW^−1^ h^−1^), yielding 42 states. For a given substrate, S, the baseline electron flow rate was calculated as B_S_ = J_e_(S,0), the response amplitude as A_S_ = J_e_(S,20) − B_S_, and the normalized increase over baseline as g_S_(d)=[J_e_(S,d)-B_S_]/A_S_. The six normalized curves overlapped closely and the empirical shared response f(d) was calculated as the mean of these curves at each tested disposal capacity, where d represents the disposal capacity as parameterized in this analysis by the imposed O_2_-uptake capacity. The level of agreement between the normalized curves was quantified by pairwise correlations. The fraction of the variance in electron flux captured by the first principal component was calculated by singular-value decomposition of the baseline-subtracted expansion matrix at non-zero O_2_ capacities. In other words, most of the apparently substrate-specific expansion could be described by a single shared response shape. A balanced two-way decomposition of the raw 6 × 7 response matrix was performed to partition variance into substrate, O_2_-capacity and interaction components; the analysis was also repeated after normalization by the substrate carbon number and by the degree of reduction. These deterministic states were analysed descriptively. The resulting relationship, J_e_(S,d) ≈ B_S_ + A_S_ f(d) therefore describes the observed response surface without imposing a functional form on f(d) (Supplementary Note 12, Supplementary Tables 22a–b and Supplementary Fig. S12).

### Electron flux calculations from experimental *E. coli* ^13^C-fluxes

We calculated electron fluxes from the ^13^C fluxes reported by Gonzalez et al.^14^ using the *E. coli* electron-transfer network, study-specific reactions and biomass definitions. NAD(P)H and FADH_2_ production and consumption were assigned two electron equivalents per reduced cofactor. Carrier conversions were treated as relay steps that did not contribute to net electron generation, whereas direct formate→CO_2_+H_2_ was assigned two electrons per turnover. The mapped states yielded balanced source-to-sink electron fluxes and were normalized by carbon number and γ using substrate stoichiometry. Because Gonzalez et al.^14^ did not report joint reaction-flux covariance data, marginal lower/upper limits were propagated independently to conservative envelopes rather than via formal joint 95% confidence intervals (Supplementary Note 13 and Supplementary Tables 6 and 23).

### Electron flux calculations from experimental yeast ^13^C-fluxes

Starting from the ^13^C-flux maps reported by Jouhten et al.^22^ for glucose-limited *S. cerevisiae* at five oxygen levels, positive NADH-, NADPH- and FADH_2_-generating contributions (two electrons per reduced cofactor) were summed and normalized by the measured glucose uptake rate. We refer to this observable as electron generation, which, unlike J_e_, does not require a balanced source-to-sink reconstruction. For Fig. 4c, yeast-GEM was constrained only by glucose and O_2_ uptakes measured at each condition, with anaerobic model adjustments at 0% O_2_. Growth was predicted and no intracellular ^13^C-flux was fitted. Electron generation was then calculated using the same procedure applied to the experimental ^13^C-flux data (Supplementary Note 14 and Supplementary Table 24a).

For *K. phaffii*, experimental ^13^C-fluxes were obtained from Baumann et al.^23^ for glucose-limited control and Fab-expressing strains at 21%, 11% and 8% inlet O_2_. Because the published flux maps did not close redox or O_2_ balances, we calculated electron generation rather than balanced source-to-sink flux. Signed NADH-, NADPH- and FADH_2_-producing stoichiometries were converted using two electron equivalents per reduced cofactor, positive source contributions were summed and normalized by measured glucose uptake, and no genome-scale FBA or pFBA solution entered this calculation. Error bars were propagated from reported marginal flux s.d. (Supplementary Note 14, Supplementary Table 24b and Supplementary Fig. S14).

### *C. shehatae* whole-cell electron equivalent analysis

Whole-cell electron retention in glucose- and xylose-grown *C. shehatae* was calculated from the carbon yields reported by Fromanger et al.^24^ using degree of reduction per carbon (4 for glucose/xylose, 6 for ethanol, 4.4 for polyols, 4.67 for glycerol and 4.34 for measured biomass). Because CO_2_ is fully oxidized, it carries no reducing equivalents. O_2_-delivery fraction (representing the proportion of reducing power balanced by oxygen uptake) was calculated as [4 − Σ_i_(Y_C_,iγ_i_)]/4. Sensitivity to the pooled pyruvate/succinate/fumarate assignment was <0.4 percentage points, and aerobic estimates agreed with independent CO_2_/RQ balances (Supplementary Note 6, Supplementary Table 25 and Supplementary Fig. S18).

### Predictive value of the response shape across respiratory architectures

Using acetate formation data reported by Steinsiek et al.^16^ for the common Δ*nuoB* parental strain TBE029, which retains all three terminal oxidases, and derived strains retaining only bo, bd-II or bd-I across six levels of aerobiosis, the initial source response shape was defined by normalizing the TBE029 acetate series between 0% and 150% aerobiosis states. The architecture-specific baselines and acetate response amplitudes for each single-oxidase target strain were then calculated using only the corresponding 0% and 150% states. The normalized TBE029 response was then rescaled between these endpoints to predict acetate formation at 20, 50, 80 and 100% aerobiosis, yielding 12 predictions. R^2^ and RMSE metrics were calculated across these intermediate states only. Robustness was assessed using 100% rather than 150% aerobiosis as the high endpoint, each respiratory architecture in turn as the source response, leave-one-architecture-out prediction, omission of individual aerobiosis levels, omission and interpolation of corresponding source-response levels, and propagation of the reported acetate uncertainties. Rank-one decomposition of the baseline-subtracted response matrix and balanced two-way variance decomposition were used to assess shared response structure independently of endpoint scaling (Supplementary Note 8, Supplementary Table 16 and Supplementary Fig. S19).

### Predictive value of the response shape across carbon sources

These calculations were based on the data reported by Weusthuis et al.^25^ for *S. cerevisiae* CBS 8066 grown in chemostats at a dilution rate of 0.10 h^−1^ on glucose or maltose with graded O_2_ feeds. Net ethanol production was used as a physiological readout of the shift from fermentative towards oxidative carbon use. These measurements support a physiological response comparison rather than the calculation of reaction-resolved electron fluxes. The medium contributed approximately 12 mM ethanol from the growth-factor stock, so net specific ethanol production was calculated as q_ethanol_ = D(C_ethanol,out_ − C_ethanol,in_)/X, where D is dilution rate and X is biomass concentration. Positive values denote net ethanol formation and negative values net consumption.

The transferable response shape was obtained from the glucose series only. The two ethanol concentrations reported as not detected (<1 mM) for the glucose cultures at the highest O_2_ feeds were set to 0 mM in the primary analysis, and net ethanol production was fitted as a decreasing function of O_2_ feed using monotonic regression (scikit-learn). The fitted glucose response was normalized between 0 and 30.8 l^−1^ h^−1^ O_2_ feed. For maltose, the baseline was set to the mean of the two zero-O_2_ measurements and the high-oxygen endpoint to the 30.8 mmol l^−1^ h^−1^ O_2_ state. The five intermediate maltose states at 5.4, 10.8, 18.7, 22.3 and 24.8 l^−1^ h^−1^ were held out from both response fitting and endpoint scaling and predicted by scaling the normalized glucose response between the maltose-specific endpoints. Sensitivity analyses performed with values between 0 and 1 mM for the censored glucose measurements confirmed that the predictive result was insensitive to variations in this parameter (Supplementary Note 15, Supplementary Table 26 and Supplementary Fig. S20).

### Statistics and reproducibility

Network enrichment analyses were performed using one-sided Fisher exact tests with Benjamini–Hochberg correction of p-values for multiple testing. Acceptor-series relationships were summarized by Pearson or Spearman correlation coefficients. Agreement between the yeast-GEM predictions and the electron generation calculated from the Jouhten et al. ^13^C-flux data was evaluated using Pearson *r*, RMSE, range-normalized RMSE and mean absolute percentage error; no intracellular ^13^C-fluxes were fitted. Deterministic model screens, factorial decompositions, normalizations and feasibility analyses were treated descriptively. For *E. coli* electron flux envelopes were constructed by propagating marginal limits independently; for *S. cerevisiae* electron generation, error bars were calculated as sample s.d. across two cultures; and for *K. phaffii* electron generation calculations from ^13^C-fluxes, error bars are the first-order propagated s.d. from marginal flux uncertainties. Extended acceptor analyses were based on pooled and within-organism-scaled correlations as defined above. For the predictive transfer of the response shape across carbon sources, Pearson r, predictive R^2^ = 1 − Σ(y − ŷ)^2^/Σ(y − ⍰)^2^ and RMSE were calculated across the five intermediate maltose states used for prediction only (the zero- and highest-O_2_ maltose endpoints used for scaling were excluded). For the predictive transfer of the response shape across respiratory architectures, R^2^ and RMSE were calculated across the 12 intermediate states, and robustness analyses were performed for the endpoint and source-architecture choice, leave-one-architecture-out prediction, individual-level omission and propagation of reported measurement uncertainty as detailed in Supplementary Note 8. Network-wide electron-flux redistributions were quantified using base-2 Jensen–Shannon distance and total-variation distance between normalized edge-flux distributions. Targeted flux variability/equal-parsimony analyses were used to assess optimum dependencies. Exact tolerances, software versions, provenance and results are provided in Supplementary Note 16, Supplementary Data and the repository.

### AI disclosure

ChatGPT (OpenAI) assistance was used for code cleanup and packaging to support reproducibility. The code was checked and approved by the authors.

## Data availability

The genome-scale models iML1515, Yeast9, iBB1018, iYS1720 and iMT1026 are provided in the repository. The Gonzalez^14^, Jouhten^22^ and Baumann^23^ publications and their original supplementary files, the Fromanger^24^ publication tables and the Weusthuis^25^ chemostat data were used as sources of the experimental flux, whole-cell carbon/redox and predictive-transfer analyses. Reaction-level tables derived from previously published data, uncertainty summaries, electron flux calculations and electron equivalent balances are provided as Supplementary Data. The source data for all quantitative panels are supplied with this manuscript. Portnoy^15^ and Steinsiek^16^ provided the experimental data used for the ECOM4LA and respiratory-architecture analyses.

## Code availability

All code used to reconstruct electron transfer networks, calculate electron fluxes, perform cross-species and perturbation analyses, calculate electron fluxes from experimental ^13^C-flux maps, perform the predictive-transfer analyses and generate figures are available at https://github.com/MetaSys-LISBP/Cellular_electron_economy.

## Acknowledgements

The authors thank Benjamin Pfeuty (Université de Lille, Lille, France), Louis Casteilla (RESTORE, Toulouse, France), Brice Enjalbert (TBI, Toulouse, France), and Mathieu Cardoso (Toulouse, France) for insightful discussions.

## Author contributions

Funding acquisition: PM, JCP; Project administration: PM, JCP; Conceptualization: PM, JCP; Investigation: PM; Methodology: PM, JCP; Software: PM; Formal Analysis: PM; Visualization: PM; Writing – original draft: PM; Writing – review & editing: PM, JCP.

## Competing interests

The authors declare no competing interests.

## Funding

This work was supported by the MITI program of CNRS (METABO-FLOW grant), the MICA department of INRAE (COCA-COLI grant), the INRAE Metaprogram DIGIT-BIO (HEPATOTWIN grant), the ANR (ASCENDING grant, ANR-21-F2II-0002), and BPIFRANCE (OPTISTEM grant, DOS0168827/00). MetaboHub-MetaToul (Metabolomics & Fluxomics facilities, Toulouse, France; https://mth-metatoul.com) is part of the French national infrastructure MetaboHUB and is funded by the ANR with grant number MetaboHUB-ANR-11-INBS-0010.

## Notes

https://github.com/MetaSys-LISBP/Cellular_electron_economy

### Competing Interest Statement

The authors have declared no competing interest.

