## Supplementary information for "Organizing principles of the cellular electron economy"

### Content

This Supplementary Information provides the technical description, validation, physiological configurations, experimental analyses, and robustness analyses supporting this study. Quantitative values are traceable to machine^-^readable files in the accompanying Supplementary Data and repository.

Supplementary Notes 1–16.

Supplementary Figs. S1–S20.

Supplementary Tables 1–26.

### Supplementary Note 1 — Chemistry-driven reconstruction of electron transfer networks

#### Chemical basis and supported domain

The electron transfer network (ETN) reconstruction is based on the degree of reduction γ, an electron-equivalent measure computed from elemental composition and formal charge. For a molecular formula containing C, H, O, N, P and S and formal charge Q, γ = 4C + H − 2O − 3N + 5P + 6S − Q. The coefficients correspond to reference oxidation states C(+4), H(+1), O(−2), N(−3), P(+5) and S(+6). Pairwise changes in γ are used only after a reactant–product molecular correspondence has been established. A non-zero Δγ then reports the number and direction of electron equivalents transferred by that correspondence.

The supported formula domain is deliberately conservative. Metabolites with missing formulae, generic placeholder composition, transition-metal chemistry or unsupported elements are retained in the model but cannot be chemically scored by the formula-based procedure. This boundary is explicit in reaction-level outputs. The method therefore does not infer electron transfer from reaction names, EC annotations or hand-written reaction-specific rules when chemical composition is insufficient.

#### Preprocessing and universal redox anchors

Reaction stoichiometry is expanded into reactant and product occurrences. Exact same-species transport between compartments is resolved first and assigned as simple translocation with zero electron transfer. Integer stoichiometric multiplicities up to eight are represented as repeated occurrences. This allows the same global matching machinery to resolve stoichiometric cases such as disproportionation using the shared matching rules.

Where applicable, 16 universal redox pairs are matched independently of compartment to establish initial reactant–product correspondences: NAD⁺/NADH, NADP⁺/NADPH, FAD/FADH₂, FMN/FMNH₂, ubiquinone-8/ubiquinol-8, menaquinone-8/menaquinol-8, 2-demethylmenaquinone-8/2-demethylmenaquinol-8, O₂/H₂O, O₂/H₂O₂, NH₄⁺/NO₂⁻, NO₂⁻/NO₃⁻, NO/NO₂⁻, oxidized/reduced thioredoxin, oxidized/reduced glutathione, oxidized/reduced ferredoxin and oxidized/reduced cytochrome c. Additional correspondences are inferred from elemental composition, charge and stoichiometry using the global matching procedure described below. For both anchored and chemically inferred correspondences, electron-transfer stoichiometries are calculated from changes in degree of reduction. The anchors therefore assist molecular matching within a chemistry-driven reconstruction that identifies redox transformations beyond the predefined pairs. In iML1515, 41.6% of retained matched redox-pair instances are inferred independently of direct anchor matching. Supplementary Note 3 reports these reconstruction statistics and an anchor-removal analysis assessing the anchors’ contribution to reconstruction coverage and molecular mapping.

#### Global reactant–product matching

After transport and anchor matching, remaining occurrences are paired globally within each reaction by maximum-cardinality, minimum-cost bipartite matching. A formula candidate must conserve C, N, P and S within the pair and must share at least one elemental constituent. H, O and charge may differ. Among eligible candidates, the implemented cost is |ΔH| + |ΔO| + 0.1|ΔQ|. Maximum cardinality is optimized before total cost. This global formulation prevents local greedy choices from consuming a chemically preferable correspondence needed elsewhere in the same reaction.

For each accepted pair, Δγ = γ_product_ − γ_reactant_. A negative Δγ identifies an electron-donating correspondence in the written reaction direction, whereas a positive Δγ identifies an electron-accepting correspondence. The resolved donated and accepted electron totals are calculated independently. Reactions are classified as fully resolved when both sides are present and equal within tolerance, partially resolved when both are present but unequal, donor-side-only or acceptor-side-only when only one side is chemically identified. The point estimate of electron-transfer magnitude is max(resolved donation, resolved acceptance). For incomplete states this is a lower bound. Minimum unresolved donation or acceptance is recorded explicitly as the smallest additional contribution needed to balance the identified side.

#### Ambiguity detection and independent support logic

Formula and charge identify molecular correspondence uniquely for the chemically resolved subset of multi-substrate reactions. For balanced formula-only residual problems containing at most seven occurrences on each side, the implementation enumerates competing assignments to determine whether the minimum-cost redox-looking assignment has an equally plausible group-transfer alternative. An internal support set is inferred only from balanced reactions that are independently anchored by the universal redox cofactors. This support is applied in a single non-recursive step. An otherwise ambiguous formula-only mapping is accepted only when at least one inferred redox pair has independent support. Otherwise the reaction is returned as explicitly ambiguous. The method uses one chemistry-based rule set across all reactions.

#### Electron transfer edges and direction-aware electron flux calculation

The static electron-transfer network contains only confidently redox, fully resolved, electron-balanced reactions. Nodes represent reduced (i.e. electron-bearing) species. For a donor correspondence D_red_→D_ox_ and acceptor correspondence A_ox_→A_red_, forward flux generates the edge D_red_→A_red_. Reverse physiological flux reverses that electron-transfer direction. Multi-pair reactions are represented by multiple reaction-labelled edges, with pair-specific electron stoichiometry preserving the total electron number in the reaction. Partially and one-sided resolved reactions are excluded from the fully resolved static topology but remain in flux accounting through reaction-specific unresolved nodes, so incomplete chemistry is not silently discarded.

#### Algorithm box

For each model:
1. Parse metabolite formula and charge. Compute γ where chemically supported.
2. For each non-structural reaction, expand integer stoichiometric occurrences (≤8).
3. Match exact same-species transport occurrences.
4. Match the 16 universal redox-anchor pairs, independent of compartment.
5. Match remaining occurrences by maximum-cardinality/minimum-cost bipartite matching:
candidate ⇔ C/N/P/S conserved and elemental overlap present
cost = |ΔH| + |ΔO| + 0.1|ΔQ|.
6. For small balanced formula-only residuals (≤7 occurrences/side), test competing assignments.
7. Build an independent support set from balanced seed-anchored reactions only.
8. Accept an ambiguous formula-only assignment only if at least one inferred redox pair has independent support. Otherwise label ambiguous.
9. Compute pair Δγ, resolved donated/accepted electrons and resolution status.
10. Build the fully resolved static ETN. Retain incomplete reactions through unresolved nodes for flux accounting.
11. Calculate direction-aware electron fluxes from a metabolic flux vector using reaction-specific electron-transfer stoichiometries, then calculate system-level observables.

This pseudocode mirrors the implemented order in etn/matching.py, etn/reaction_electron_balance.py and etn/electron_flow_analysis.py. Thresholds used here are MAX_EXPLICIT_STOICH_OCCURRENCES = 8, MAX_AMBIGUITY_ENUMERATION = 7 and a numerical matching/electron-balance tolerance of 10^−9^ unless a downstream analysis specifies a larger activity threshold.

### Supplementary Note 2 — Cross-species reconstruction and response to oxygen restriction

**Rationale and model selection.** To test whether the organizing principles of the cellular electron economy generalize beyond *E. coli*, the same chemistry-driven reconstruction was applied to independent genome-scale models of *Saccharomyces cerevisiae*^18^, *Bacillus subtilis*^19^, *Salmonella enterica*^20^ and *Komagataella phaffii*^21^ (*Pichia pastoris*), alongside the primary *E. coli*^12^ reconstruction. The panel spans bacterial and compartmentalized yeast metabolism, allowing shared electron-flux organization to be tested across different cellular and respiratory architectures. The core degree-of-reduction calculation, reactant–product matching, ambiguity detection and electron balancing were not modified by organism-specific redox curation. Only input-format adapters, compartment parsing and physiological exchange configurations were model specific. Exact bundled files and SHA256 hashes are given in Supplementary Table 7 and data/cross_species/ in the repository.

**Technical transferability.** Across the five reconstructions, 81.5–96.1% of chemically analysable reactions were confidently classified as redox or non-redox, and 70.2–82.2% of identified redox reactions were fully resolved on both donor and acceptor sides. The resulting ETNs contained 105–222 electron-bearing metabolites, 112–198 distinct biochemical redox couples, 117–240 contributing reactions and 157–287 directed edges. The largest weakly connected component contained 60.8–100% of electron-bearing metabolites. This range reflects genuine differences in model size, chemical annotation and compartmentalization. Each organism retains the chemical annotation and reconstruction coverage of its source model.

**Physiological configurations.** The experimentally constrained *E. coli* states are those used in the main analysis. Yeast used the Yeast9 v9.1.0 model-provided anaerobic configuration, including its anaerobic biomass and exchange edits. *B. subtilis* and *S. enterica* were analysed at fixed glucose uptake with nitrate uptake blocked so that oxygen withdrawal tested fermentative anoxia rather than alternative nitrate respiration. *K. phaffii* was analyzed over an oxygen-uptake titration. Because this yeast is strongly respiration dependent, the reported low-oxygen endpoint is severe oxygen limitation (0.5 mmol gDW^-1^ h^-1^) rather than mathematical zero oxygen.

**Electron flux compression.** Net source-to-sink electron flux normalized to glucose uptake decreased in every paired organism: from 7.91 to 4.12 e^-^ per glucose in *E. coli*, 11.60 to 3.97 in *S. cerevisiae*, 12.55 to 4.10 in *B. subtilis*, 11.07 to 4.11 in *S. enterica*, and 12.70 to 4.21 in *K. phaffii* under severe oxygen limitation (Supplementary Table 8). The five low-oxygen endpoints therefore fall in a narrow 3.97–4.21 e^-^ per glucose interval (mean 4.10 ± 0.09, s.d., coefficient of variation 2.1%). Cumulative electron-transfer activity per glucose decreases by 60–71% across the same paired comparisons. These are model-defined physiological states except for the experimentally imposed *E. coli* exchange constraints.

**Carbon source attribution.** In the anoxic *E. coli*, *S. enterica* and *B. subtilis* solutions, the GAPD reaction accounts for approximately 95–97% of net electron-source flux and contributes about 3.9–4.0 e^-^ per glucose. In anaerobic *S. cerevisiae*, GAPD together with the two oxidative pentose-phosphate dehydrogenase steps accounts for approximately 94% of net source flux. These calculations support the interpretation that removing respiratory electron-accepting capacity suppresses additional oxidative electron-generation routes, leaving a low-flux regime dominated by the glucose catabolism backbone. The remaining electron-source contributions retain organism- and condition-specific routes.

**Routing remains organism specific.** Carrier usage and transfer depth remain organism specific despite the convergence of baseline electron fluxes. Effective transfer depth decreases strongly in *E. coli* and *S. enterica*, changes little in *B. subtilis*, and increases slightly in both yeasts. NADH relay is maintained in *E. coli* (55.1 to 60.1 mmol e^-^ gDW^-1^ h^-1^) but falls substantially in *S. cerevisiae* (78.0 to 35.4 mmol e^-^ gDW^-1^ h^-1^), *B. subtilis* (92.0 to 40.1 mmol e^-^ gDW^-1^ h^-1^), *S. enterica* (83.6 to 40.2 mmol e^-^ gDW^-1^ h^-1^) and *K. phaffii* (99.4 to 39.9 mmol e^-^ gDW^-1^ h^-1^) (main-text Fig. 4b, Supplementary Fig. S13). Thus, the robust cross-species signature is the compression of electron flux, whereas the remaining flux is routed through organism-specific intracellular routes.

#### Model and flux robustness

**Oxygen supply sensitivity and pFBA non-uniqueness.** The main cross-species conclusions were evaluated across alternative optimal pFBA solutions. System-level observables remain stable across these solutions, whereas pathway-level assignments are reported as model-resolved routes for the selected solution. In a targeted robustness analysis of three representative organisms, targeted flux variability analysis (FVA) showed that high-activity individual reaction assignments are least constrained in aerobic yeast, whereas most high-activity reactions are sign-invariant in anaerobic yeast and *B. subtilis*. Equal-parsimony alternative optima are more informative for the global ETN quantities reported here: in aerobic yeast the tested alternatives span only 1.60% in net e^-^ per glucose, 0.129% in effective transfer depth and 2.38% in NADH relay. Anaerobic yeast and the tested *E. coli* and *B. subtilis* alternatives are essentially invariant at the reported precision. The oxygen-titration analyses documented in the accompanying Source Data further show that the electron-flux response is graded with oxygen availability rather than dependent on a single arbitrary binary flux solution.

**Independent *E. coli* reconstruction.** To test whether the compression is peculiar to iML1515, the same standardized glucose/O_2_ comparison was repeated with iJO136634. With glucose uptake fixed to 10 mmol gDW^-1^ h^-1^ and an aerobic oxygen uptake upper bound of 20 mmol gDW^-1^ h^-1^, iML1515 gives 11.110 e^-^ per glucose aerobically and 4.103 anaerobically, whereas iJO1366 gives 10.749 and 4.158 e^-^ per glucose, respectively. Thus, both independently reconstructed *E. coli* models reproduce a ~61–63% decrease and the ~4.1 e^-^ per glucose anoxic endpoint.

### Supplementary Note 3 — Extensive chemical validation and reconstruction quality control

#### Degree-of-reduction validation

**Method.** The degree of reduction γ = 4C + H - 2O - 3N + 5P + 6S - Q was evaluated for phosphate, the adenylate series (ATP, ADP and AMP), and eight additional reference redox/non-redox pairs using the formula and charge encoded for each compound in iML1515.

**Result.** Two internal consistency properties are recovered exactly: γ(phosphate)=0, because phosphate is the phosphorus reference state, and γ(ATP)=γ(ADP)=γ(AMP)=30, so adenylate interconversion by phosphate transfer carries no change in degree of reduction. Across ten pairwise comparisons, ATP/ADP and ADP/AMP are not eligible formula-based correspondences because phosphorus stoichiometry differs, whereas the eight additional reference pairs are either seed- or formula-matched. All canonical redox couples yield the expected electron count, while glucose 6-phosphate/fructose 6-phosphate is formula-matched with Δγ=0 and is therefore correctly identified as non-redox.

#### 52-reaction categorical validation

**Method.** A 52-reaction set was assembled to sample the major categories of redox and non-redox chemistry encountered in the reconstruction: isomerases, kinases, a hydratase, transport reactions, phosphate group transfer, NAD(P)-dependent dehydrogenases, flavoprotein reductases, quinone- and menaquinone-linked dehydrogenases, transhydrogenases, an amino-acid oxidase and ribonucleotide reduction. Four reactions were included specifically to test handling of chemistry outside the method’s scorable domain: three genuine redox reactions involving Fe(II)/Fe(III) or a thioredoxin cofactor encoded with a generic ‘X’ formula, and one non-redox ferrochelatase reaction involving an unscorable metal. The set was hand-curated by the authors and is therefore a categorical validation set rather than an independent blinded benchmark. Confusion-matrix metrics were calculated on the 48 chemically scorable reactions. The four unscorable reactions were evaluated separately for whether they were correctly flagged as outside the method’s domain.

**Result.** All 48 scorable reactions were classified correctly (TP=35, TN=13, FP=0, FN=0, with accuracy=precision=recall=specificity=F1=1.00), and all four unscorable reactions were flagged as such rather than assigned a redox status (Supplementary Table 1).

#### Genome-scale model classification coverage

**Method.** Each of the 2,712 reactions in iML1515 was assigned to one mutually exclusive outcome: confidently redox, confidently non-redox, ambiguous formula-only mapping, chemically unscorable because of unsupported elemental chemistry or generic formulae, or structurally excluded (biomass, exchange, demand or sink). Coverage is reported relative to all reactions and to the 2,373 chemically analysable reactions remaining after 339 structural exclusions.

**Result.** The method confidently classifies 2,053 reactions, corresponding to 75.7% of all reactions and 86.5% of chemically analysable reactions. Of these, 270 are redox and 1,783 non-redox. Six reactions are retained as ambiguous rather than forced. Generic formula groups are the largest single source of missing chemical coverage (160 reactions), followed by transition-metal chemistry (127 reactions, Supplementary Table 2 and Supplementary Fig. S1).

#### Matching specificity and explicit ambiguity

**Method.** Formula and charge identify molecular correspondence uniquely for the chemically resolved subset of multi-substrate reactions. The algorithm therefore performs global maximum-cardinality/minimum-cost bipartite matching on reactant/product occurrences, after same-species transport and universal seed-pair matching. Formula candidates conserve C/N/P/S, require elemental overlap and use the cost |ΔH|+|ΔO|+0.1|ΔQ|. Integer stoichiometric coefficients up to eight are expanded into explicit molecular occurrences. For balanced formula-only residual problems containing at most seven occurrences per side, alternative assignments are enumerated to identify an equally stoichiometrically plausible group-transfer correspondence. An internal redox-pair support set is inferred only from balanced reactions independently anchored by universal seed cofactors. A balanced ambiguous formula-only mapping is accepted only when at least one inferred redox couple has this independent support, otherwise it is reported as ambiguous. This strategy applies one chemistry-based rule set across all reactions.

**Result.** Only six iML1515 reactions remain ambiguous from formula and charge alone: NDPK1, NDPK8, VALTA, ACOXT, DHAPT and CRNBTCT (Supplementary Table 3). They are not counted as redox or non-redox in the confident classification. Conversely, difficult genuine redox cases including coupled inorganic chemistry and disproportionation are retained when the global stoichiometric/formula evidence is sufficient. The explicit ambiguous state is therefore a methodological boundary rather than a manually curated correction.

The matching procedure enables specific assignments with a shared chemistry-based rule set, but formula and charge define two distinct resolution boundaries. First, some balanced multi-substrate reactions admit alternative correspondences that remain ambiguous from elemental composition alone, six such reactions are labelled ambiguous. Second, a reaction may be confidently redox because a cofactor couple is resolved while the corresponding organic substrate-product transformation changes C, N, P or S through coupled group transfer. Glyceraldehyde-3-phosphate dehydrogenase and pyruvate dehydrogenase illustrate this second case: both are quantified as 2-electron NAD^+^/NADH reactions, while the organic counterpart remains unresolved.

The static electron-transfer graph contains fully resolved, electron-balanced reactions. Reaction-level and flux-weighted analyses represent any unresolved molecular partner explicitly. Nodes represent reduced electron-bearing species: for a donor correspondence D_red_→D_ox_ and an acceptor correspondence A_ox_→A_red_, the forward edge is D_red_→A_red_ and the chemically reversed edge is A_red_→D_red_. Partially or one-sided resolved reactions remain in the reaction-level and flux-weighted analyses, where the missing side is represented explicitly. Constitutional and stereochemical isomers require structural information or atom mapping beyond formula and charge. These cases remain outside the current confidence framework.

#### Reconstruction and network quality control

Across iML1515, 505 matched redox-pair instances are retained: 295 arise from direct universal-anchor matching and 210 (41.6%) from formula/stoichiometry-based inference. The latter 210 reactions are molecular correspondences inferred ab initio independently of seed matching. The fully resolved electron-transfer network contains 151 electron carriers and 263 directed simple edges. 147 nodes lie in the largest weakly connected component (WCC). The reconstructed network is shown in Supplementary Fig. S2.

#### Contribution of redox anchors

To assess the contribution of redox anchors, the iML1515 reconstruction was repeated with all 16 predefined anchor pairs removed. This also eliminated the support inferred from anchored reactions, while chemical matching rules and ambiguity criteria were unchanged. Without anchors, the procedure identified 230 of the 270 redox reactions recovered by the standard reconstruction (85.2%). Of the remaining 40 reactions, 38 became ambiguous and two were classified as non-redox. Including the anchors increased the number of fully resolved, electron-balanced redox reactions from 172 to 219 and reduced ambiguous classifications from 44 to 6. Among the 230 retained redox identifications, 229 retained the same electron-transfer estimate. Catalase changed from a fully resolved 2-electron transfer to an acceptor-side-only 4-electron estimate. These results show that chemistry alone recovers most redox identifications, while the generic anchors improve molecular correspondence, ambiguity resolution and reconstruction completeness.

### Supplementary Note 4 — Network characterization and complementary topology analyses

Topology and community analyses characterize the static fully resolved *E. coli* electron transfer network and test how its organization compares with conventional metabolite connectivity and how sensitive the resulting graph properties are to construction choices.

The complete set of 161 distinct model-level inferred redox-pair IDs is provided in Supplementary Data (results/ab_initio_redox_pairs.csv). After collapsing compartment duplicates and reverse orientations, these correspond to 150 biochemical pair identities. Pair order follows the reaction-direction orientation of the inferred correspondence whereas condition-specific electron-flux direction is reported in the flux-weighted outputs. Here “inferred” identifies the algorithmic origin of a pair found independently of direct seed matching.

For comparison with conventional metabolite connectivity, an undirected metabolite co-occurrence graph is constructed from all non-biomass, non-exchange, non-demand and non-sink reactions after excluding 18 compartment-specific bookkeeping species: h_c, h_p, h_e, h2o_c, h2o_p, h2o_e, atp_c, adp_c, amp_c, pi_c, pi_p, ppi_c, co2_c, co2_p, nh4_c, o2_c, o2_p and coa_c. Redox carriers such as NAD(H), NADP(H), flavins and quinones are retained. For the electron-transfer graph, the directed reaction-resolved network is converted to an undirected graph by counting each distinct reaction once per unordered metabolite pair: opposite edges generated by a reversible reaction are not double-counted, whereas distinct reactions connecting the same pair are additive. Both graphs are then induced on their 149 common metabolites before direct comparison. The induced electron graph contains 190 undirected edges and the metabolic graph 268. All 190 electron edges are contained in the metabolic graph, giving a Jaccard similarity of 0.7090. Weighted-degree Spearman correlation is ρ=0.577 (P=1.27×10^-14^). At random_state=0 the induced electron graph contains twelve Louvain communities and the metabolic graph nine, with ARI=0.494 and NMI=0.532. Across ten seeds the electron partition is invariant, whereas the metabolic graph varies between nine and ten communities.

Community robustness, community-resolution sensitivity and conventional-network comparison sensitivity were calculated with analyse_community_robustness.py, analyse_community_resolution_sensitivity.py and analyse_network_comparison_sensitivity.py. The corresponding canonical tables are provided in Supplementary Data.

### Supplementary Note 5 — *E. coli* electron economy under aerobic and anaerobic conditions

This note documents the physiological constraints used to convert reaction-level electron-transfer assignments into condition-specific electron fluxes. To generate electron flux maps for wild-type *E. coli* K-12 MG1655 grown on glucose under matched aerobic and anaerobic conditions, we first used the experimental data reported by Chen et al.^13^ to define the model constraints. These data included intracellular ^13^C metabolic flux analysis (^13^C-MFA) measurements and extracellular flux measurements. Specifically, glucose uptake was 8.7 and 14.9 mmol gDW^-1^ h^-1^, and oxygen uptake was 11.9 and 0 mmol gDW^-1^ h^-1^ under aerobic and anaerobic conditions, respectively, where gDW denotes gram dry weight. Chen et al. also reported no detectable lactate, succinate, formate, or ethanol production under aerobic conditions. Accordingly, the corresponding exchange fluxes were constrained to zero only in the aerobic state. The consistency of these experimentally derived constraints with the metabolic model was independently demonstrated in the parsimonious flux balance analysis (pFBA) reported by Jamialahmadi et al.^37^, which reproduced the same glucose and oxygen uptake constraints and validated the corresponding flux solution. We therefore used these experimentally supported constraints to generate electron flux maps. Importantly, the glucose and oxygen uptake rates and the aerobic product non-detections originate from experimental measurements reported by Chen et al. and were used as model constraints. By contrast, predicted growth, unconstrained product exchange rates and intracellular reaction fluxes are model outputs rather than ^13^C-MFA measurements.

A reaction’s matched redox pairs are defined relative to the written reaction direction, whereas physiological electron-flux direction is determined by the sign of the pFBA flux. For a donor correspondence D_red_→D_ox_ and an acceptor correspondence A_ox_→A_red_, positive flux generates the reduced-pool edge D_red_→A_red_, whereas negative flux generates A_red_→D_red_. For example, ALCD2x is written as ethanol + NAD^+^ ↔ acetaldehyde + NADH. Chemically, ethanol donates electrons to NAD^+^. After accepting them, NAD^+^ is converted to NADH. Because the electron-transfer network follows the molecular species carrying those electrons immediately before and after the reaction, the forward edge is therefore ethanol→NADH, while reverse flux gives NADH→ethanol.

Cumulative electron-transfer activity sums fluxes for every active electron-transfer flux and therefore counts the same electron again when it is relayed through successive carriers. To obtain a complementary net-flux measure, the directional pFBA-weighted electron graph is analyzed as a conserved flow layer separately for each condition. Before aggregation, the generic unresolved-organic partner is replaced by a reaction-specific node (for example, unresolved:RXN_ID), which keeps unresolved chemistry from unrelated reactions separate. Parallel electron-transfer contributions are then summed. For each node *i*, inflow_i_ and outflow_i_ are calculated and b_i_ = outflow_i_ − inflow_i_ defines net electron injection (b_i_>0), net terminal removal (b_i_<0), or balanced relay. Net source-to-sink electron flux is J_e_ = Σ_i_ max(b_i_,0), which equals total sink delivery within numerical tolerance. Carrier relay flux is min(inflow_i_,outflow_i_). Effective transfer depth is cumulative edge activity divided by J_e_, providing a decomposition-independent measure of the number of electron-transfer events per net source-to-sink electron equivalent. Raw terminal sinks are classified as respiration, fermentation, biosynthesis/other or unresolved according to the reduced sink species. For the physiological interpretation in main-text Fig. 2d, only the reaction-specific unresolved sink unresolved:ACALD is reassigned to a fermentative reporting category because ACALD is the first 2-electron reduction in the acetyl-CoA-to-ethanol branch. Reaction matching, electron accounting and network construction remain unchanged by this reporting-layer reassignment.

Net source-to-sink electron flux is 68.832 mmol e^-^ gDW^-1^ h^-1^ aerobically and 61.330 anaerobically, despite the increase in measured glucose uptake from 8.7 to 14.9 mmol gDW^-1^ h^-1^. Normalized to glucose uptake, net electron flux falls from 7.912 to 4.116 electron equivalents per glucose, while effective transfer depth decreases from 2.754 to 2.118 transfers per net electron equivalent. Raw terminal delivery is 69.2% respiratory aerobically and 0% respiratory anaerobically. In the anaerobic state, resolved fermentation sinks account for 42.0% of terminal delivery and unresolved sinks for 50.4%. Unresolved:ACALD contributes 25.598 mmol e^-^ gDW^-1^ h^-1^ ^1^ and, when this first reduction of the ethanol branch is grouped with resolved fermentation sinks for interpretation, fermentative delivery is 83.8% and other unresolved delivery 8.7%. NADH relay flux remains high (55.081 to 60.122), whereas ubiquinol-8 relay falls from 47.599 to zero. Cumulative electron-transfer activity decreases from 189.568 to 129.921 mmol e^-^ gDW^-1^ h^-1^, or from 21.79 to 8.72 electrons per glucose.

### Supplementary Note 6 — Carbon source dependence and baseline electron-generating backbones

**Carbon sources.** Glucose, fructose, xylose, galactose, gluconate and glycerol span differences in carbon number, degree of reduction and entry into central metabolism. Paired aerobic and anaerobic electron flux maps were calculated for these substrates at fixed substrate uptake of 10 mmol gDW^-1^ h^-1^. Net electron flux decreases under anaerobiosis for all six substrates, but the low-oxygen endpoint varies from 2.545 e^-^ per gluconate to 4.103 e^-^ per glucose or fructose. A minimal set of electron-generating reactions accounts for 94.9–99.1% of the model-defined anaerobic net electron flux: GAPD for glucose, fructose, xylose and galactose, GAPD and GND for gluconate, and G3PD2 and GAPD for glycerol. These reactions therefore predict the dominant core contribution of the compressed model state rather than a rigid physiological endpoint.

All six substrates were supplied at the same uptake rate (10 mmol gDW^-1^ h^-1^) in aerobic and anaerobic states. The anaerobic net source-to-sink baseline ranges from 2.54 e^-^ per gluconate to 4.10 e^-^ per glucose or fructose. Glucose, fructose, xylose and galactose are dominated by GAPD. Gluconate additionally uses GND, and glycerol uses G3PD2 plus GAPD. The selected dominant reactions account for 94.9–99.1% of net anaerobic source-to-sink flux. These reactions were identified from each computed electron flux map before calculating their contribution and were not chosen to force the approximately 4 e^-^ hexose result.

The ~4 e^-^ per glucose state should therefore not be interpreted as an acceptor-imposed constant. It emerges from the dominant electron-generating reactions that remain active when extensive oxidation is curtailed. Disposal capacity permits oxidative expansion above this substrate-dependent baseline, while carbon source chemistry determines the baseline itself.

#### Cross-species test of carbon source and acceptor-dependent expansion

**Systematic cross-species design.** To test the two-part organization across the complete candidate set, we considered all six core carbon sources (glucose, fructose, galactose, xylose, gluconate and glycerol) in all five reconstructions, giving 30 candidate organism–substrate combinations. Substrate uptake was fixed to 10 mmol gDW^-1^ h^-1^. Alternative respiratory acceptors were closed so that the imposed O_2_-uptake capacity parameterized respiratory disposal capacity. To minimize this capacity, oxygen uptake was set to zero in *E. coli*, *B. subtilis* and *S. enterica*, 0.5 mmol gDW^-1^ h^-1^ in *K. phaffii*, and the model-provided anaerobic biomass/nutrient configuration in *S. cerevisiae*. Aerobic conditions used an O_2_ uptake upper bound of 20 mmol gDW^-1^ h^-1^. Canonical substrate exchanges absent from a reconstruction were recorded as unavailable, and states without a positive growth optimum were retained as infeasible rather than omitted.

Across 22 feasible organism–substrate pairs spanning all five organisms, baseline electron flux ranged from 2.14 to 5.51 e^-^ per substrate. Increasing the imposed O_2_-uptake capacity increased net source-to-sink electron flux in every feasible pair, by 3.24–8.76 e^-^ per substrate. Carbon source also changed baseline electron flux within several organisms, whereas the three feasible *S. cerevisiae* hexoses clustered tightly. Thus a carbon-dependent baseline electron-flux regime and expansion with disposal capacity recur across microorganisms, while the quantitative baseline and routing remain organism specific. The complete 30-combination feasibility matrix is retained in Supplementary Table 10.

**Normalization robustness.** To determine whether carbon-source-dependent baseline electron flux simply reflected carbon number or total substrate reducing-equivalent content, each feasible baseline was additionally normalized by substrate carbon number and substrate γ. Substantial within-organism variation persisted in *E. coli* (0.424–1.339 e^-^ per C, 0.116–0.287 of γ), *S. enterica* (0.356–1.333 e^-^ per C, 0.097–0.286 of γ) and *K. phaffii* (0.701–1.103 e^-^ per C, 0.175–0.276 of γ). In *B. subtilis*, the four feasible substrates collapsed to 0.667–0.686 e^-^ per C and 0.167–0.172 of γ, while the three feasible *S. cerevisiae* hexoses collapsed still more tightly (0.661–0.664 e^-^ per C, 0.165–0.166 of γ). Substrate stoichiometry therefore explains much of the variation in some organisms but not in others. Expansion with disposal capacity remains positive for all 22 feasible pairs under both normalization because each pair is scaled by the same substrate-specific denominator.

***S. cerevisiae.*** The systematic screen used the model-provided anaerobic biomass/nutrient configuration for every anaerobic yeast condition so that the physiological treatment was consistent with the validated glucose comparison. Glucose, fructose and galactose supported paired states. Xylose and glycerol supported aerobic growth but not the configured anaerobic condition, and canonical gluconate uptake was absent from the reconstruction. These outcomes are reported rather than filtered.

**Interpretation.** The cross-species extension supports the organizing principle across organism-specific mechanisms and quantitative response magnitudes. The recurring feature is the separation between a baseline electron-flux regime whose scale reflects substrate chemistry and the organism, and electron-flux expansion enabled by disposal capacity. Carbon-number and substrate-γ normalization show that the baseline variation is not simply a consequence of substrate size in most tested organisms, while the *B. subtilis* collapse provides an explicit counterexample in which substrate reducing-equivalent content explains most of the observed carbon-source difference. Detailed electron-generating reactions, routing responses and relationships to CO_2_ production remain organism specific.

#### Independent yeast whole-cell validation of carbon source effects and electron disposal

Fromanger et al.^24^ provide an experimental endpoint-level test of both carbon source and oxygen availability in the xylose-fermenting yeast *Candida shehatae*. Cells were grown separately on glucose or xylose, with three controlled oxygen-limited phases for each carbon source. The study reports mean specific oxygen uptake rates (OUR), carbon yields into biomass and extracellular reduced products, and aerobic reference cultures. Carbon and degree-of-reduction balances were reported to close within 10% during oxygen limitation and within 6% aerobically. These data therefore permit whole-cell electron-equivalent accounting independent of the intracellular flux calculations used elsewhere in this study.

**Electron-equivalent calculation.** Glucose and xylose both have a degree of reduction of 4 e^-^ per carbon. For each condition, the retained reducing equivalents were calculated as the sum of each reported carbon yield (Cmol product per Cmol substrate) multiplied by the product degree of reduction per carbon. The fraction delivered to O_2_ was then estimated as [4 − Σ(Y_i_γ_i_)]/4. We used γ/C = 6 for ethanol, 4.4 for xylitol, arabitol and ribitol, 4.67 for glycerol, and 4.34 for biomass from the experimentally measured biomass formula C_1_H_2.08_O_0.66_N_0.14_ reported by Fromanger et al. CO_2_ has γ = 0. The pooled organic-acid yield contains pyruvate, succinate and fumarate. The polyol term is the reported sum of xylitol, arabitol and ribitol. Because individual proportions were not reported, the primary calculation uses γ/C = 3.25, with a sensitivity bracket of 3.0–3.5. This choice changes inferred O_2_ delivery by <0.4 percentage points in every condition.

Across the oxygen-limited series, the inferred fraction of substrate electron equivalents delivered to O_2_ increased from 2.02% to 15.91% on glucose and from 7.08% to 23.39% on xylose. Aerobic product/biomass balances were 32.70% on glucose and 36.06% on xylose (Supplementary Table 25 and Supplementary Fig. S18). As an independent check, the aerobic CO_2_ yields and respiratory quotients imply O_2_-electron fractions of 33.39% and 34.87%, respectively, within 1.2 percentage points of the product/biomass degree-of-reduction estimates (Supplementary Fig. S18). Thus, increasing acceptor availability shifts terminal electron disposal toward O_2_ on both carbon sources, while the reported ethanol/polyol partition remains strongly substrate dependent.

**Interpretation.** The result strengthens the joint organizing role of carbon entry and electron disposal in a phylogenetically distant yeast using whole-cell endpoint accounting: acceptor availability controls the fraction of substrate reducing power disposed through oxygen on both glucose and xylose, whereas carbon source influences the routing of electrons under oxygen limitation. The original study reports whole-cell carbon and redox balances, which support endpoint-level electron-equivalent partition.

### Supplementary Note 7 — Respiratory electron disposal and respiratory-architecture controls

**Rationale.** The main text uses ECOM4LA and the Steinsiek linear-respiratory-chain strains to distinguish ambient oxygen from the respiratory disposal capacity that is actually realized by the respiratory architecture and metabolic state. This note documents the experimental inputs, state-constrained electron-flux calculation, uncertainty/robustness controls and independent respiratory-chain datasets. Reported physiology, isotope-derived oxidative data and genome-scale electron-flux quantities calculated by the framework are kept explicitly distinct.

**ECOM4LA.** Portnoy et al.^15^ compared oxic wild-type *E. coli* MG1655 with ECOM4LA, an evolved strain lacking the major terminal cytochrome oxidase systems together with *ygiN*. The measured phenotype provides a deliberately extreme perturbation in which oxygen remains present while oxygen utilization is nearly abolished. The state-constrained analysis fixes the reported glucose, oxygen, D-lactate and acetate rates in iML1515, maximizes growth and applies pFBA before calculating electron fluxes from the resulting balanced metabolic flux state using reaction-specific electron-transfer stoichiometries. This calculation completes the measured physiological state in the genome-scale model.

**Endpoint electron-delivery calculation from reported measurements and uncertainty.** A model-independent endpoint observable was calculated from measured O_2_ uptake and D-lactate secretion as 4qO_2_ + 2q_lactate_, normalized by glucose uptake. The published rate standard deviations were propagated by Monte-Carlo sampling. This endpoint calculation is a sink-side experimental constraint and is not equated with balanced genome-scale net source-to-sink electron flux.

**Robustness.** Allowing secretion of otherwise unreported ethanol, formate and succinate changes the WT calculated electron flux only minimally while preserving the ECOM4LA endpoint. The main mechanistic inference is stable when secretion of the unreported products is allowed. The complete edge-, sink- and path-level outputs and the state-constrained analysis script are provided in the repository under data/experimental/portnoy2010_ecom4la/.

**Steinsiek respiratory-chain series.** Steinsiek et al.^16^ constructed strains with different terminal oxidase architectures and quantified by-product formation across 0, 20, 50, 80, 100 and 150% aerobiosis. The production of the by-product acetate, i.e. the range of acetate overflow, progressively diverge as oxygen supply increases: the *bo*-only architecture nearly suppresses acetate production at high aerobiosis, whereas bd-II-only and bd-I-only retain substantial overflow. The analysis quantifies the architecture × oxygen interaction using acetate formation as the physiological readout.

**Anand et al. synthetic unbranched electron-transport variants^38^.** Four synthetic electron-transport architectures span the published aerobicity order 1H < 3H < 2H < 4H. We calculated the fraction of glucose carbon secreted as acetate from the replicate extracellular rates. The inverse ordering of acetate overflow is present before evolution and persists after adaptive evolution even though evolved growth rates converge. The published intracellular flux maps provide model-derived flux states.

### Supplementary Note 8 — Predictive transfer of the oxygen-response structure across respiratory architectures

#### Experimental dataset and physiological observable

Steinsiek et al.^16^ reported mean acetate formation across 0, 20, 50, 80, 100 and 150% aerobiosis for the common Δ*nuoB* parental strain TBE029, which retains all three terminal oxidases, and for strains retaining only bo, bd-II or bd-I. These are published byproduct measurements and provide an architecture-resolved physiological readout. Acetate formation provides the measured physiological readout for the transfer test.

#### Predictive transfer analysis

The TBE029 acetate series was normalized between its 0% and 150% aerobiosis endpoints to define the source response shape. For each single-oxidase target, only its 0% and 150% states set the target-specific baseline and amplitude. The normalized parental response was then rescaled to predict acetate formation at 20, 50, 80 and 100% aerobiosis. Those 12 intermediate values were held out from both response-shape inference and endpoint scaling.

Across the 12 test conditions, predictive R^2^ was 0.897 and RMSE was 0.48 mmol gDW^-1^ h^-1^. A baseline^-^subtracted rank-one decomposition captured 99.78% of response variance. Balanced two-way decomposition attributed 83.92% of variance to aerobiosis, 10.67% to architecture and 5.42% to their interaction.

#### Robustness and uncertainty boundary

Using 100% rather than 150% aerobiosis as the high endpoint gave predictive R^2^ = 0.826 and RMSE = 0.61 mmol gDW^-1^ h^-1^. Changing the source architecture gave predictive R^2^ values from 0.795 to 0.933. Leave-one-architecture-out transfer gave predictive R^2^ = 0.936 and RMSE = 0.39. Omitting one parental intermediate level at a time gave predictive R^2^ values from 0.760 to 0.931. The supplied Steinsiek input contains means but no pointwise standard deviations or replicate-level observations. Accordingly, the analysis reports point estimates together with source-choice and omission sensitivity tests.

### Supplementary Note 9 — Whole-network routing-redistribution robustness

**Purpose and definition.** Main Fig. 4b uses change in effective transfer depth together with NADH relay because these directional, interpretable observables separate organism-specific routing responses despite a common compression of net electron flux. To test whether this conclusion depends on the chosen observables, we additionally compared the complete reaction-resolved donor–acceptor electron-flux distributions between paired aerobic and low-respiratory-capacity glucose conditions in all five organisms. For each state, absolute direction-aware donor–acceptor edge fluxes were normalized to unit total activity over the union of edges present in either condition. Redistribution was quantified by base-2 Jensen–Shannon distance and total-variation distance.

**Whole-network redistribution is extensive but less discriminating.** Jensen–Shannon distance was high in every organism (0.772–0.844) and total-variation distance ranged from 0.721 to 0.840 (Supplementary Table 9). Thus oxygen restriction causes substantial reaction-level rerouting in all five organisms, corroborating the interpretation of Fig. 4b. However, the narrow cross-organism range of the whole-network distances provides less directional discrimination among organism-specific responses than effective transfer depth and NADH relay: for example, transfer depth decreases strongly in *E. coli* and *S. enterica* but increases in the yeasts. We therefore use effective transfer depth and NADH relay in the main figure to distinguish the direction and nature of routing changes, and retain whole-network redistribution as a complementary robustness analysis.

### Supplementary Note 10 — Terminal electron acceptors

**Rationale.** The cross-species glucose analyses identify recurrent compression of net electron flux when oxygen availability is removed or strongly restricted. To determine whether this response is specific to oxygen, and whether the approximately 4 e^-^ per glucose low-flux regime simply reflects glycolytic NADH stoichiometry, we challenged the *E. coli* iML1515 model by varying terminal electron acceptors and carbon substrates. O_2_, nitrate, TMAO, DMSO and fumarate were selected to span established aerobic and anaerobic respiratory pathways in *E. coli*. This panel encompasses oxygen respiration, reduction of inorganic oxyanion (nitrate), organic N- and S-oxides (TMAO and DMSO), and a dicarboxylate (fumarate). These pathways differ in terminal respiratory enzymes and quinone usage, providing biologically distinct routes for electron disposal. All analyses used the same electron-transfer reconstruction and electron-flux calculation rules as in the main text.

**Alternative terminal electron acceptors.** Glucose uptake was fixed to 10 mmol gDW^-1^ h^-1^. O_2_, nitrate and trimethylamine N-oxide (TMAO) uptake capacities were titrated independently, with growth maximized and pFBA applied at each condition. At zero terminal acceptor, net source-to-sink electron flux is 4.103 e^-^ per glucose. Increasing O_2_ to 20 mmol gDW^-1^ h^-1^ raises net electron flux to 11.110 e^-^ per glucose. Setting nitrate uptake to 10 mmol gDW^-1^ h^-1^ raises it to 10.055 e^-^ per glucose. High-capacity TMAO uptake raises it to 14.920 e^-^ per glucose. At lower TMAO capacity, and at representative dimethyl sulfoxide (DMSO) and fumarate capacities, cumulative electron-transfer activity can increase substantially while net source-to-sink electron flux remains close to the compressed state (Supplementary Fig. S8). Thus disposal capacity can increase electron relay without necessarily increasing net electron generation to the same extent.

**Carbon oxidation.** Across the complete O_2_, nitrate and TMAO titration series (27 model states; nine imposed capacities per acceptor), net electron flux closely tracks net CO_2_ production (Pearson r=0.9933, Spearman ρ=0.9474). This supports the interpretation that terminal electron acceptors increase net electron flux when their use permits additional carbon oxidation. The corresponding main-text analysis is shown in Fig. 3a,b. The correlation coefficients are descriptive summaries of structured model titrations rather than inferential evidence from independent biological observations.

#### Independent non-O_2_ acceptor datasets

**Rationale.** The model analyses above predict that disposal capacity can expand net electron flux above a carbon-source-dependent baseline electron-flux regime, but the nitrate/TMAO/DMSO perturbations in that analysis are model-derived. We therefore reanalyzed three independent published *E. coli* datasets that probe non-O_2_ terminal acceptors. The datasets provide complementary observables and are not treated as interchangeable: Denby et al.^35^ directly constrain terminal electron delivery during TMAO respiration, Toya et al.^17^ provide ^13^C-constrained central carbon fluxes from which electron generation can be calculated, and Perrenoud & Sauer^36^ provide an acceptor-specific nitrate-versus-DMSO routing contrast.

**TMAO respiration — direct terminal-delivery test.** Denby et al.^35^ studied anaerobic glucose-limited chemostats of *E. coli* K-12 MG1655 (20 mM feed glucose, dilution rate 0.2 h^-1^) and perturbed an established fermentative steady state with TMAO. At the final TMAO-respiring steady state, 45.8 ± 1.4 mM TMA was measured. Normalization to feed glucose gives 2.29 ± 0.07 TMA per glucose and, because TMAO→TMA accepts two electrons per turnover, 4.58 ± 0.14 e^-^ per glucose of directly constrained terminal electron delivery. This conversion uses only the measured TMA and reaction stoichiometry. It is calculated directly from measured TMA and reaction stoichiometry.

**Independent source-side accounting and matched model state.** The reducing-equivalent accounting reported by Denby et al.^35^ corresponds to 5.44 e^-^ per glucose, 0.80 e^-^ per glucose consumed by the explicitly represented succinate/ethanol sinks, and 4.64 e^-^ per glucose assigned to TMAO reduction. Separately, we imposed the experimentally observed TMAO/glucose ratio on iML1515 by fixing glucose uptake to 10 mmol gDW^-1^ h^-1^ and allowing TMAO uptake to 22.9 mmol gDW^-1^ h^-1^. The resulting electron flux map predicts net source-to-sink electron flux to increase from 4.103 e^-^ per glucose in fermentation to 6.043 e^-^ per glucose in the TMAO-matched state (+47.3%), with CO_2_ production of 8.499 mmol gDW^-1^ h^-1^. The 6.043 e^-^ per glucose value is a model output at an experimentally matched acceptor/substrate ratio. It must not be interpreted as an experimentally measured whole-cell electron flux. Likewise, agreement of the model TMA sink with 4.58 e^-^ per glucose is constrained by the imposed TMAO uptake and is not an independent prediction.

**Physiological redistribution.** Experimentally, acetate increases from 13.7 to 24.5 mM, whereas formate, ethanol and succinate decrease from 13.6 to 6.7 mM, 10.9 to 2.3 mM and 3.9 to 1.5 mM, respectively. The matched iML1515 state reproduces all four directions of change, although it overstates the suppression of ethanol and succinate. This comparison is therefore used as directional physiological support rather than as a quantitative validation of secretion magnitudes.

**Electron accounting.** Toya et al. quantified carbon fluxes in *E. coli* BW25113 grown in continuous cultures at a dilution rate of 0.2 h^-1^. Supplementary Table SII provides marginal flux limits for the WT anaerobic state (oxygen absent), WT nitrate respiration and Δ*arcA* nitrate respiration, normalized to glucose uptake = 100. We transcribed the redox-generating reactions GAPD, G6PDH, GND, PDH, ICDH, AKGDH, SDH and MDH and converted positive oxidative flux to two electron equivalents per turnover. Negative SDH flux in the anaerobic WT state represents fumarate reduction and is therefore an electron sink rather than an electron source. The resulting quantity is termed electron generation because it is calculated from the reactions resolved in the published central carbon model.

**Nitrate-dependent expansion.** WT electron generation increases from 4.270 e^-^ per glucose under anaerobic conditions to 6.384 e^-^ per glucose under nitrate respiration (+49.5%). Independent propagation of the reported marginal lower and upper flux limits gives conservative envelopes of 4.166–4.358 and 5.738–7.034 e^-^ per glucose, respectively. These envelopes are not joint confidence intervals. Nevertheless, even the nitrate lower bound exceeds the anaerobic upper bound by 31.7%. The glycolytic contribution is nearly unchanged (3.472→3.438 e^-^ per glucose), whereas oxidative-PPP, PDH and oxidative-TCA contributions increase from 0.518→1.898, 0.202→0.324 and 0.078→0.724 e^-^ per glucose. The explicitly represented fermentative electron sinks decrease from 2.798 to 0.008 e^-^ per glucose. Thus, the electron fluxes calculated from the ^13^C-flux data support expansion through additional oxidation rather than through increased electron generation from glycolysis alone.

**Network-topology and flux-state dependence.** Under nitrate respiration, electron generation in Δ*arcA* increases further to 8.458 e^-^ per glucose (+32.5% relative to WT nitrate), predominantly through an oxidative-TCA contribution of 3.894 e^-^ per glucose. This comparison is used as a mechanistic refinement: terminal-acceptor availability permits an oxidative electron-flux regime, whereas regulatory and network state influences how much of that accessible capacity is realized.

**Alternative electron acceptors do not consistently increase tricarboxylic acid-cycle flux.** Perrenoud & Sauer reported that the reference *E. coli* strain retains a bifurcated, non-cyclic TCA configuration under strict anoxia and in the presence of 40 mM DMSO, whereas 40 mM nitrate produces a low but significant respiratory cyclic TCA flux of approximately 0.2 mmol gDW^-1^ h^-1^. This provides an experimentally grounded counterexample to a simplistic “acceptor present = oxidative expansion” interpretation.

**Standardized model comparison.** To compare acceptor-dependent oxidative routing qualitatively under standardized model constraints, we compared iML1515 states at a common acceptor uptake upper bound of 10 mmol gDW^-1^ h^-1^. Net source-to-sink electron flux is 4.103 e^-^ per glucose without an acceptor, 10.055 with nitrate and 4.192 with DMSO. The original publication reports nitrate and DMSO medium concentrations, whereas the standardized model comparison applies common uptake upper bounds and evaluates the qualitative acceptor-specific ordering. In both experiment and standardized model analysis, nitrate supports substantially greater oxidative routing than DMSO under the tested conditions.

**Synthesis.** These datasets provide independent checks on distinct predictions. Applying the framework to reported TMAO-respiration observables supports substantial terminal electron delivery to a non-O_2_ acceptor during strict anoxia. Applying it to published nitrate flux maps supports increased electron generation through additional oxidative pathways while leaving glycolytic generation nearly unchanged. The DMSO-versus-nitrate comparison indicates that terminal-acceptor presence alone is insufficient to predict identical oxidative routing. Together, these analyses support a permissive interpretation in which terminal electron-accepting capacity enables oxidative expansion above a carbon-source-dependent baseline regime, while acceptor identity and metabolic/regulatory state determine how much of that accessible oxidative regime is realized.

**Primary-source provenance.** Denby KJ et al., Environmental Microbiology 17, 2477–2491 (2015), DOI 10.1111/1462-2920.12726. Toya Y et al., Molecular BioSystems 8, 2593–2604 (2012), DOI 10.1039/C2MB25069A. Perrenoud A & Sauer U, Journal of Bacteriology 187, 3171–3179 (2005), DOI 10.1128/JB.187.9.3171-3179.2005. Provenance notes, derived result tables and automated regression tests are bundled in the repository under data/experimental/ and results/publication/.

### Supplementary Note 11 — Extended coverage of terminal-acceptor chemistry and central carbon-source entry points

**Purpose.** The additional coverage screens test whether the organizing relationship depends on the respiratory pathways and carbon-entry points examined in the main analyses. We therefore performed a systematic model-encoded terminal-acceptor screen across all five reconstructions and an independent carbon-source screen extending to acetate, pyruvate and succinate.

**Extended terminal-acceptor capability screen.** At fixed glucose uptake, 24 model-encoded acceptor capabilities were tested across the five reconstructions and 21 produced feasible states with actual uptake. Seventeen of these were obtained with non-carbon electron acceptors. Nitrite was functional in *E. coli*, *B. subtilis* and *S. enterica*. *S. enterica* additionally supported tetrathionate and thiosulfate. Fumarate was retained as a capability state but excluded from the non-carbon correlation because uptake can also supply carbon. Across the 17 states with non-carbon acceptors, electron-flux expansion correlated strongly with additional carbon oxidation (Pearson r = 0.858, Spearman ρ = 0.848) but only weakly with nominal acceptor capacity (Pearson r = 0.350, Spearman ρ = 0.117). Within each multi-acceptor bacterium, expansion versus additional carbon oxidation gave r = 0.994 for *B. subtilis*, 0.9998 for *E. coli* and 0.997 for *S. enterica*. After scaling expansion and oxidation by their within-organism maxima, the 15 states from these three organisms fell onto a common relationship (Pearson r = 0.995, Spearman ρ = 0.957, Supplementary Fig. S15). This supports disposal capacity, rather than acceptor identity or nominal electron stoichiometry, as the relevant system-level determinant.

**Extended carbon-source coverage.** Acetate, pyruvate and succinate were tested across all five reconstructions to extend entry-point coverage from the core sugars/glycerol/gluconate set to acetyl-CoA, the pyruvate node and the TCA cycle. Thirteen of 15 organism–substrate combinations supported aerobic growth. In contrast, acetate and succinate did not support growth when respiratory capacity was minimized in any model, and the only condition where (small) growth was observed on pyruvate under minimal respiratory capacity was observed for *E. coli* (0.00499 h^-1^ versus 0.31394 h^-1^ aerobically, 1.59%). These substrates extend the test to entry through acetyl-CoA, the pyruvate node and the TCA cycle, where growth is predominantly respiration dependent. The scarcity of growth-supporting states at minimal respiratory capacity identifies conditions in which a paired baseline–expansion comparison cannot be established.

### Supplementary Note 12 — Formal carbon-entry × disposal capacity response surface

**Factorial design.** To formalize the organizing principle, electron flux maps were calculated considering glucose, fructose, galactose, xylose, gluconate and glycerol, each supplied to iML1515 at 10 mmol gDW^-1^ h^-1^, while the O_2_ uptake upper bound (the maximum allowed oxygen-exchange flux, not the realized uptake) was set independently to 0, 1, 2, 5, 10, 15 or 20 mmol gDW^-1^ h^-1^. Other tested respiratory acceptors were closed in the model. The same precomputed chemistry-driven *E. coli* electron-transfer reconstruction was used for all 42 states. The complete state table is distributed as results/publication/factorial_carbon_oxygen_states.csv and Source Data for main Fig. 3f.

**Separability analysis.** For each substrate, the zero-O_2_ baseline was subtracted and the remaining response was scaled by the full expansion at the highest tested O_2_ uptake upper bound. The six scaled response curves are highly concordant (minimum pairwise r = 0.956, mean r = 0.986), and the first singular component captures 99.51% of expansion variance. A balanced two-way decomposition of the raw matrix attributes 4.63% of total variance to carbon × O_2_ capacity interaction. This falls to 1.36% after normalization per substrate carbon and 2.18% after normalization by substrate degree of reduction. The empirical relation J_e_(s,d) ≈ B_s_ + A_s_ f(d) is therefore a compact description of the tested response surface rather than a new conservation law.

**Five-organism context.** The paired glucose comparison in main Fig. 4a contains all five modelled organisms. The systematic multi-substrate cross-species analysis in main Fig. 4d attempts all 30 combinations of six carbon sources with the five reconstructions and retains 22 feasible organism–substrate pairs with both low- and high-disposal capacity conditions. *S. cerevisiae* contributes glucose, fructose and galactose paired states using the model-provided anaerobic biomass/nutrient configuration. Unavailable or infeasible combinations are retained explicitly in Supplementary Table 10.

### Supplementary Note 13 — Validation using independent *E. coli* ^13^C-MFA flux maps

**Experimental flux maps.** Gonzalez, Long and Antoniewicz^14^ quantified intracellular fluxes for aerobic glucose, aerobic xylose, anaerobic glucose and anaerobic xylose growth of *E. coli* using parallel isotope tracers and integrated ^13^C-MFA. Net metabolic fluxes and marginal 95% confidence limits were extracted directly from the supplementary data. The four condition-specific ^13^C-MFA models and biomass reactions were used to identify redox-carrier stoichiometry. The four intracellular flux maps were taken from the published ^13^C-MFA results. Electron fluxes were calculated from those published metabolic fluxes using reaction-specific electron-transfer stoichiometries.

**Electron accounting.** The ^13^C-MFA model explicitly represents reduced nicotinamide and flavin cofactor equivalents (NADH, NADPH and FADH_2_). For each fitted net reaction flux, reduced-cofactor production and consumption were converted to two electron equivalents per NADH, NADPH or FADH_2_. Carrier-to-carrier conversions were treated as relay rather than new electron generation by using the difference between produced and consumed reduced-cofactor equivalents for source or sink assignment. Direct formate→CO_2_+H_2_ was counted as a two-electron transfer per turnover. Condition-specific biomass NAD(P)H stoichiometry was included exactly as supplied.

**Internal validation.** We verified that the parser reproduces every NADH/FADH_2_ and NADPH value in the published Gonzalez cofactor-balance table to a maximum absolute discrepancy of 1.4×10^-14^ in normalized flux units. Source and sink electron fluxes balance to better than 1.2×10^-5^ e^-^ per substrate in all four conditions. Calculated aerobic respiratory delivery corresponds to oxygen uptake of approximately 13.9 and 13.2 mmol gDW^-1^ h^-1^ for glucose and xylose, respectively, matching the authors’ reported 13.8 and 13.2 estimates.

**Electron flux compression from experimental data.** Net electron flux decreases from 8.534 to 4.376 e^-^ per glucose and from 7.634 to 4.347 e^-^ per xylose. Cumulative activity decreases from 18.322 to 10.153 and from 16.238 to 8.885 e^-^ per substrate. Measured substrate uptake increases anaerobically (glucose 8.8→13.1, xylose 9.5→10.8 mmol gDW^-1^ h^-1^), whereas absolute net electron flux decreases (glucose 75.10→57.33 mmol e^-^ gDW^-1^ h^-1^, xylose 72.52→46.95 mmol e^-^ gDW^-1^ h^-1^). Thus the decoupling between carbon uptake and electron flux is present in independent experimentally inferred intracellular flux states.

**Substrate-size and reducing-power normalization.** Glucose and xylose differ in carbon number and total reducing-equivalent content (C6 and γ=24 for glucose, C5 and γ=20 for xylose). We therefore repeated the comparison of electron flows after normalizing net electron flux by substrate carbon number and by substrate degree of reduction. The decrease in net electron-flux during the transition from aerobiosis to anaerobiosis was 0.693 e^-^ per carbon for glucose and 0.657 e^-^ per carbon for xylose, corresponding to 0.173 and 0.164 of substrate γ, respectively. Thus, anaerobic compression is preserved after controlling for substrate size and intrinsic reducing power. The anaerobic glucose and xylose envelopes still overlap after normalization. These data support an acceptor-dependent component for both carbon sources and overlapping anaerobic baselines.

**Uncertainty.** Gonzalez et al. report marginal reaction-flux limits, which support conservative envelope propagation for the derived electron fluxes. We therefore propagated each reaction’s reported marginal 95% lower and upper flux limits independently to produce conservative envelopes. These envelopes do not overlap between aerobic and anaerobic states for either substrate, and represent conservative 95% marginal-limit envelopes.

**Anaerobic source decomposition.** GAPD (see Supplementary Table 1) contributes 3.666 e^-^ per glucose (83.8% of net source flux) and 3.237 e^-^ per xylose (74.5%). The contribution of GAPD on xylose closely matches the ~3.285 e^-^ per xylose dominant-core value predicted by the pFBA carbon source analysis. However, the experimental flux map also contains formate→H_2_ (0.361 e^-^/xylose), PDH (0.339), NADP-dependent malic enzyme (0.308) and smaller TCA/biosynthetic sources, which raise total net flux to 4.347 e^-^ per xylose. The data therefore support a dominant carbon-source-dependent backbone, with electron flux modulated by auxiliary physiology rather than determined solely by the substrate.

### Supplementary Note 14 — Validation for yeast under various oxygen levels

**Experimental flux maps.** Jouhten et al.^22^ quantified central carbon flux distributions in glucose-limited *Saccharomyces cerevisiae* CEN.PK113-1A chemostats at a fixed dilution rate of 0.1 h^-1^ and five inlet-oxygen levels (20.9%, 2.8%, 1.0%, 0.5% and 0%) using ^13^C-MFA. The published Figure 2 reports two independent cultivations at each oxygen condition, with fluxes normalized to glucose uptake = 100 and replicate-specific glucose-uptake rates. The companion repository bundles the original Figure 2 and published stoichiometric model together with checksums and the directly transcribed reaction fluxes used below.

**Electron accounting.** Reduced-equivalent generation was calculated from seven reaction-level NADH/NADPH/FADH_2_-generating fluxes in the published central carbon model, using two electron equivalents per reduced nicotinamide or flavin cofactor equivalent. The 2-oxoglutarate-to-oxaloacetate mitochondrial branch was oriented in the oxidative direction, generating two NADH and one FADH_2_ equivalent (6 e^-^ per turnover), although the compact stoichiometric-model text writes the reduced cofactors on the reactant side. For each replicate, positive electron-source contributions were summed and normalized to glucose uptake. The resulting quantity is electron generation across the reactions resolved by the published central carbon model.

**Experimental oxygen-gradient response and matched prediction.** Electron generation decreases from 10.46 and 10.10 e^-^ per glucose in the two 20.9% O_2_ cultivations to 3.64 and 3.64 e^-^ per glucose under 0% O_2_. Condition means are 10.28, 10.10, 6.61, 4.96 and 3.64 e^-^ per glucose at 20.9%, 2.8%, 1.0%, 0.5% and 0% inlet O_2_, respectively. To compare like-for-like, yeast-GEM was constrained only by the measured mean glucose uptake and published mean O_2_ uptake at each condition. No intracellular ^13^C flux was used as a model constraint. Model-predicted electron generation was 11.98, 10.44, 6.84, 5.66 and 3.97 e^-^ per glucose across the same series (Supplementary Table 24a). Across all five conditions, prediction and experiment were strongly concordant (Pearson r = 0.987, RMSE = 0.855 e^-^ per glucose, range^-^normalized RMSE = 0.129, mean absolute percentage error = 9.31%).

**Uncertainty and interpretation.** Experimental error bars are sample standard deviations across the two independent cultivations at each oxygen condition (n = 2), not propagated reaction-flux confidence intervals. The matched model comparison uses the same electron generation observable on both sides and therefore provides a direct quantitative validation of the graded response. Growth was predicted rather than fitted, and the model was not calibrated against the intracellular ^13^C fluxes. The net source-to-sink flux remains a distinct observable and is not substituted for the experimental value.

The experimental observable in this note is the electron generation from the reported central carbon reactions. The matched model series is calculated using the same definition of electron generation, enabling direct quantitative comparison in main Fig. 4c. Balanced genome-scale source-to-sink flux is retained separately elsewhere in the analysis.

#### Validation *for K. phaffii* under various oxygen levels

**Experimental flux maps and scope.** Baumann et al.^23^ quantified central carbon fluxes in glucose-limited *Pichia pastoris* (now *Komagataella phaffii*) control and Fab-expressing strains at 21%, 11% and 8% inlet O_2_ using ^13^C-metabolic flux analysis. Published net reaction fluxes and standard deviations were reported in Additional file 5 and reaction stoichiometry in Additional file 8. The original MFA calculation quantifies central carbon fluxes independently of NADH, NADPH, ATP, O_2_ and CO_2_ balances. The reported observable is therefore electron generation.

**Electron accounting.** For each published reaction, reduced-cofactor stoichiometry was converted to a signed electron coefficient using two electron equivalents per NADH, NADPH or FADH_2_. Positive terms represent electron generation and negative terms explicit reduced-equivalent consumption. The transhydrogenase-like conversion has a net coefficient of −2 e^-^ per turnover. Condition-specific protein-biomass coefficients calculated from the reported NADH/NADPH requirements were −2.820, −2.802 and −2.916 e^-^ per turnover at 21%, 11% and 8% O_2_, respectively. Electron generation was summed over positive reaction contributions and normalized by measured glucose uptake.

**Experimental response and interpretation.** Electron generation per glucose decreased monotonically as oxygen was restricted: from 8.295 ± 0.475 to 7.562 ± 0.345 and 5.118 ± 0.448 e^-^ per glucose in control cells, and from 8.187 ± 0.489 to 7.412 ± 0.482 and 4.944 ± 0.303 in Fab-expressing cells at 21%, 11% and 8% inlet O_2_, respectively. From 21% to 8% O_2_ this corresponds to 38.3% and 39.6% compression, despite 88.5% and 78.6% increases in glucose uptake (Supplementary Fig. S14). Thus, substrate-normalized electron generation does not scale simply with carbon uptake in this independent yeast dataset. Absolute electron generation increases over the same interval. The analysis quantifies electron generation within the published central carbon model. Error bars are first-order propagated standard deviations from the reported marginal flux standard deviations.

### Supplementary Note 15 — Predictive transfer of the oxygen-response structure across carbon sources

#### Experimental dataset and physiological observable

Weusthuis et al.^25^ measured steady-state *S. cerevisiae* CBS 8066 chemostats at a dilution rate of 0.10 h^-1^ on glucose or maltose across graded O_2_ feed. The glucose–maltose comparison tests whether the normalized response to oxygen supply transfers from a monosaccharide to a disaccharide under the same chemostat dilution rate. The medium contained approximately 12 mM ethanol from the growth-factor stock. Ethanol production rate was therefore calculated as q_ethanol_ = D(C_ethanol,out_ − C_ethanol,in_)/X, where D is the dilution rate and X is biomass concentration. Positive values denote ethanol formation and negative values consumption of ethanol entering with the feed. The experiment provides a physiological readout of the shift between fermentative and oxidative carbon use. It provides the measured physiological readout for the transfer analysis.

#### Glucose-derived response

The response shape was inferred from the glucose series only. Ethanol concentrations reported as below detection (<1 mM) at the two highest O_2_ feeds were set to 0 mM in the primary analysis. Ethanol production was then fitted by decreasing isotonic regression as a function of O_2_ feed.

#### Held-out maltose prediction

The glucose-derived response was converted to a normalized response coordinate between zero O_2_ feed and 30.8 mmol l^-1^ h^-1^ O_2_ feed. Maltose contributed only its carbon-source-specific scaling: the mean of the two zero-O_2_ maltose measurements specified the zero-O_2_ baseline (11.516 mmol gDW^-1^ h^-1^), and the 30.8 mmol l^-1^ h^-1^ state specified the high-O_2_ endpoint (0.954 mmol gDW^-1^ h^-1^). The intermediate maltose states at 5.4, 10.8, 18.7, 22.3 and 24.8 mmol l^-1^ h^-1^ O_2_ feed were not used to infer the response or the two endpoint conditions used for scaling. Their observed ethanol production rates were 8.292, 4.275, 2.881, 1.155 and 0.522 mmol gDW^-1^ h^-1^, compared with predictions of 7.710, 4.801, 2.508, 1.738 and 1.297 mmol gDW^-1^ h^-1^, respectively. Across these five test conditions, prediction and experiment agreed with r = 0.988, predictive R^2^ = 0.956 and RMSE = 0.58 mmol gDW^-1^ h^-1^ (Supplementary Fig. S20).

#### Censoring sensitivity

The predictive result is insensitive to the exact treatment of the two cases where glucose was the carbon source and ethanol was not produced. Assigning values from 0 to 1 mM across the reported <1 mM interval changed predictive R^2^ only from 0.956 to 0.957 and RMSE from 0.582 to 0.573 mmol gDW^-1^ h^-1^. The primary 0 mM treatment is therefore used in Fig. 4f and Supplementary Fig. S20. The test evaluates the shared-response implication of the near-separable organization using O_2_ supply as the experimentally controlled disposal variable. Ethanol production is used as the physiological response variable.

### Supplementary Note 16 — Robustness and reproducibility

#### Flux-state and model robustness

Key conclusions are checked against flux non-uniqueness using flux variability analysis and equal-parsimony alternatives, and against reconstruction dependence using iJO1366 as an independent *E. coli* model. Oxygen-supply sensitivity is reported separately from the paired endpoint comparisons. These robustness analyses support the qualitative compression and routing conclusions across alternative optimal pFBA flux distributions.

#### Software and numerical reproducibility

The repository (<https://github.com/MetaSys-LISBP/Cellular_electron_economy>) provides the code, model and data snapshots with SHA256 checksums, condition configurations, canonical result tables, source data, figures, automated tests and numerical audit scripts. The reference computational environment uses Python 3.13.5, COBRApy 0.32.1, swiglpk 5.0.13, python-libSBML 5.21.1, optlang 1.9.1, pandas 2.2.3, NumPy 2.3.5, NetworkX 3.6.1, python-louvain 0.16, SciPy 1.17.0, scikit-learn 1.8.0, statsmodels 0.14.6, matplotlib 3.10.8 and pytest 9.0.2.

#### Supplementary Data and source data traceability

The Supplementary Data package distinguishes detailed analysis outputs from figure-facing Source Data. FIGURE_SOURCE_DATA_MANIFEST.csv maps every quantitative figure panel to the corresponding machine^-^readable Source Data. Fromanger whole-cell balances are traced through the *C. shehatae* carbon-partition input, derived electron-equivalent balance and Supplementary Fig. S18 Source Data. Baumann electron generation results and both predictive-transfer calculations are traceable from experimental inputs to Supplementary Figs. S14, S19 and S20 and main Fig. 4e–f. Extended acceptor, carbon source, factorial and routing analyses are linked to their publication result tables and Source Data.

### Supplementary Figures

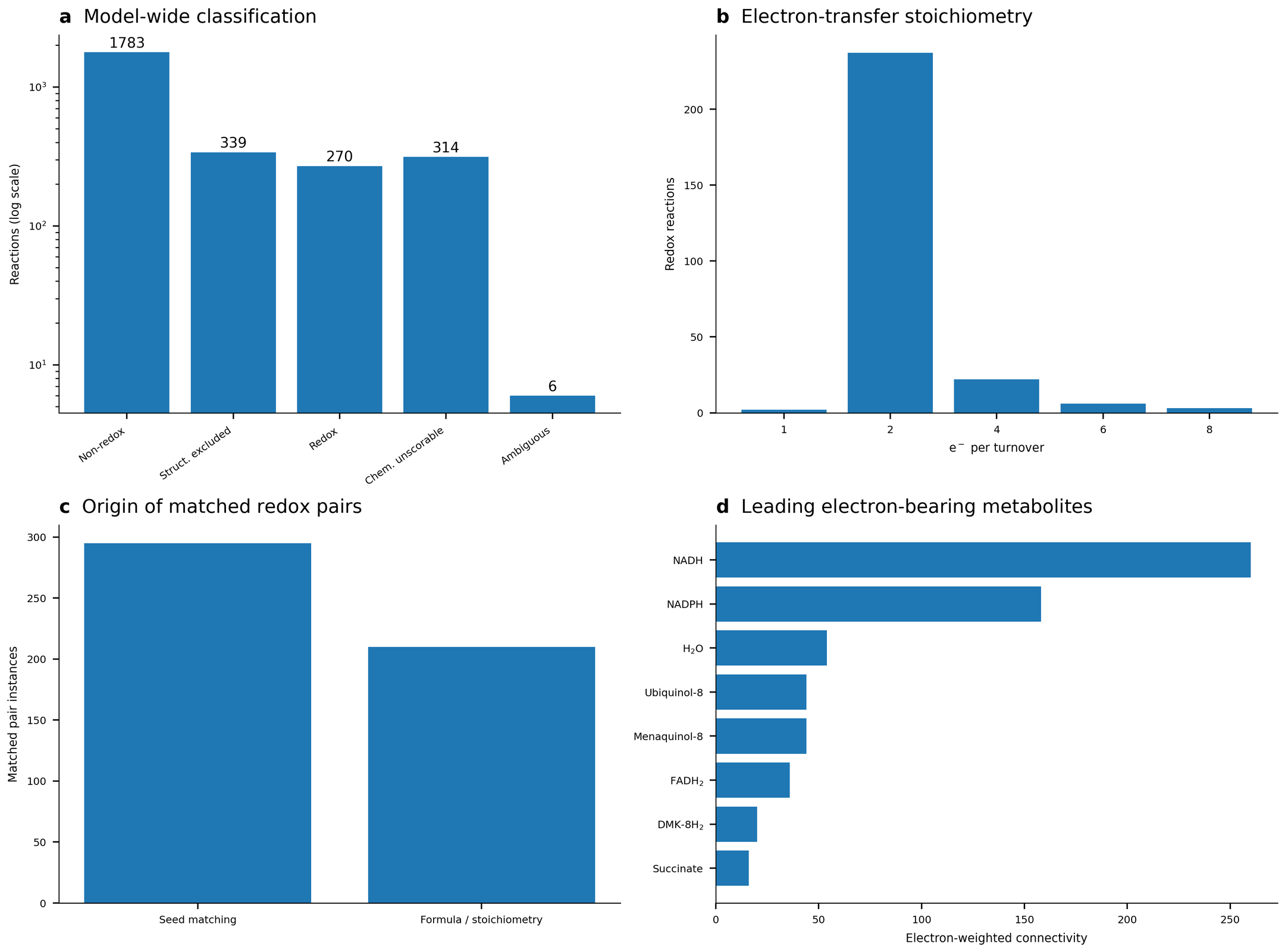

**Supplementary Figure S1.** Detailed reconstruction diagnostics. a, Model-wide reaction-classification outcome. b, Electron-transfer stoichiometry of confidently identified redox reactions. c, Origin of matched redox-pair instances, distinguishing direct seed matching from formula/stoichiometry-based inference. d, Electron-weighted connectivity of leading electron carriers. These diagnostics document reconstruction quality and the structure of the electron-transfer network.

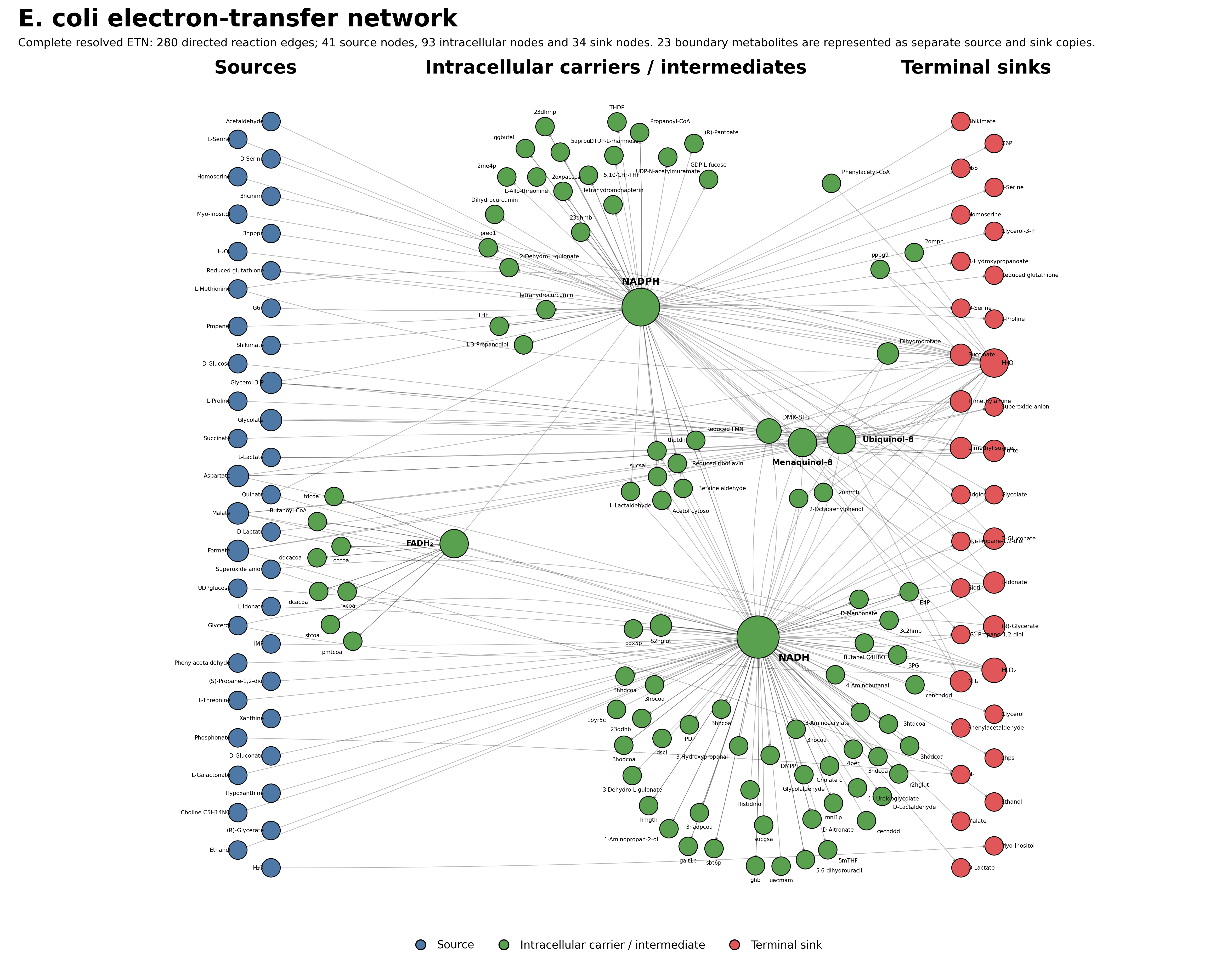

**Supplementary Figure S2.** Reconstructed electron transfer network of *Escherichia coli* iML1515. The complete network contains 280 directed reaction edges connecting 41 source nodes, 93 intracellular carrier or intermediate nodes and 34 terminal sink nodes. The 280 reaction-labelled edges correspond to 263 unique directed donor–acceptor edges reported in the main text; boundary metabolites that function as both sources and sinks are duplicated only for visualization. Colours distinguish the three node roles, and node size represents connectivity. The 23 boundary metabolites that act as both sources and sinks are displayed as separate copies in the corresponding boundary columns.

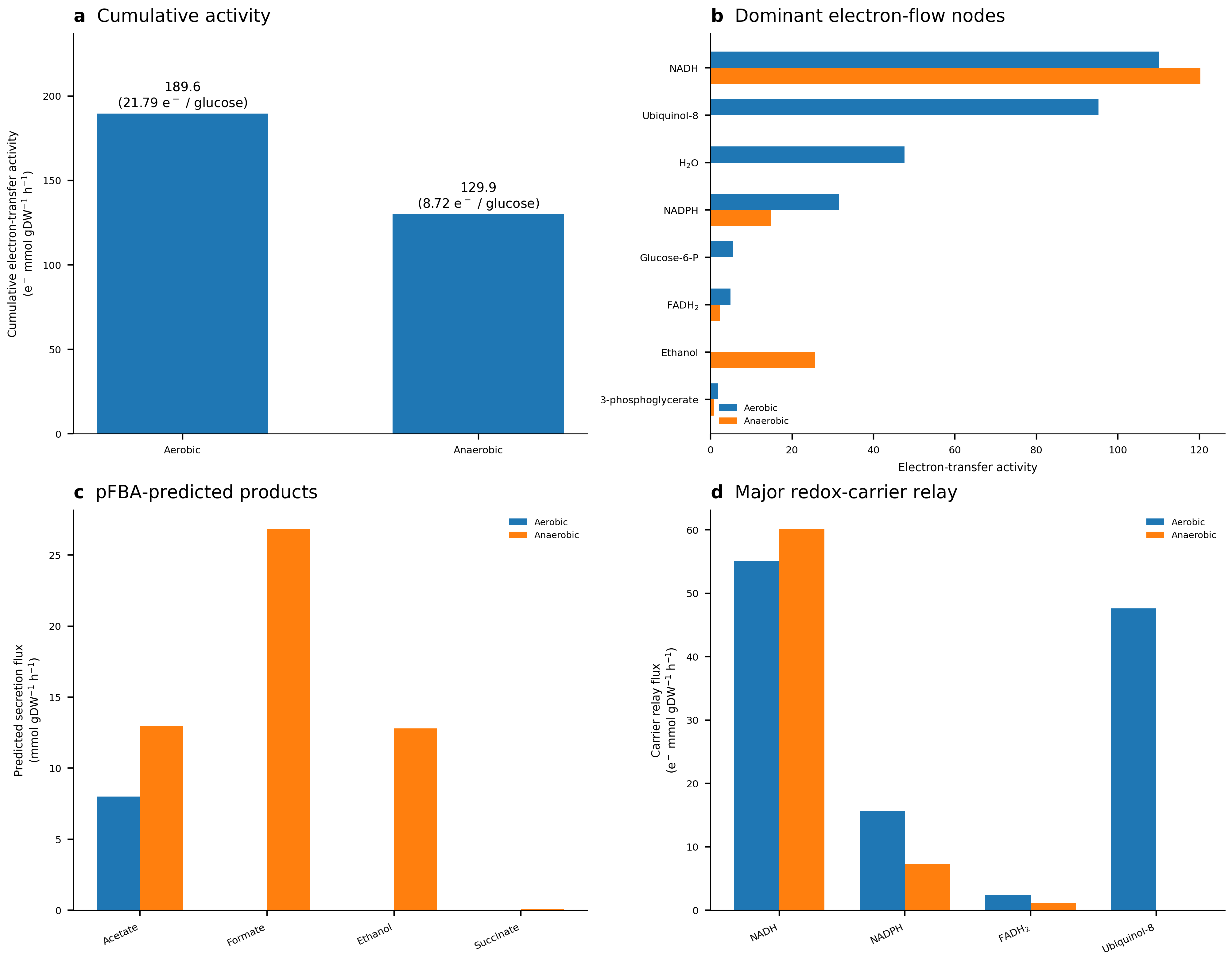

**Supplementary Figure S3.** Additional diagnostics for the aerobic and anaerobic glucose electron flux maps. a, Cumulative electron-transfer activity in absolute units and normalized to measured glucose uptake. b, Dominant electron-flux nodes ranked by total electron-transfer activity (inflow + outflow) in each condition. c, Extracellular product exchange rates in the calculated electron fluxes. These product rates are model outputs rather than measured fluxes, except that aerobic formate, ethanol, succinate and lactate exchanges were constrained to zero from reported non-detection. Glucose and oxygen uptake are the quantitative reported constraints. d, Relay through major redox carriers.

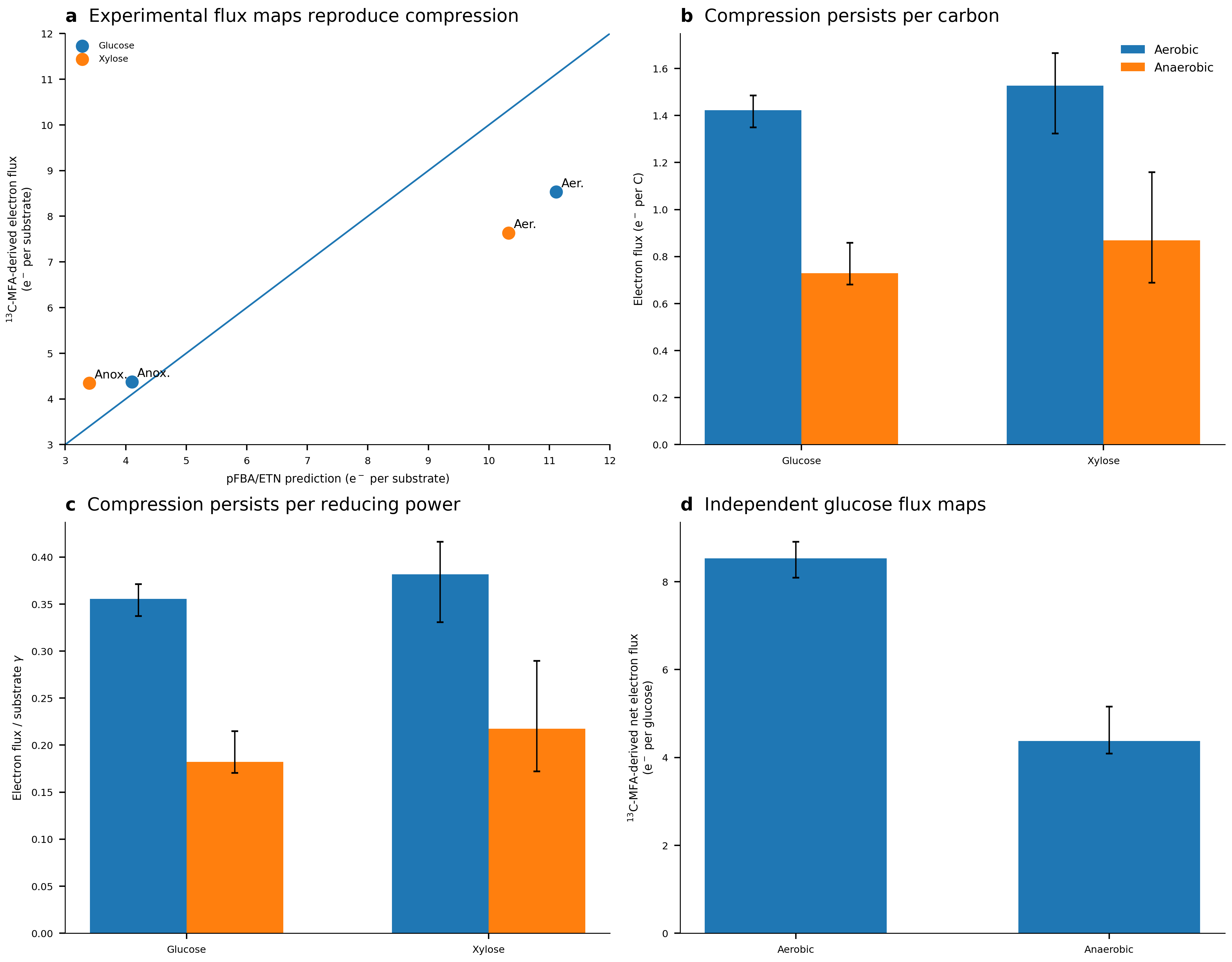

**Supplementary Figure S4.** Validation of electron flow compression on glucose and xylose from experimental data, and normalization robustness. a, Comparison of genome-scale electron flux predictions with electron fluxes calculated from experimental ^13^C-flux maps. The experimentally-derived electron flux maps reproduce compression for glucose and xylose but overestimate the aerobic expanded state and underestimate the anaerobic xylose total. b, The same electron fluxes normalized by substrate carbon number. c, Electron flux normalized by total substrate degree of reduction γ. Error bars in b and c are the conservative envelopes obtained by deterministic scaling of the independently propagated marginal reaction-flux limits. In both normalizations, aerobic electron flux remains higher than anaerobic flux for glucose and xylose.

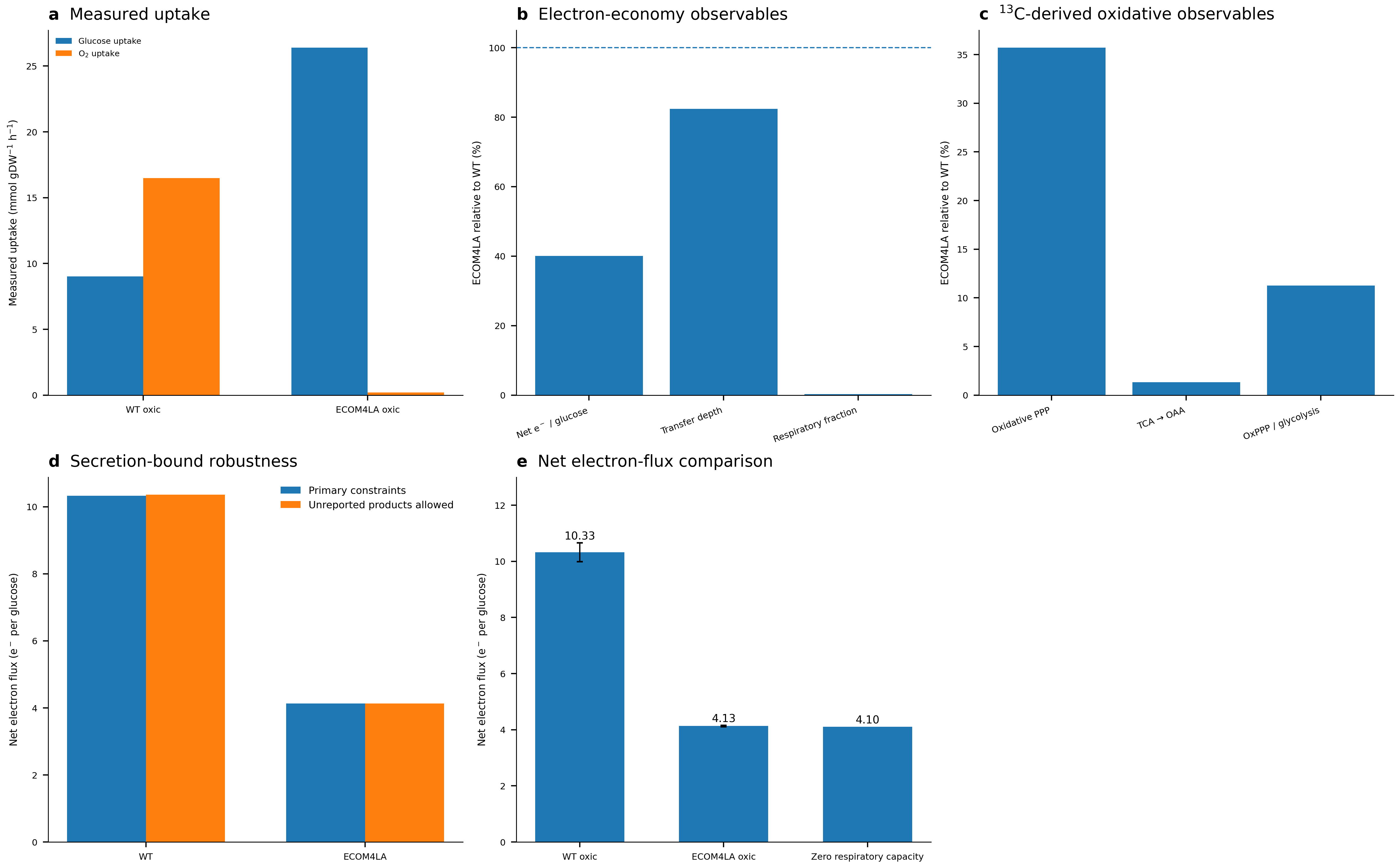

**Supplementary Figure S5.** ECOM4LA provides a case where oxygen is available but respiratory disposal capacity is limited, thereby distinguishing disposal capacity from oxygen availability. a, Measured glucose and O_2_ uptake in aerobic WT and aerobic ECOM4LA. b, ECOM4LA cellular electron economy observables compared to WT. c, ^13^C-MFA data used to calculate electron fluxes and expressed relative to WT flux values. d, Robustness of the calculated net electron-flux endpoint to allowing otherwise unreported secretion products. The panels deliberately distinguish reported measurements, electron-flux observables calculated by the framework and genome-scale model completion. e, Net electron flux in aerobic wild type, aerobic ECOM4LA and the zero-respiratory-capacity model state.

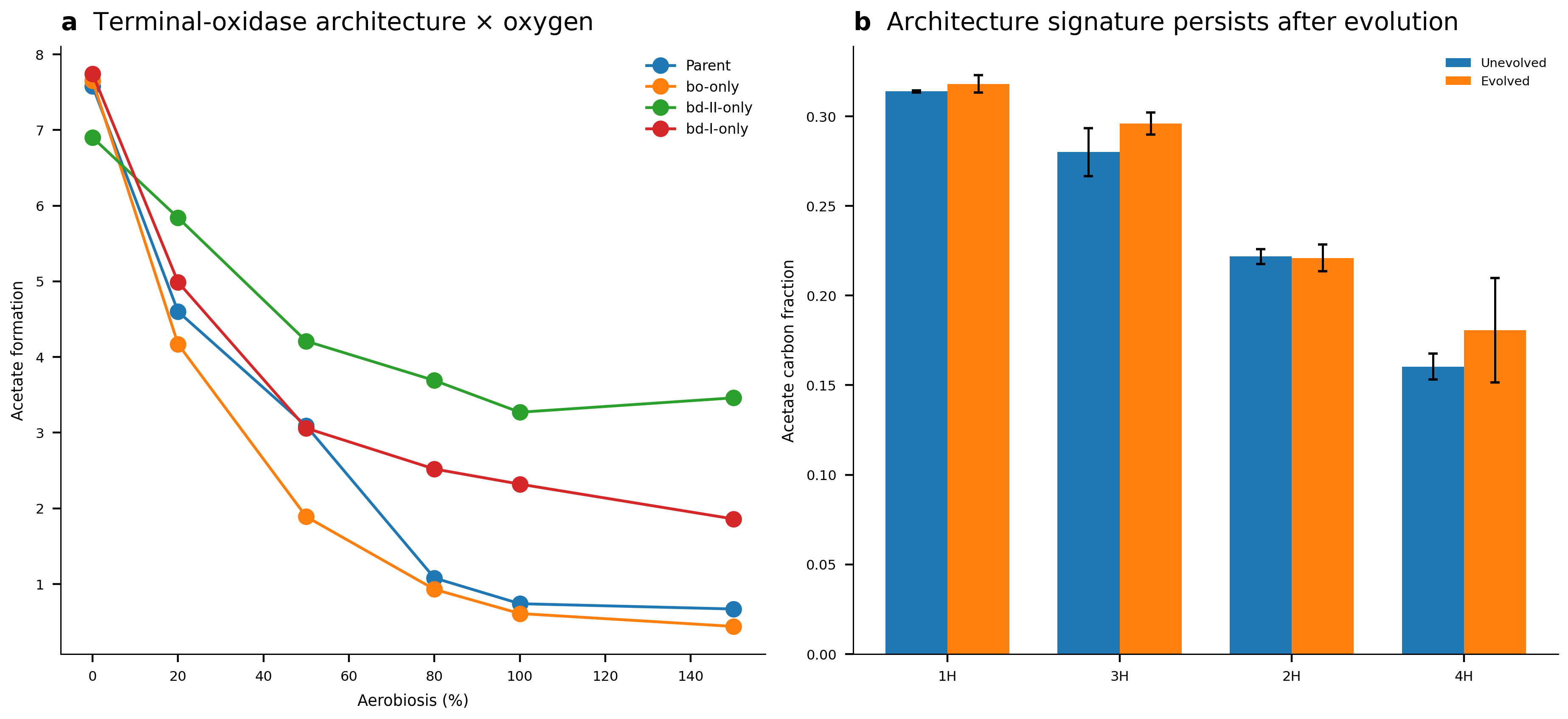

**Supplementary Figure S6.** Independent respiratory-architecture controls. a, Published mean acetate formation for the parental TBE029 strain and three single-oxidase derivatives across the Steinsiek aerobiosis series. Lines connect categorical aerobiosis levels only as a visual guide. b, Carbon secreted as acetate, calculated from reported extracellular glucose and acetate measurements for the Anand et al.^38^ synthetic unbranched-ETS architectures before and after adaptive evolution. Error bars are s.d. across the supplied replicate measurements. These panels provide physiological architecture controls based on replicated extracellular measurements.

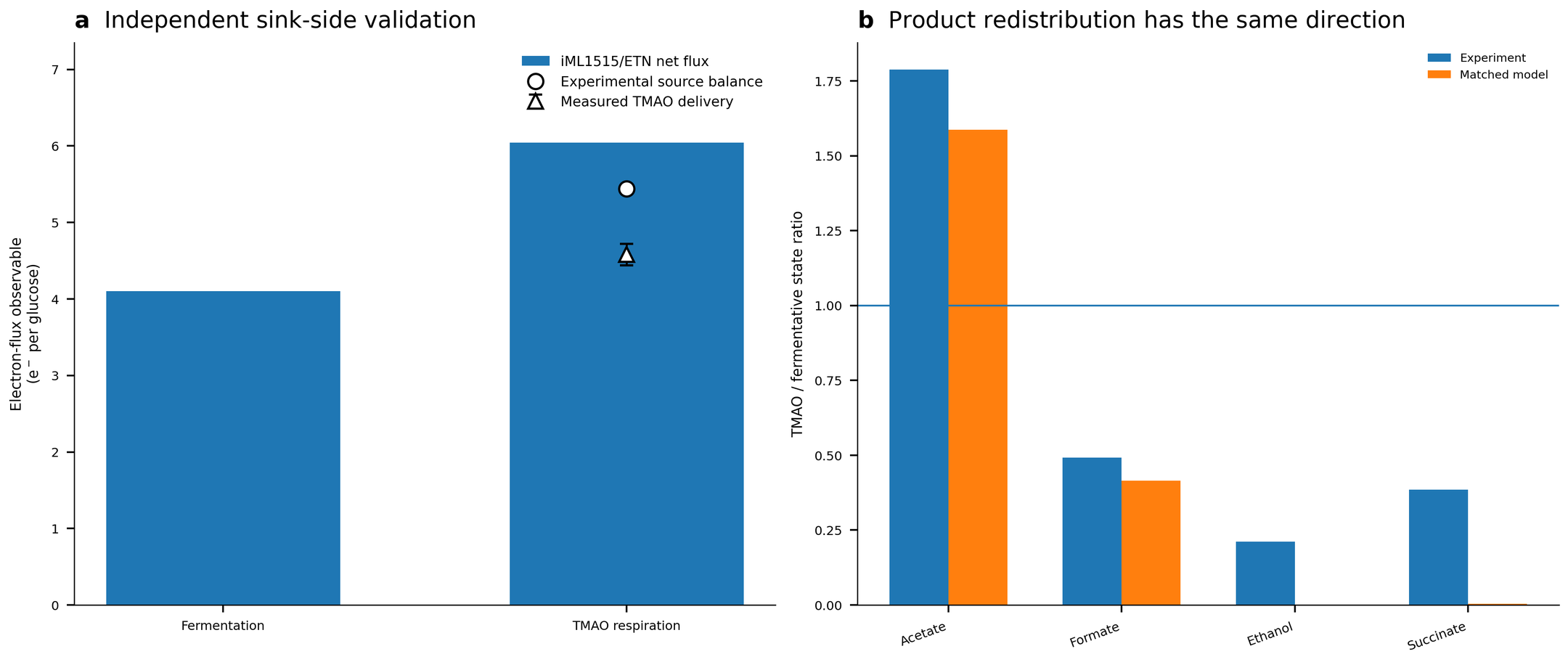

**Supplementary Figure S7.** Independent TMAO-respiration validation. a, iML1515/ETN net source-to-sink flux in fermentation and at the experimentally observed TMAO/glucose ratio. Open symbols show the distinct experimental source-balance and terminal-delivery observables. b, Experimental and matched-model product redistribution. The model reproduces the direction of change for acetate, formate, ethanol and succinate, and the comparison evaluates the direction of secretion changes.

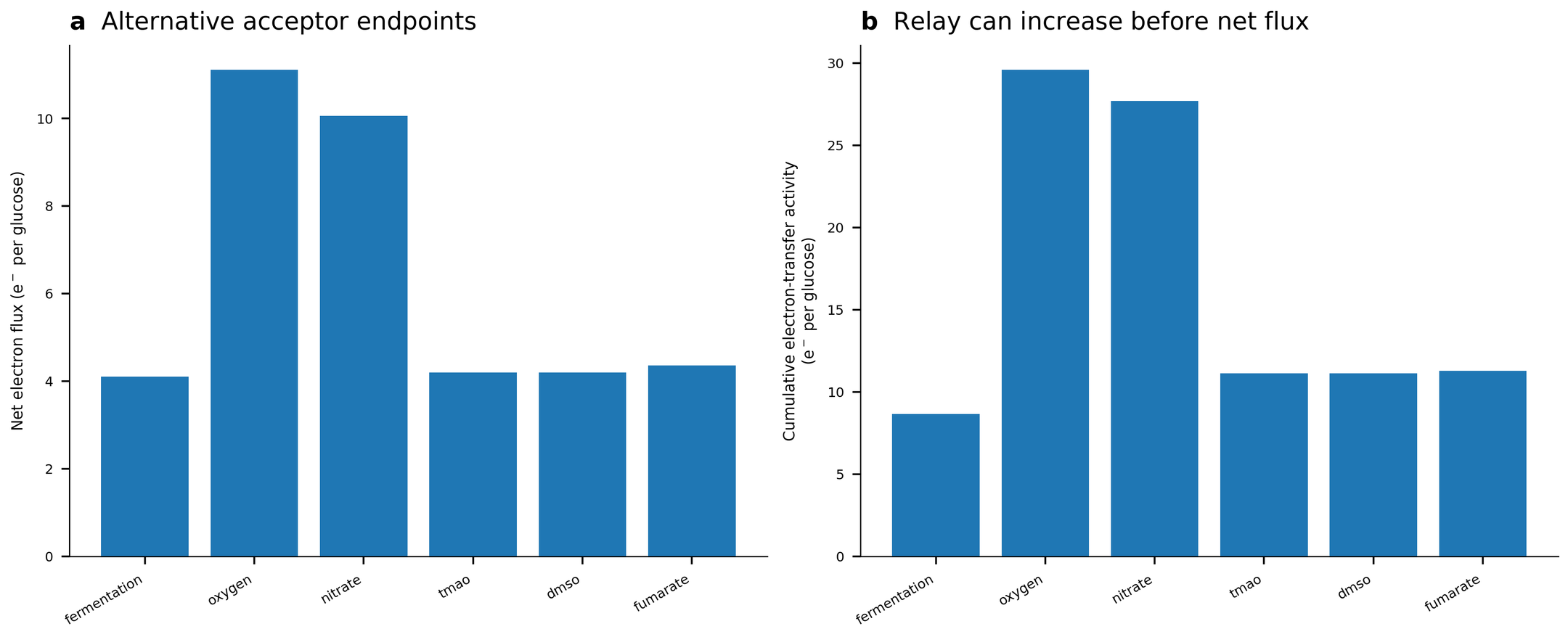

**Supplementary Figure S8.** Alternative terminal electron acceptors differentially expand the electron flow. a, Net source-to-sink electron flux at representative endpoints for fermentation (no acceptor), oxygen, nitrate, TMAO, DMSO and fumarate. b, Cumulative electron-transfer activity for the same regimes. TMAO, DMSO and fumarate at the illustrated capacity can increase relay activity before producing a large increase in net electron flux.

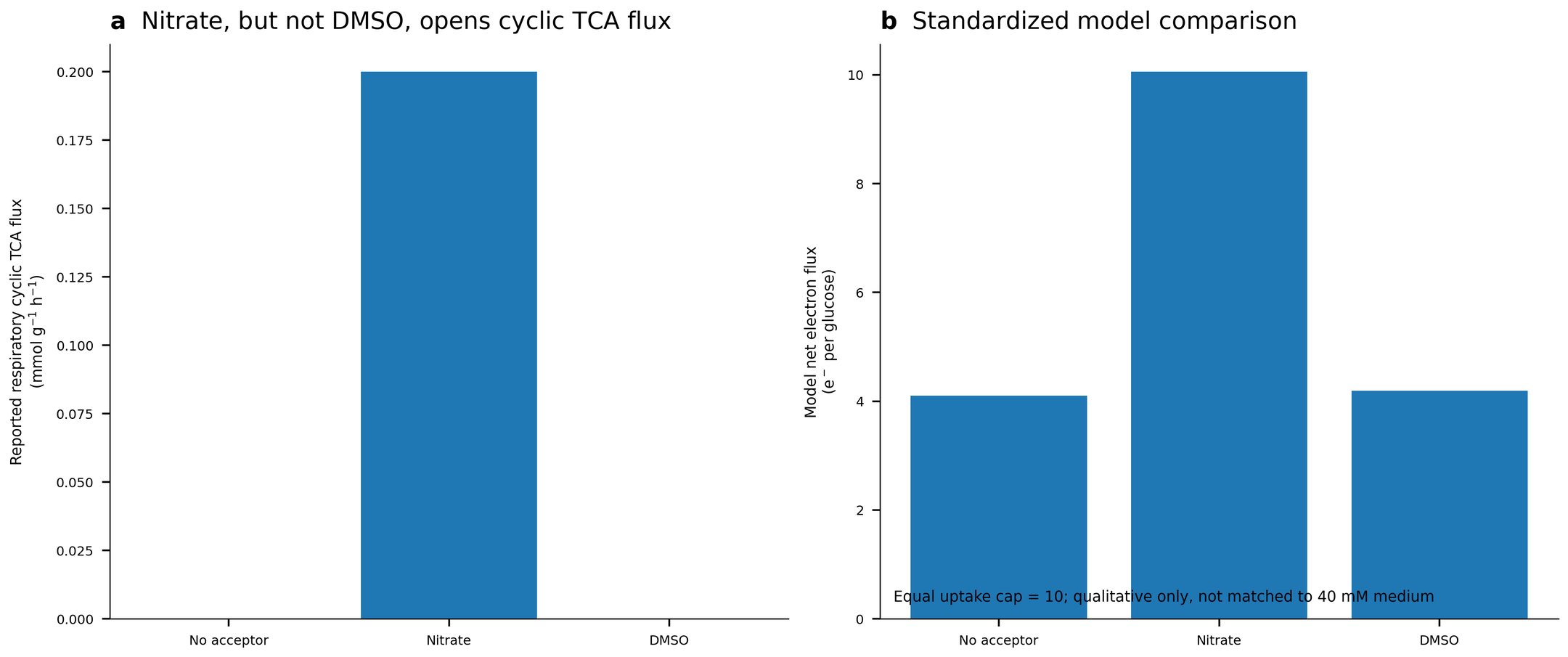

**Supplementary Figure S9.** Acceptor identity and effective capacity shape oxidative routing. a, Respiratory cyclic TCA flux reported by Perrenoud & Sauer for anaerobic cultures without an external electron acceptor, with nitrate or with DMSO. Cyclic flux was observed only with nitrate, whereas the TCA pathway remained bifurcated in the other two conditions. b, Standardized iML1515/ETN comparison at an equal acceptor uptake upper bound of 10 mmol gDW^-1^ h^-1^. Panel b is a qualitative model comparison only. Panel a reports 40 mM medium concentrations, whereas panel b applies the standardized model uptake upper bound.

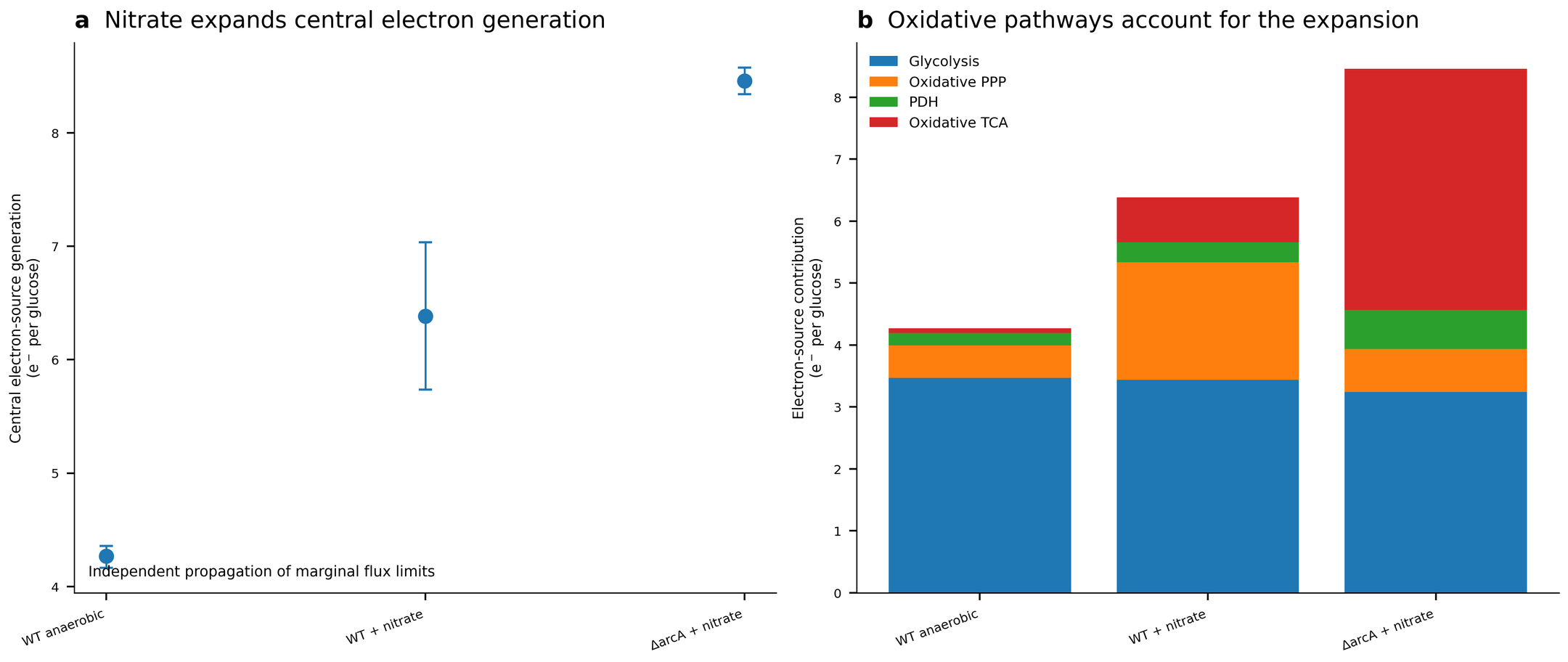

**Supplementary Figure S10.** Nitrate-dependent electron generation calculated from the data in Toya et al. a, Total electron generation with conservative independently propagated marginal-flux envelopes. b, Decomposition into glycolysis, oxidative pentose-phosphate pathway, PDH and oxidative TCA contributions. Nitrate increases electron generation without increasing the glycolytic contribution. Δ*arcA* reveals additional regulatory control over the magnitude realized under nitrate.

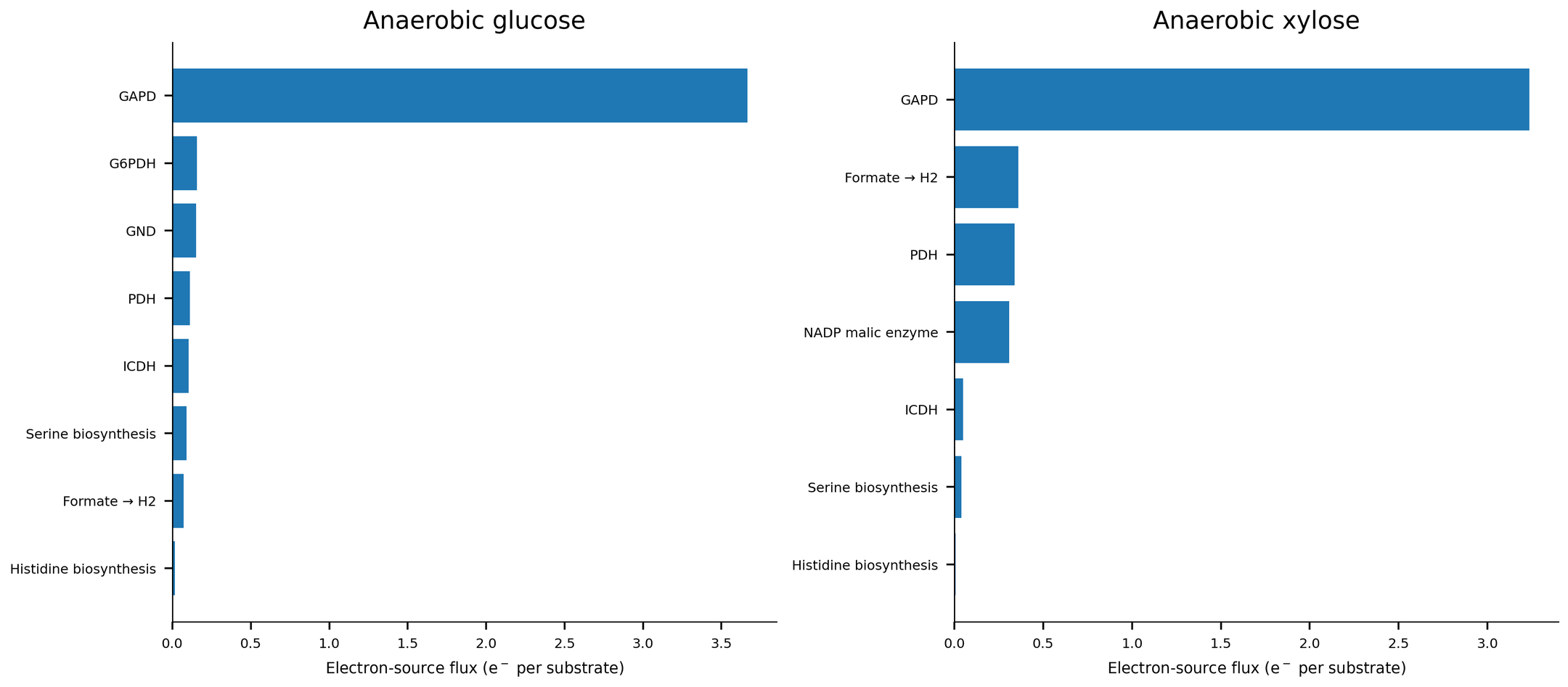

**Supplementary Figure S11.** Reaction-level decomposition of anaerobic net electron-source flux calculated from the Gonzalez ^13^C-flux data. GAPD is the dominant source for both substrates, while anaerobic xylose contains larger auxiliary contributions from formate-to-H_2_, pyruvate dehydrogenase and NADP-dependent malic enzyme.

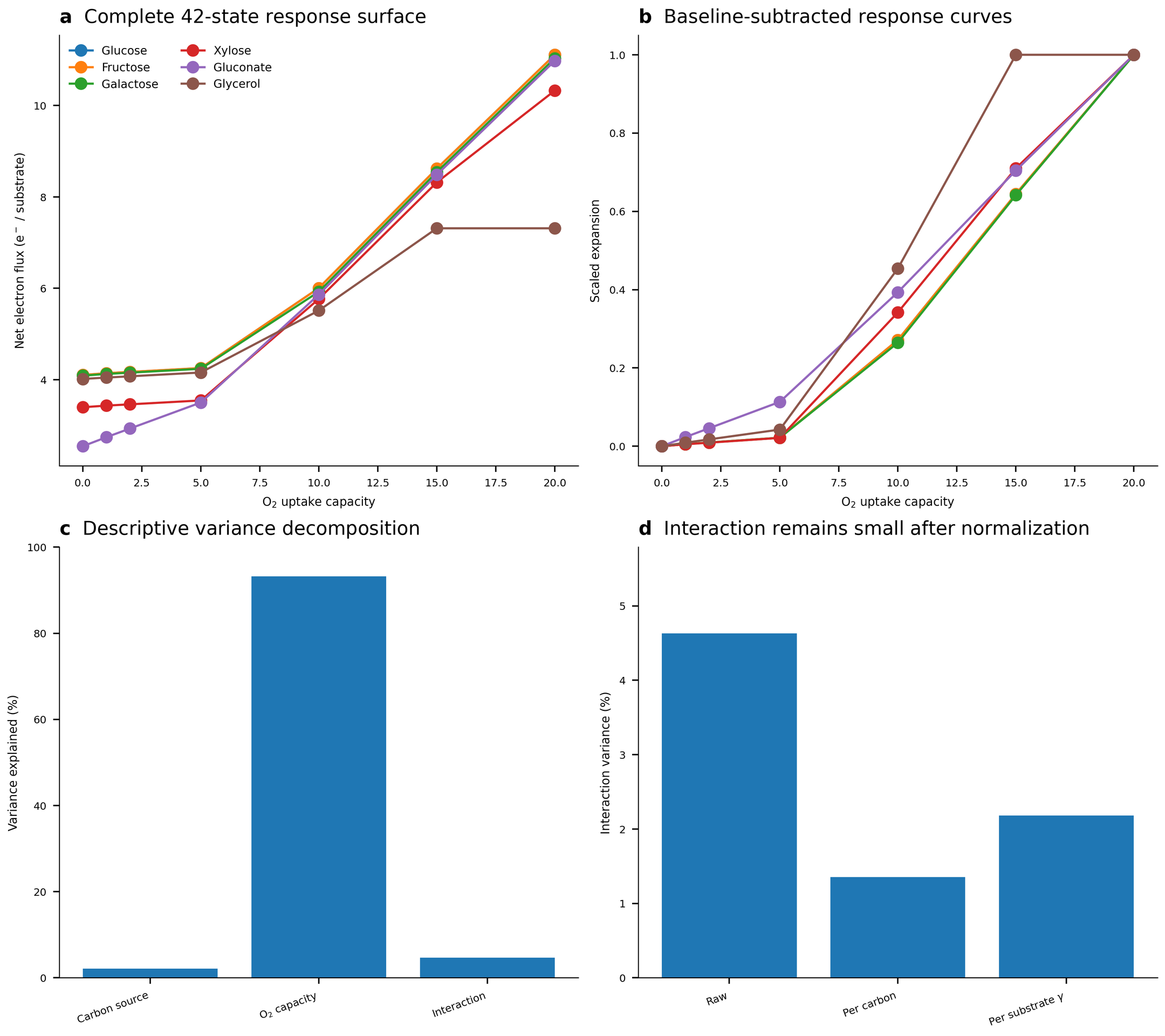

**Supplementary Figure S12.** Formal carbon-entry × disposal capacity response surface in *E. coli*. a, Net electron flux across all 42 substrate × O_2_ capacity states. b, Baseline^-^subtracted, substrate-scaled expansion curves. c, Descriptive two-way variance decomposition of the raw response matrix. d, Residual interaction variance in the raw response and after normalization by substrate carbon number or substrate degree of reduction γ. Points represent deterministic model states.

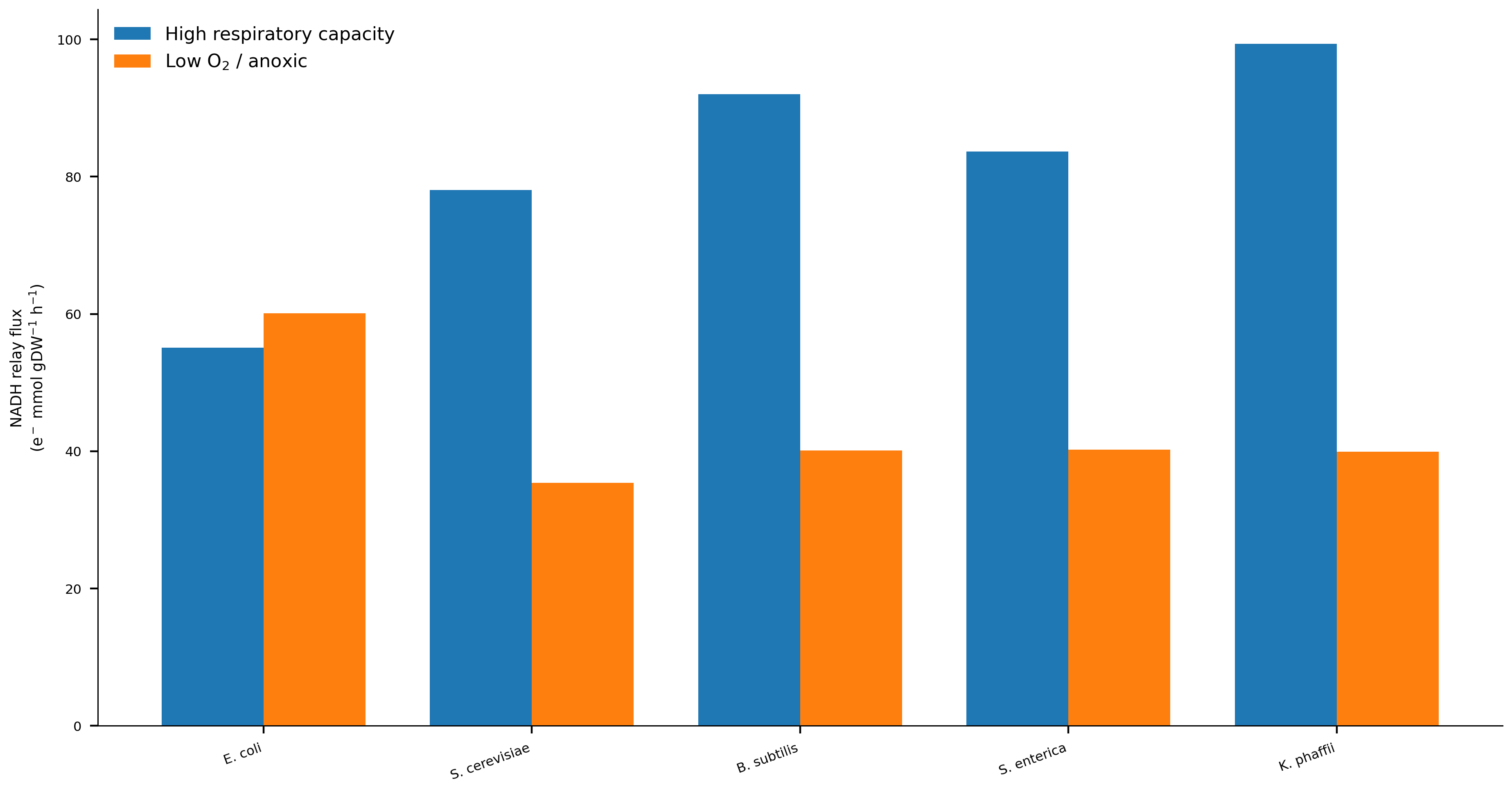

**Supplementary Figure S13.** NADH-relay responses to oxygen restriction are organism-specific. Lines connect aerobic and low-O_2_/anaerobic states for the five organisms with paired physiological analyses. NADH relay remains high in *E. coli* but decreases strongly in *S. cerevisiae*, *B. subtilis*, *S. enterica* and *K. phaffii*, despite the shared compression of net electron flux shown in main-text Fig. 4a.

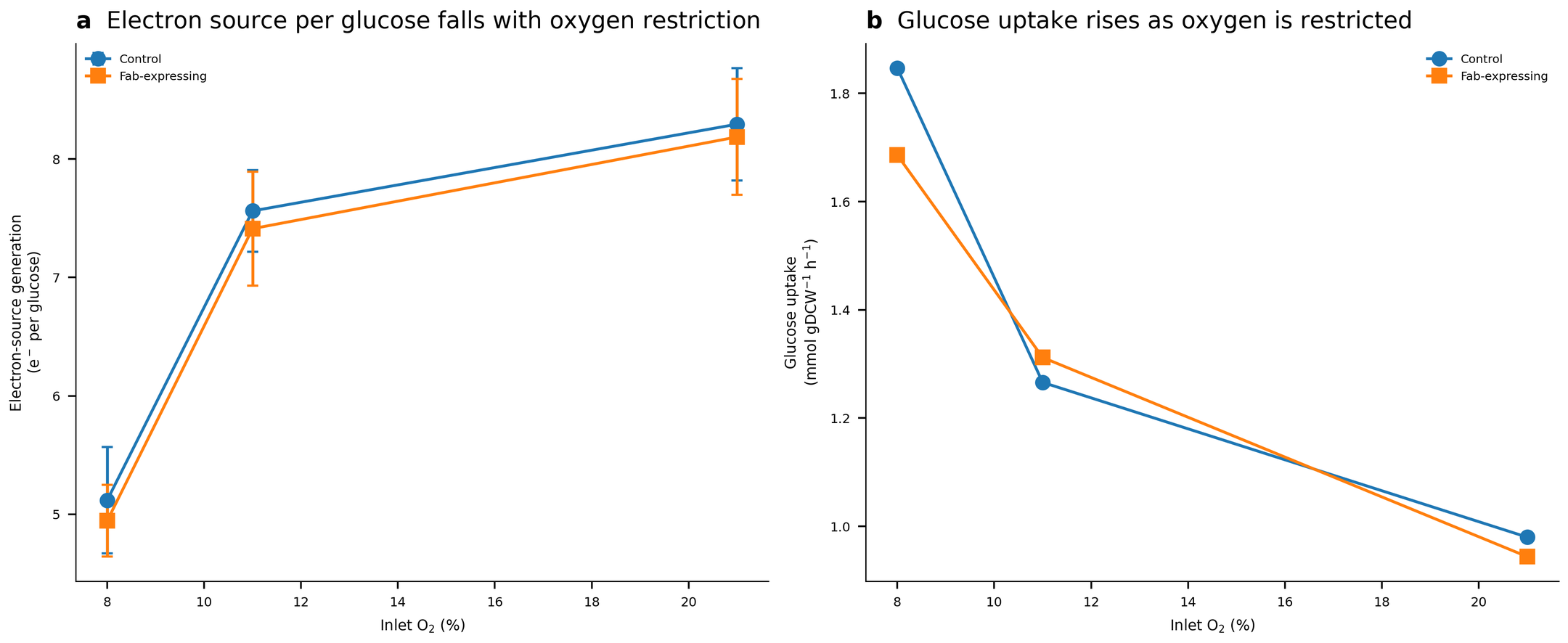

**Supplementary Figure S14.** Validation of oxygen limitation effects for *K. phaffii* based on ^13^C-MFA data. a, Electron generation per glucose calculated from Baumann et al. central carbon flux maps for control and Fab-expressing strains at 21%, 11% and 8% inlet O_2_. Error bars are first-order propagated s.d. from reported marginal flux s.d. b, Measured glucose uptake over the same oxygen series. Electron generation per glucose falls by 38.3% and 39.6% from 21% to 8% O_2_ while glucose uptake rises by 88.5% and 78.6%, respectively. Panel a reports electron generation calculated from the published central carbon metabolic flux distributions.

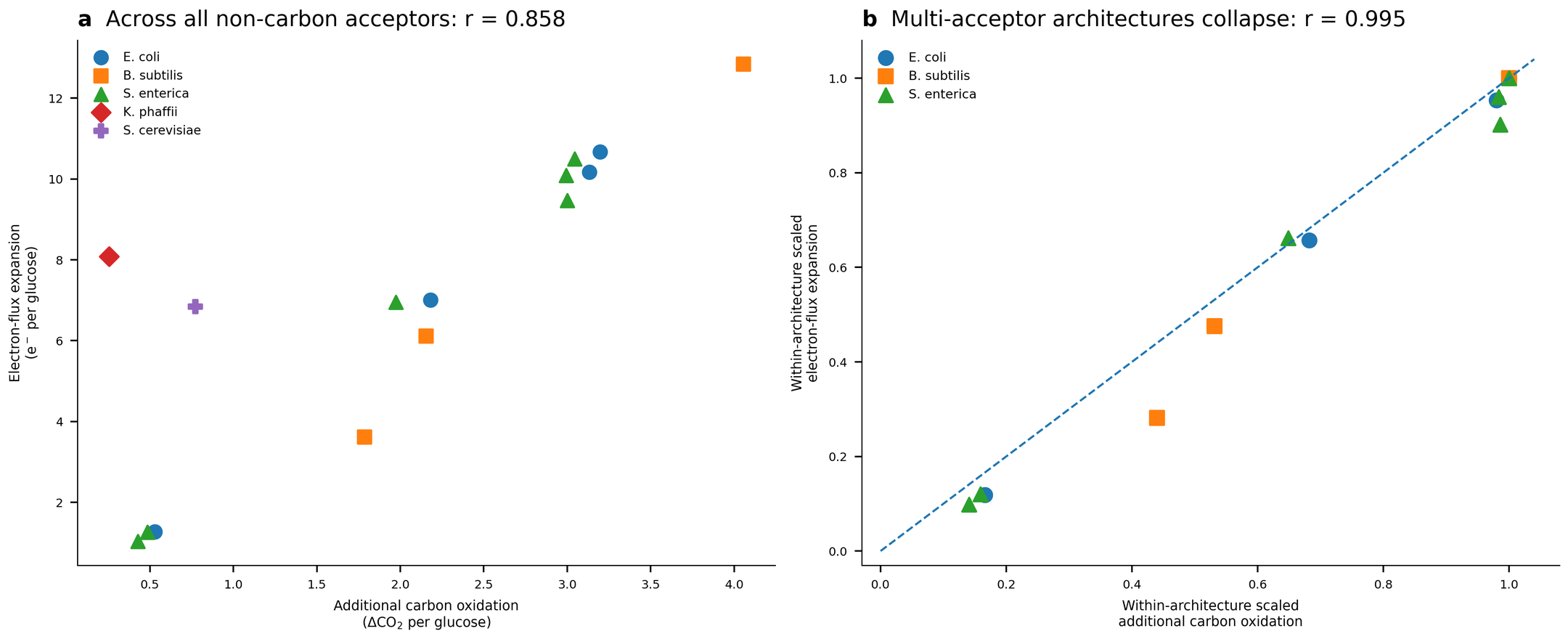

**Supplementary Figure S15.** Extended terminal-acceptor generality. a, Across all 17 feasible non-carbon acceptor states with uptake, electron-flux expansion is related to additional carbon oxidation (Pearson r = 0.858), with organism-specific scales. b, After scaling electron-flux expansion and additional carbon oxidation by their within-organism maxima in *E. coli*, *B. subtilis* and *S. enterica*, 15 states collapse onto a common relationship (Pearson r = 0.995). Points are deterministic model states. Acceptor identities and numerical values are given in Supplementary Table 19 and Source Data.

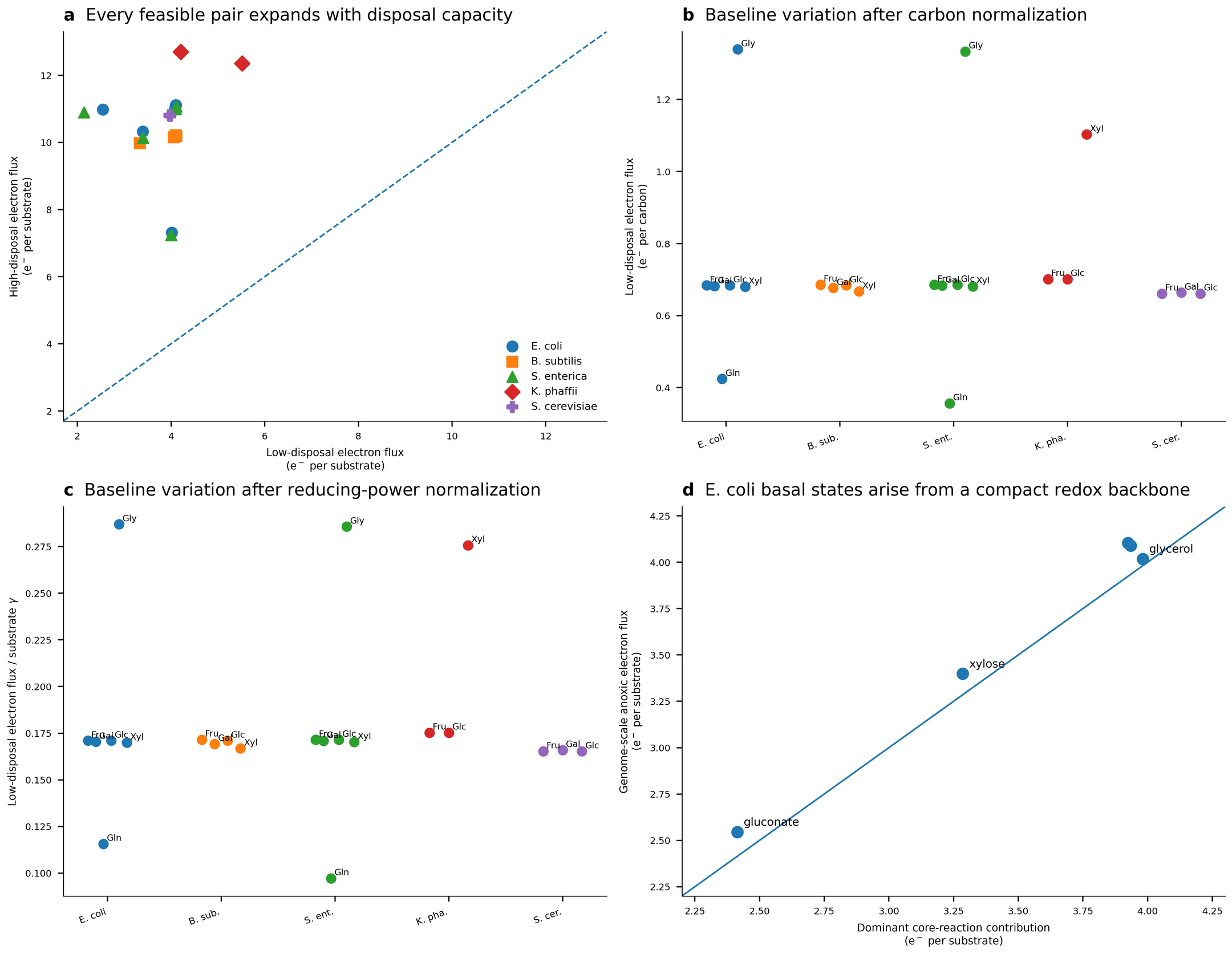

**Supplementary Figure S16.** Cross-species test and normalization robustness of the carbon-dependent baseline and disposal-dependent expansion. a, Baseline versus aerobic net source-to-sink electron flux for all 22 feasible organism–substrate pairs from the systematic 30-combination screen across five organisms. Every point lies above the 1:1 line. b, Baseline electron flux normalized by substrate carbon number. c, Baseline electron flux normalized by total substrate degree of reduction γ. Substantial substrate dependence remains after normalization in *E. coli* and *S. enterica*, with intermediate behaviour in *K. phaffii*, whereas the tested *B. subtilis* and *S. cerevisiae* hexose states largely collapse. d, In *E. coli*, the dominant core redox-generating reaction set quantitatively accounts for the substrate-dependent anaerobic baseline. Points are deterministic model states. Unavailable and infeasible states are listed in Supplementary Table 10.

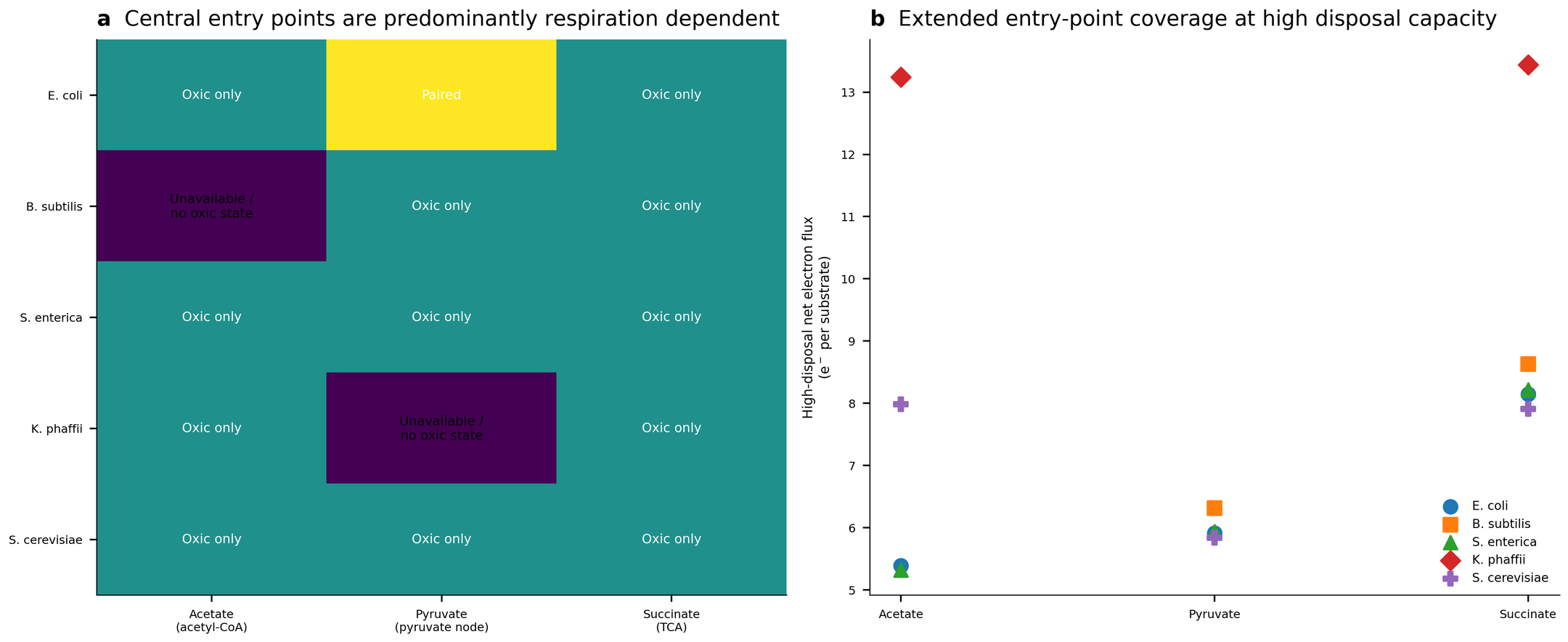

**Supplementary Figure S17.** Extended central carbon-source coverage. a, Feasibility matrix for acetate, pyruvate and succinate under aerobic and minimal respiratory-capacity conditions across five reconstructions. b, Aerobic net electron flux for feasible states. Central entry substrates broaden the range of metabolic entry points tested but are predominantly respiration dependent, which defines their complementary role alongside the six-substrate paired factorial.

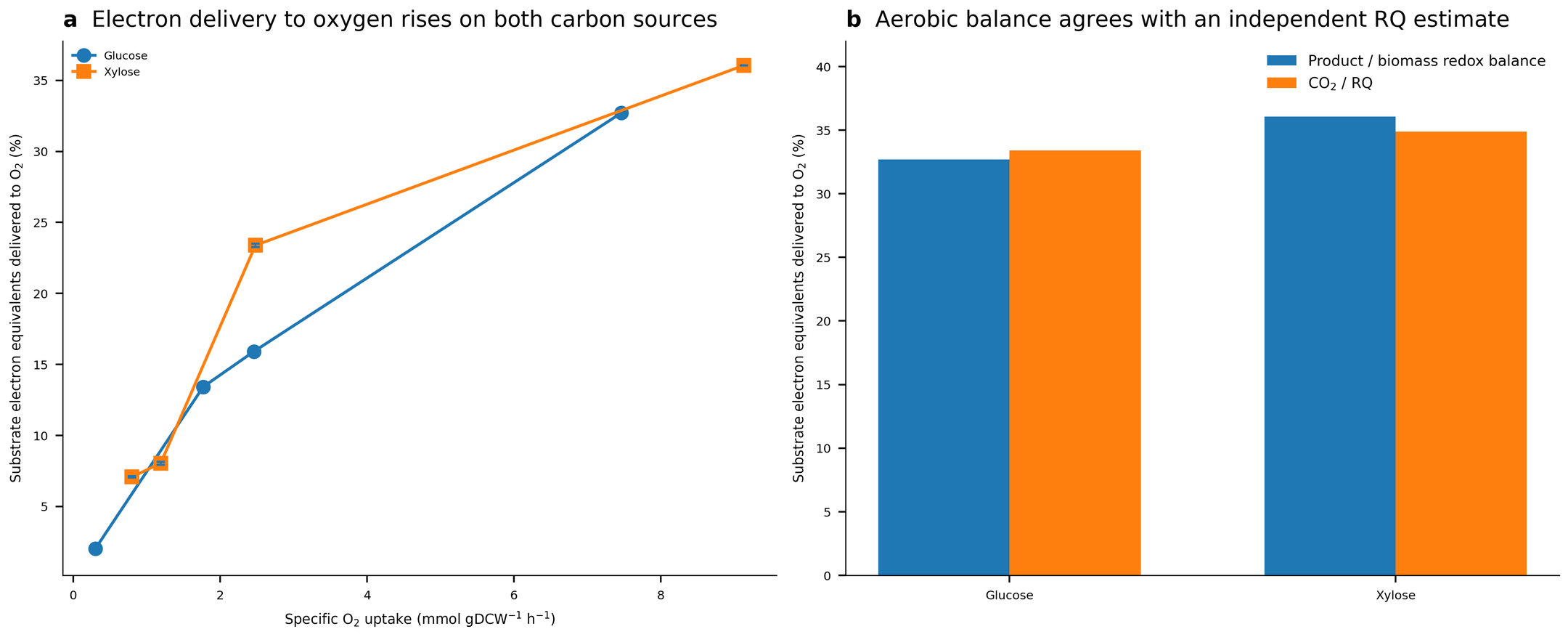

**Supplementary Figure S18.** Independent yeast whole-cell test of carbon source and oxygen availability. a, Fraction of substrate electrons delivered to O_2_ inferred from the product/biomass degree-of-reduction balance reported by Fromanger et al. across glucose and xylose oxygen-series states. The small vertical uncertainty bars show the full sensitivity to assigning the pooled organic-acid fraction γ/C between 3.0 and 3.5. Experimental redox-balance closure was reported within 10% under oxygen limitation. b, Aerobic endpoint cross-check. Product/biomass degree-of-reduction estimates agree with an independent estimate obtained from reported CO_2_ yield divided by respiratory quotient (difference ≤1.2 percentage points). These values are whole-cell terminal-partition observables.

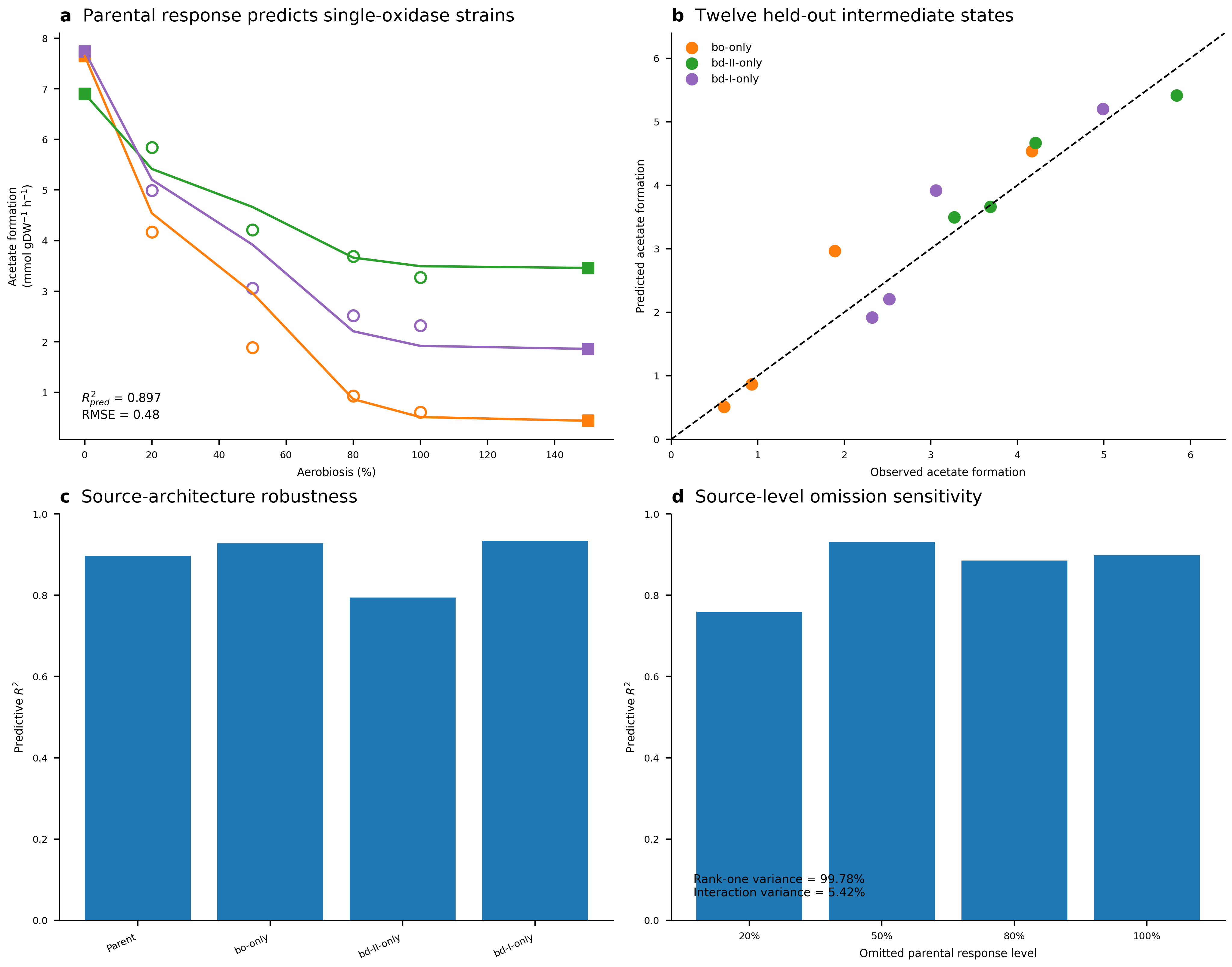

**Supplementary Figure S19.** Respiratory-architecture predictive-transfer diagnostics. a, The response shape inferred from parental TBE029 is scaled only by the 0% and 150% endpoints of each single-oxidase target. Open circles are held-out observations and filled squares are scaling endpoints. b, Predicted versus observed acetate formation for all 12 held-out intermediate states. The dashed line is identity. c, Predictive R^2^ when each architecture is used as the source response. d, Sensitivity to omission and interpolation of individual parental response levels. Statistics use the condition means and are reported as point estimates.

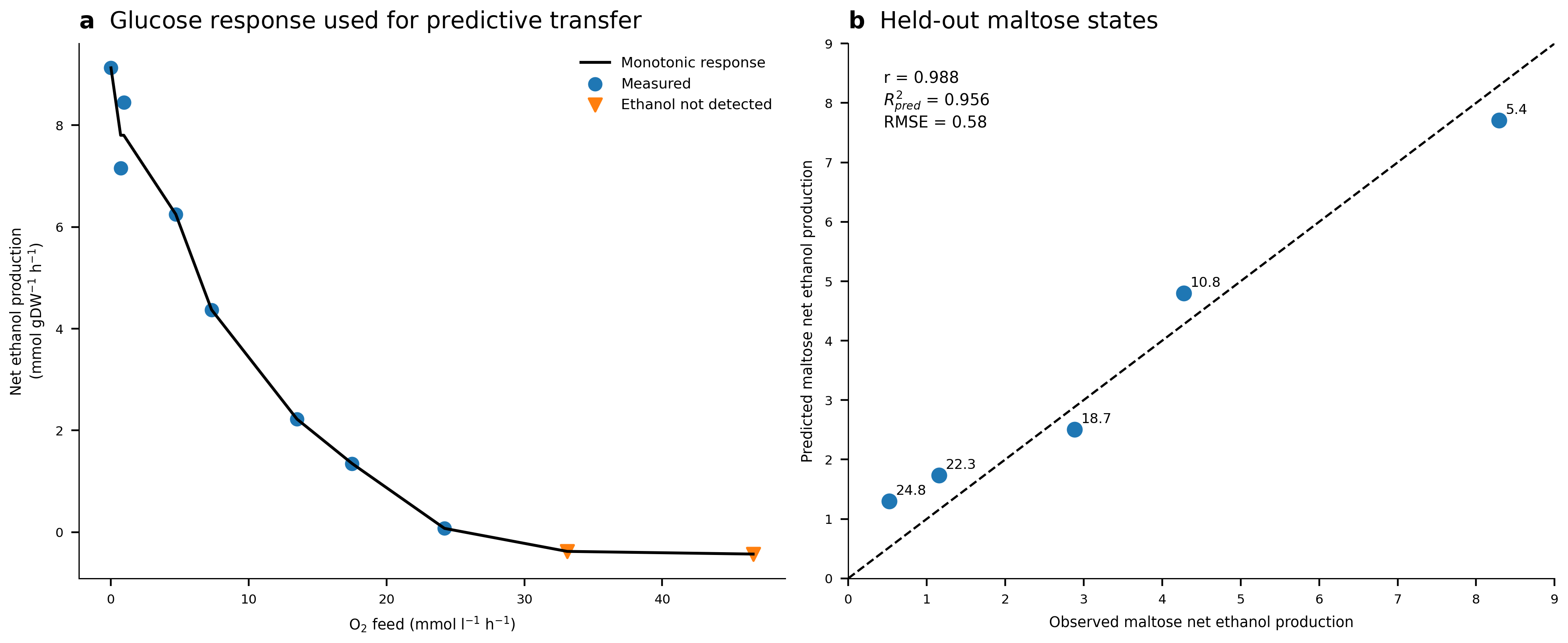

**Supplementary Figure S20.** Diagnostics for predictive transfer from glucose to maltose in *S. cerevisiae*. a, Ethanol production across the glucose O_2_-feed series used to infer the response shape. The black line is the decreasing isotonic fit. Downward triangles indicate the two cases where glucose was the carbon source but ethanol was not detected, treated as 0 mM culture ethanol in the primary analysis. b, Predicted versus observed ethanol production. The dashed line is identity.

### Supplementary Tables

**Supplementary Table 1.** 52-reaction validation set. All 48 chemically scorable reactions are classified correctly (TP=35, TN=13, FP=0, FN=0, with accuracy, precision, recall, specificity and F1=1.00). The four reactions outside the supported chemical domain are flagged as unscorable. This categorical validation set evaluates the chemistry-based classification rules.

| Reaction | Category | Ground truth | Predicted status | e^-^ |
| --- | --- | --- | --- | --- |
| PGI | isomerase | non-redox | non-redox | 0 |
| TPI | isomerase | non-redox | non-redox | 0 |
| RPI | isomerase | non-redox | non-redox | 0 |
| PFK | kinase | non-redox | non-redox | 0 |
| PYK | kinase | non-redox | non-redox | 0 |
| HEX1 | kinase | non-redox | non-redox | 0 |
| ADK1 | kinase | non-redox | non-redox | 0 |
| FUM | hydratase (near redox pair) | non-redox | non-redox | 0 |
| PIuabcpp | transporter | non-redox | non-redox | 0 |
| NO3t7pp | transporter (antiport of a redox pair) | non-redox | non-redox | 0 |
| NH4tpp | transporter | non-redox | non-redox | 0 |
| GLCptspp | transporter (PTS, group translocation) | non-redox | non-redox | 0 |
| PPA | phosphate group transfer (pyrophosphatase) | non-redox | non-redox | 0 |
| ALCD2x | NAD-dehydrogenase | redox | redox | 2 |
| GAPD | NAD-dehydrogenase + phosphoryl transfer | redox | redox | 2 |
| MDH | NAD-dehydrogenase | redox | redox | 2 |
| G6PDH2r | NADP-dehydrogenase | redox | redox | 2 |
| ICDHyr | NADP-dehydrogenase | redox | redox | 2 |
| LDH_D | NAD-dehydrogenase | redox | redox | 2 |
| HACD1 | NAD-dehydrogenase (beta-oxidation) | redox | redox | 2 |
| ALCD19 | NAD-dehydrogenase | redox | redox | 2 |
| GLUDy | NADP-dehydrogenase | redox | redox | 2 |
| AKGDH | decarboxylating dehydrogenase | redox | redox | 2 |
| PDH | decarboxylating dehydrogenase + CoA transfer | redox | redox | 2 |
| SUCDi | quinone-dehydrogenase | redox | redox | 2 |
| MDH2 | quinone-dehydrogenase | redox | redox | 2 |
| MDH3 | menaquinone-dehydrogenase | redox | redox | 2 |
| GLYCTO_2_ | quinone-dehydrogenase | redox | redox | 2 |
| POX | quinone-linked oxidative decarboxylation | redox | redox | 2 |
| DHORD2 | quinone-dehydrogenase | redox | redox | 2 |
| DHORD5 | menaquinone-dehydrogenase | redox | redox | 2 |
| FDH4pp | quinone-dehydrogenase (formate/CO_2_) | redox | redox | 2 |
| FDH5pp | menaquinone-dehydrogenase (formate/CO_2_) | redox | redox | 2 |
| FRD2 | menaquinol-linked reductase | redox | redox | 2 |
| FRD3 | menaquinol-linked reductase | redox | redox | 2 |
| FADRx | flavoprotein reductase | redox | redox | 2 |
| FMNRx2 | flavoprotein reductase | redox | redox | 2 |
| GTHOr | glutathione reductase | redox | redox | 2 |
| NADH16pp | respiratory complex | redox | redox | 2 |
| CYTBO3_4pp | terminal oxidase | redox | redox | 2 |
| CYTBDpp | terminal oxidase | redox | redox | 2 |
| CYTBD2pp | terminal oxidase | redox | redox | 2 |
| NO3R1pp | anaerobic respiration | redox | redox | 2 |
| NTRIR2x | anaerobic respiration, 6-electron | redox | redox | 6 |
| THD2pp | transhydrogenase | redox | redox | 2 |
| NADTRHD | transhydrogenase | redox | redox | 2 |
| ASPO6 | amino-acid oxidase | redox | redox | 2 |
| RNDR1 | ribonucleotide reductase (generic-formula cofactor, thioredoxin) | redox | unscorable | — |
| SHCHD2 | NAD-dehydrogenase (tetrapyrrole biosynthesis) | redox | redox | 2 |
| FEROpp | Fe(II)/Fe(III) redox (metal-excluded) | redox | unscorable | — |
| FE3Ri | Fe(II)/Fe(III) redox (metal-excluded) | redox | unscorable | — |
| SHCHF | metal chelation (metal-excluded) | non-redox | unscorable | — |

**Supplementary Table 2.** Classification outcomes for all 2,712 iML1515 reactions. Fractions of chemically analysable reactions use 2,373 as denominator. Structurally excluded reactions are reported separately.

| Category | Number of reactions | % of all (2,712) | % of analysable (2,373) |
| --- | --- | --- | --- |
| Scorable non-redox | 1,783 | 65.7% | 75.1% |
| Exchange | 331 | 12.2% | — |
| Redox | 270 | 10.0% | 11.4% |
| Generic formula group | 160 | 5.9% | 6.7% |
| Transition metal | 127 | 4.7% | 5.4% |
| Metalloid | 12 | 0.4% | 0.5% |
| Generic formula group + transition metal | 11 | 0.4% | 0.5% |
| Ambiguous formula mapping | 6 | 0.2% | 0.3% |
| Demand | 6 | 0.2% | — |
| Other unsupported element (F) | 2 | 0.1% | 0.1% |
| Biomass pseudoreaction | 2 | 0.1% | — |
| Other unsupported element (Te) | 1 | 0.0% | 0.0% |
| Other unsupported element (Cs) | 1 | 0.0% | 0.0% |

**Supplementary Table 3.** Formula-only mappings left explicitly ambiguous by the algorithm. These cases are excluded from confident redox/non-redox counts because formula and charge admit competing molecular correspondences.

| Reaction ID | Reaction name | Reaction equation | Reason for ambiguity |
| --- | --- | --- | --- |
| NDPK1 | Nucleoside-diphosphate kinase (ATP:GDP) | ATP + GDP ⇌ ADP + GTP | Competing formula-compatible group-transfer/redox correspondences. Left ambiguous |
| NDPK8 | Nucleoside-diphosphate kinase (ATP:dADP) | ATP + dADP ⇌ ADP + dATP | Competing formula-compatible group-transfer/redox correspondences. Left ambiguous |
| VALTA | Valine transaminase | 2-Oxoglutarate + L-valine ⇌ 3-Methyl-2-oxobutanoate + L-glutamate | Competing formula-compatible group-transfer/redox correspondences. Left ambiguous |
| ACOXT | Acetyl-CoA:oxalate CoA-transferase | Acetyl-CoA + oxalate ⇌ acetate + oxalyl-CoA | Competing formula-compatible group-transfer/redox correspondences. Left ambiguous |
| DHAPT | Dihydroxyacetone phosphotransferase | Dihydroxyacetone + phosphoenolpyruvate → dihydroxyacetone phosphate + pyruvate | Competing formula-compatible group-transfer/redox correspondences. Left ambiguous |
| CRNBTCT | Gamma-butyrobetainyl-CoA:carnitine CoA-transferase | Gamma-butyrobetainyl-CoA + L-carnitine ⇌ L-carnitinyl-CoA + gamma-butyrobetaine | Competing formula-compatible group-transfer/redox correspondences. Left ambiguous |

**Supplementary Table 4.** Network, validation and flux summary statistics. This table consolidates reconstruction size, validation performance, topology and the principal *E. coli* condition-specific electron-flux metrics.

| **Metric** | **Value** |
| --- | --- |
| Metabolic model | iML1515 (2,712 reactions, 1,877 metabolites, 1,516 genes) |
| Confident redox reactions | 270 |
| Confident non-redox reactions | 1,783 |
| Ambiguous formula-only mappings | 6 |
| Chemically unscorable reactions | 314 |
| Fully resolved redox reactions | 219 |
| Partially resolved redox reactions | 5 |
| Donor-side-only redox reactions | 14 |
| Acceptor-side-only redox reactions | 32 |
| Matched redox-pair instances | 505 |
| Formula/stoichiometry-inferred matched-pair instances | 210 (41.6%) |
| Distinct biochemical redox couples | 162 total. 150 inferred independently of direct seeding |
| Static graph metabolites | 151 |
| Static graph directed edges | 263 |
| Reaction-resolved static edges | 279 |
| Reactions contributing to static graph | 218 |
| Weak components | 3 |
| Largest component | 147 metabolites (97.4%) |
| Full electron-network communities | 7 (ARI 0.971–1.000 across 10 seeds) |
| Significant community-subsystem tests | 3 / 182 (q<0.05) |
| Common nodes for network comparison | 149 |
| Edge Jaccard on common-node graphs | 0.7090 |
| Weighted-degree Spearman ρ | 0.5774 (P=1.27×10^-14^) |
| Community ARI / NMI | 0.4942 / 0.5318 |
| Glucose uptake, aerobic / anaerobic, taken from Chen et al.^13^ | 8.7 / 14.9 mmol gDW^-1^ h^-1^ |
| Oxygen uptake, aerobic / anaerobic, taken from Chen et al.^13^ | 11.9 / 0 mmol gDW^-1^ h^-1^ |
| Predicted growth, aerobic / anaerobic | 0.5627 / 0.2639 h^-1^ |
| Cumulative electron-transfer activity | 189.568 / 129.921 mmol e^-^ gDW^-1^ h^-1^ |
| Cumulative activity per measured glucose uptake | 21.79 / 8.72 e^-^ per glucose |
| Net source-to-sink electron flux | 68.832 / 61.330 mmol e^-^ gDW^-1^ h^-1^ |
| Net flux per measured glucose uptake | 7.912 / 4.116 e^-^ per glucose |
| Effective transfer depth | 2.754 / 2.118 transfers per net e^-^ |
| Respiratory terminal-delivery fraction | 69.2% / 0% |
| Fermentative terminal-delivery fraction | ~0% / 83.8% |
| NADH relay flux | 55.081 / 60.122 mmol e^-^ gDW^-1^ h^-1^ |
| Ubiquinol-8 relay flux | 47.599 / 0 mmol e^-^ gDW^-1^ h^-1^ |

**Supplementary Table 5.** Carbon-substrate dependence of electron-flux compression and dominant core redox contribution in pFBA-defined *E. coli* states.

| **Substrate** | **Aerobic net e^-^/substrate** | **Anaerobic net e^-^/substrate** | **Compression (%)** | **Dominant core reactions** | **Core e^-^/substrate** | **Core fraction** |
| --- | --- | --- | --- | --- | --- | --- |
| glucose | 11.11 | 4.103 | 63.07 | GAPD | 3.924 | 95.64 |
| fructose | 11.11 | 4.103 | 63.07 | GAPD | 3.924 | 95.64 |
| xylose | 10.33 | 3.399 | 67.09 | GAPD | 3.285 | 96.67 |
| galactose | 11.03 | 4.089 | 62.94 | GAPD | 3.935 | 96.22 |
| gluconate | 10.98 | 2.545 | 76.82 | GAPD;GND | 2.414 | 94.86 |
| glycerol | 7.313 | 4.018 | 45.07 | G3PD2;GAPD | 3.981 | 99.09 |

**Supplementary Table 6.** Reaction-level electron-source contributions calculated from the anaerobic Gonzalez ^13^C-flux maps. Fractions are percentages of total net source-to-sink electron flux. Abbreviations as in Supplementary Table 1.

| **Condition** | **Reaction no.** | **Electron source** | **e^-^ per substrate** | **Fraction of net source** |
| --- | --- | --- | --- | --- |
| Anaerobic glucose | 6 | GAPD | 3.666 | 83.78 |
| Anaerobic glucose | 9 | G6PDH | 0.1573 | 3.595 |
| Anaerobic glucose | 10 | GND | 0.1517 | 3.465 |
| Anaerobic glucose | 22 | PDH | 0.1122 | 2.563 |
| Anaerobic glucose | 25 | ICDH | 0.1035 | 2.364 |
| Anaerobic glucose | 48 | Serine biosynthesis | 0.09089 | 2.077 |
| Anaerobic glucose | 39 | Formate → H_2_ | 0.071 | 1.622 |
| Anaerobic glucose | 63 | Histidine biosynthesis | 0.0161 | 0.368 |
| Anaerobic xylose | 6 | GAPD | 3.237 | 74.48 |
| Anaerobic xylose | 39 | Formate → H_2_ | 0.3609 | 8.303 |
| Anaerobic xylose | 22 | PDH | 0.3394 | 7.808 |
| Anaerobic xylose | 33 | NADP malic enzyme | 0.3076 | 7.076 |
| Anaerobic xylose | 25 | ICDH | 0.04797 | 1.104 |
| Anaerobic xylose | 48 | Serine biosynthesis | 0.04007 | 0.922 |
| Anaerobic xylose | 63 | Histidine biosynthesis | 0.00712 | 0.164 |

**Supplementary Table 7.** Cross-species model provenance and electron-transfer reconstruction quality control.

a, Model provenance and exact bundled input hashes. The SHA256 digest identifies the exact model file analysed.

| **Organism** | **Model** | **Primary DOI** | **Bundled file** | **SHA256** |
| --- | --- | --- | --- | --- |
| *Escherichia coli* K-12 MG1655 | iML1515 | 10.1038/nbt.3956 | iML1515.xml | 9b15b7abbb66b8ac8015c962f7fc6a95d147dfedf4d12a6c201da60b48d8e254 |
| *Saccharomyces cerevisiae* S288C | Yeast9 / yeast-GEM v9.1.0 | 10.1038/s44320-024-00060-7 | yeast-GEM-v9.1.0.xlsx | d589874d9b3ec7662bee206f10aaff86336ada6c50ae2cd6f80a38c23f685759 |
| *Bacillus subtilis* 168 | iBB1018 | 10.3390/ijms24087091 | iBB1018.xlsx | 5635ed8d4d36245e06d1afca5964901a1ce26da0456bebb3dc47dea131871e1b |
| *Salmonella enterica* pan-reactome | iYS1720 | 10.1038/s41467-018-06112-5 | iYS1720.xml.gz | de43ce568b09b78999a6faed3761a1b62e146fc84d96d45d9372c70955120cbd |
| *Komagataella phaffii* (Pichia pastoris) | iMT1026 | 10.1371/journal.pone.0148031 | iMT1026_model.xlsx | 412c0f65ebbf348a950fc432527876d0ddb3ab23cd6955443ba7032420173a4a |

b, Reconstruction quality control. Classified fraction uses chemically analyzable reactions as denominator. Fully resolved fraction uses confidently identified redox reactions as denominator.

| **Organism** | **All rxns** | **Chemical rxns** | **Classified (%)** | **Redox rxns** | **Fully resolved (%)** | **Redox couples** | **ETN nodes** | **ETN reactions** | **ETN edges** | **Largest WCC (%)** |
| --- | --- | --- | --- | --- | --- | --- | --- | --- | --- | --- |
| *E. coli* | 2712 | 2373 | 86.5 | 270 | 81.1 | 162 | 151 | 218 | 263 | 97.4 |
| *S. cerevisiae* | 4102 | 3818 | 96.1 | 292 | 82.2 | 198 | 222 | 240 | 287 | 60.8 |
| *B. subtilis* | 1580 | 1366 | 84.1 | 168 | 70.2 | 112 | 105 | 117 | 157 | 100.0 |
| *S. enterica* | 3357 | 2872 | 91.1 | 296 | 81.8 | 165 | 153 | 238 | 272 | 100.0 |
| *K. phaffii* | 2210 | 2012 | 81.5 | 323 | 72.1 | 187 | 194 | 231 | 266 | 81.4 |

**Supplementary Table 8.** Cross-species physiological conditions and electron-flux observables. Except for the measured *E. coli* glucose/O_2_ constraints, values are derived from the calculated electron flux maps. *K. phaffii* uses a severe low-O_2_ state rather than mathematical zero oxygen. Uptake data are expressed as mmol gDW^-1^ h^-1^ and electron fluxes are expressed as mmol e^-^ gDW^-1^ h^-1^.

| **Organism** | **Condition** | **Glucose uptake** | **O_2_ uptake** | **Growth** | **Cumulative e^-^ / glucose** | **Net e^-^ / glucose** | **Transfer depth** | **NADH relay** |
| --- | --- | --- | --- | --- | --- | --- | --- | --- |
| *E. coli* | aerobic | 8.70 | 11.90 | 0.563 | 21.79 | 7.91 | 2.754 | 55.08 |
| *E. coli* | anaerobic | 14.90 | 0.00 | 0.264 | 8.72 | 4.12 | 2.118 | 60.12 |
| *S. cerevisiae* | aerobic | 10.00 | 22.35 | 0.837 | 21.44 | 11.60 | 1.848 | 78.03 |
| *S. cerevisiae* | anaerobic | 10.00 | 0.00 | 0.162 | 7.82 | 3.97 | 1.973 | 35.38 |
| *B. subtilis* | aerobic | 10.00 | 24.93 | 0.924 | 25.42 | 12.55 | 2.025 | 92.02 |
| *B. subtilis* | anaerobic | 10.00 | 0.00 | 0.052 | 8.18 | 4.10 | 1.994 | 40.09 |
| *S. enterica* | aerobic | 10.00 | 20.00 | 0.964 | 29.26 | 11.07 | 2.644 | 83.64 |
| *S. enterica* | anoxic | 10.00 | 0.00 | 0.202 | 8.57 | 4.11 | 2.082 | 40.20 |
| *K. phaffii* | aerobic | 10.00 | 18.60 | 1.127 | 24.32 | 12.70 | 1.915 | 99.38 |
| *K. phaffii* | severe oxygen limitation | 10.00 | 0.50 | 0.186 | 8.32 | 4.21 | 1.978 | 39.93 |

**Supplementary Table 9.** Network-wide electron-flux redistribution between paired aerobic and low-respiratory-capacity glucose conditions.

| **Organism** | **Jensen–Shannon distance** | **Total-variation distance** | **Union edges** | **Transfer-depth change (%)** |
| --- | --- | --- | --- | --- |
| *E. coli* | 0.772 | 0.721 | 67 | -23.1 |
| *S. enterica* | 0.814 | 0.784 | 67 | -21.3 |
| *B. subtilis* | 0.844 | 0.840 | 67 | -1.5 |
| *S. cerevisiae* | 0.786 | 0.759 | 79 | +6.7 |
| *K. phaffii* | 0.807 | 0.824 | 130 | +3.3 |

Distances are calculated from normalized reaction-resolved donor–acceptor electron-flux distributions on the union of edges in each paired state. Values near 1 indicate strong redistribution. The transfer-depth change is included only to show that similar whole-network redistribution magnitudes can accompany opposite directional responses in pathway depth.

**Supplementary Table 10.** Complete systematic 5-organism × 6-carbon-source screen with model- and condition-specific definitions of the baseline B_s_ and disposal-enabled amplitude A_s_. B_s_ is the calculated net source-to-sink electron flux in the listed low-disposal state. J_high_ is the paired value at an O_2_ uptake upper bound of 20 mmol gDW^-1^ h^-1^. A_s_ = J_high_ − B_s_. All quantities are e^-^ per substrate. All 30 attempted combinations are retained. The 22 feasible pairs have A_s_ = 3.24–8.76 e^-^ per substrate.

| **Organism** | **Model** | **Carbon source** | **Status** | **Low-disposal condition** | **High-disposal condition** | **B_s_** | **J_high_** | **A_s_** |
| --- | --- | --- | --- | --- | --- | --- | --- | --- |
| *E. coli* | iML1515 | glucose | both states feasible | O_2_ upper bound 0 | O_2_ upper bound 20 | 4.103 | 11.110 | 7.007 |
| *E. coli* | iML1515 | fructose | both states feasible | O_2_ upper bound 0 | O_2_ upper bound 20 | 4.103 | 11.110 | 7.007 |
| *E. coli* | iML1515 | galactose | both states feasible | O_2_ upper bound 0 | O_2_ upper bound 20 | 4.089 | 11.035 | 6.945 |
| *E. coli* | iML1515 | xylose | both states feasible | O_2_ upper bound 0 | O_2_ upper bound 20 | 3.399 | 10.327 | 6.929 |
| *E. coli* | iML1515 | gluconate | both states feasible | O_2_ upper bound 0 | O_2_ upper bound 20 | 2.545 | 10.978 | 8.433 |
| *E. coli* | iML1515 | glycerol | both states feasible | O_2_ upper bound 0 | O_2_ upper bound 20 | 4.018 | 7.313 | 3.296 |
| *B. subtilis* | iBB1018 | glucose | both states feasible | O_2_ upper bound 0 | O_2_ upper bound 20 | 4.104 | 10.213 | 6.109 |
| *B. subtilis* | iBB1018 | fructose | both states feasible | O_2_ upper bound 0 | O_2_ upper bound 20 | 4.117 | 10.188 | 6.072 |
| *B. subtilis* | iBB1018 | galactose | both states feasible | O_2_ upper bound 0 | O_2_ upper bound 20 | 4.061 | 10.155 | 6.095 |
| *B. subtilis* | iBB1018 | xylose | both states feasible | O_2_ upper bound 0 | O_2_ upper bound 20 | 3.336 | 9.983 | 6.646 |
| *B. subtilis* | iBB1018 | gluconate | low-disposal state infeasible | O_2_ upper bound 0 | O_2_ upper bound 20 | — | — | — |
| *B. subtilis* | iBB1018 | glycerol | low-disposal state infeasible | O_2_ upper bound 0 | O_2_ upper bound 20 | — | — | — |
| *S. enterica* | iYS1720 | glucose | both states feasible | O_2_ upper bound 0 | O_2_ upper bound 20 | 4.115 | 11.066 | 6.951 |
| *S. enterica* | iYS1720 | fructose | both states feasible | O_2_ upper bound 0 | O_2_ upper bound 20 | 4.115 | 11.066 | 6.951 |
| *S. enterica* | iYS1720 | galactose | both states feasible | O_2_ upper bound 0 | O_2_ upper bound 20 | 4.100 | 10.990 | 6.891 |
| *S. enterica* | iYS1720 | xylose | both states feasible | O_2_ upper bound 0 | O_2_ upper bound 20 | 3.405 | 10.131 | 6.726 |
| *S. enterica* | iYS1720 | gluconate | both states feasible | O_2_ upper bound 0 | O_2_ upper bound 20 | 2.139 | 10.898 | 8.760 |
| *S. enterica* | iYS1720 | glycerol | both states feasible | O_2_ upper bound 0 | O_2_ upper bound 20 | 4.000 | 7.239 | 3.239 |
| *K. phaffii* | iMT1026 | glucose | both states feasible | O_2_ upper bound 0.5 | O_2_ upper bound 20 | 4.207 | 12.699 | 8.492 |
| *K. phaffii* | iMT1026 | fructose | both states feasible | O_2_ upper bound 0.5 | O_2_ upper bound 20 | 4.207 | 12.699 | 8.492 |
| *K. phaffii* | iMT1026 | galactose | low-disposal state infeasible | O_2_ upper bound 0.5 | O_2_ upper bound 20 | — | — | — |
| *K. phaffii* | iMT1026 | xylose | both states feasible | O_2_ upper bound 0.5 | O_2_ upper bound 20 | 5.514 | 12.352 | 6.838 |
| *K. phaffii* | iMT1026 | gluconate | unavailable | O_2_ upper bound 0.5 | O_2_ upper bound 20 | — | — | — |
| *K. phaffii* | iMT1026 | glycerol | low-disposal state infeasible | O_2_ upper bound 0.5 | O_2_ upper bound 20 | — | — | — |
| *S. cerevisiae* | yeast-GEM v9.1.0 | glucose | both states feasible | anaerobic configuration | O_2_ upper bound 20 | 3.966 | 10.806 | 6.840 |
| *S. cerevisiae* | yeast-GEM v9.1.0 | fructose | both states feasible | anaerobic configuration | O_2_ upper bound 20 | 3.966 | 10.806 | 6.840 |
| *S. cerevisiae* | yeast-GEM v9.1.0 | galactose | both states feasible | anaerobic configuration | O_2_ upper bound 20 | 3.984 | 10.787 | 6.804 |
| *S. cerevisiae* | yeast-GEM v9.1.0 | xylose | low-disposal state infeasible | anaerobic configuration | O_2_ upper bound 20 | — | — | — |
| *S. cerevisiae* | yeast-GEM v9.1.0 | gluconate | unavailable | anaerobic configuration | O_2_ upper bound 20 | — | — | — |
| *S. cerevisiae* | yeast-GEM v9.1.0 | glycerol | low-disposal state infeasible | anaerobic configuration | O_2_ upper bound 20 | — | — | — |

For *E. coli*, *B. subtilis* and *S. enterica*, B_s_ uses zero oxygen uptake. *K. phaffii* uses an O_2_ uptake upper bound of 0.5 mmol gDW^-1^ h^-1^. *S. cerevisiae* uses the model-provided anaerobic biomass/nutrient configuration. The stated O_2_ uptake upper bounds are maximum allowed fluxes, not realized uptake. Realized oxygen uptake is reported in results/publication/cross_species_carbon_acceptor_states.csv. All substrates are supplied at 10 mmol gDW^-1^ h^-1^ when the canonical exchange exists.

**Supplementary Table 11.** Complete physiological and optimization constraints and selected pFBA outputs for aerobic and anaerobic glucose growth. Glucose and oxygen uptake are measured constraints. Aerobic non-detected products are fixed to zero. Other product exchanges and internal fluxes are model outputs.

| Metric / setting | Aerobic | Anaerobic | Origin / interpretation |
| --- | --- | --- | --- |
| Glucose uptake qGLC (mmol gDW^-1^ h^-1^) | 8.7 | 14.9 | External constraint. Value reported by Chen et al.^13^ and fixed to the experimental value |
| Oxygen uptake qO_2_  (mmol gDW^-1^ h^-1^) | 11.9 | 0 | External constraint. Value reported by Chen et al.^13^ and fixed to the experimental value |
| D-lactate exchange constraint  (mmol gDW^-1^ h^-1^) | 0 | Not constrained | Aerobic non-detection |
| L-lactate exchange constraint  (mmol gDW^-1^ h^-1^) | 0 | Not constrained | Aerobic non-detection |
| Succinate exchange constraint  (mmol gDW^-1^ h^-1^) | 0 | Not constrained | Aerobic non-detection |
| Formate exchange constraint  (mmol gDW^-1^ h^-1^) | 0 | Not constrained | Aerobic non-detection |
| Ethanol exchange constraint  (mmol gDW^-1^ h^-1^) | 0 | Not constrained | Aerobic non-detection |
| Acetate exchange constraint  (mmol gDW^-1^ h^-1^) | Not constrained | Not constrained | Retained at iML1515 default exchange bounds |
| All other exchange bounds | iML1515 defaults | iML1515 defaults | No condition-specific constraints |
| ATP maintenance reaction (ATPM) lower bound (mmol gDW^-1^ h^-1^) | ≥6.86 | ≥6.86 | Model ATP-maintenance lower bound |
| Biomass objective | Maximise | Maximise | iML1515 core biomass reaction |
| pFBA growth optimum | 100% | 100% | Parsimonious solution at the biomass optimum |
| Predicted growth rate (h^-1^) | 0.563 | 0.264 | pFBA output |
| Predicted acetate exchange  (mmol gDW^-1^ h^-1^) | 8.00 | 12.94 | pFBA output |
| Predicted formate exchange  (mmol gDW^-1^ h^-1^) | 0 | 26.83 | Aerobic value constrained to zero. Anaerobic pFBA output |
| Predicted ethanol exchange  (mmol gDW^-1^ h^-1^) | 0 | 12.80 | Aerobic value constrained to zero. Anaerobic pFBA output |
| Predicted succinate exchange  (mmol gDW^-1^ h^-1^) | 0 | 0.088 | Aerobic value constrained to zero. Anaerobic pFBA output |
| Cumulative electron-transfer activity  (mmol e^-^ gDW^-1^ h^-1^) | 189.57 | 129.92 | Derived from pFBA-weighted electron-transfer reactions |

**Supplementary Table 12.** ECOM4LA measured physiological constraints and state-constrained electron-economy observables.

| **condition** | **qGLC** | **qO_2_** | **Measured growth** | **Predicted growth** | **lactate** | **acetate** | **Net source-to-sink e^-^ flux** | **e^-^ per glucose** | **Effective transfer depth** | **Respiratory fraction** | **Fermentation fraction** |
| --- | --- | --- | --- | --- | --- | --- | --- | --- | --- | --- | --- |
| WT_oxic | 9.02 | 16.49 | 0.71 | 0.6923 | 0 | 3.37 | 93.15 | 10.33 | 2.673 | 0.7081 | 0 |
| ECOM4LA_oxic | 26.4 | 0.21 | 0.32 | 0.2907 | 48.6 | 0 | 109.1 | 4.133 | 2.203 | 0.001768 | 0.8907 |

**Supplementary Table 13a.** ECOM4LA measured endpoint electron-delivery calculation and propagated uncertainty.

| **condition** | **qGLC** | **oxygen_terminal_delivery** | **lactate_terminal_delivery** | **measured_O_2__plus_lactate_delivery** | **measured_delivery_per_glucose** |
| --- | --- | --- | --- | --- | --- |
| WT_oxic | 9.02 | 65.96 | 0 | 65.96 | 7.313 |
| ECOM4LA_oxic | 26.4 | 0.84 | 97.2 | 98.04 | 3.714 |

**Supplementary Table 13b.** ^13^C-metabolic flux data for ECOM4LA^15^ used to calculate electron fluxes.

| **metric** | **unit** | **WT_oxic** | **ECOM4LA_oxic** |
| --- | --- | --- | --- |
| oxidative_PPP_absolute | mmol gDW^-1 h^-1 | 1.4 | 0.5 |
| TCA_contribution_to_OAA | percent | 37.6 | 0.5 |
| anaplerotic_contribution_to_OAA | percent | 62.4 | 99.5 |
| oxidative_PPP_relative_to_glycolysis | percent | 15.1 | 1.7 |

**Supplementary Table 14.** Mean acetate and ethanol formation rates from the Steinsiek respiratory-chain series.

| **strain** | **aerobiosis_percent** | **acetate_mean** | **ethanol_mean** |
| --- | --- | --- | --- |
| TBE029 | 0 | 7.58 | 6.62 |
| TBE029 | 20 | 4.6 | 2.76 |
| TBE029 | 50 | 3.09 | 0.89 |
| TBE029 | 80 | 1.08 | 0.09 |
| TBE029 | 100 | 0.74 | 0.02 |
| TBE029 | 150 | 0.67 | 0.12 |
| TBE031_bo_only | 0 | 7.65 | 7.05 |
| TBE031_bo_only | 20 | 4.17 | 3.48 |
| TBE031_bo_only | 50 | 1.89 | 1.39 |
| TBE031_bo_only | 80 | 0.93 | 0.37 |
| TBE031_bo_only | 100 | 0.61 | 0.45 |
| TBE031_bo_only | 150 | 0.44 | 0.19 |
| TBE032_bdII_only | 0 | 6.9 | 6 |
| TBE032_bdII_only | 20 | 5.84 | 2.86 |
| TBE032_bdII_only | 50 | 4.21 | 0.1 |
| TBE032_bdII_only | 80 | 3.69 | 0.1 |
| TBE032_bdII_only | 100 | 3.27 | 0.07 |
| TBE032_bdII_only | 150 | 3.46 | — |
| TBE042_bdI_only | 0 | 7.74 | 6.82 |
| TBE042_bdI_only | 20 | 4.99 | 3.69 |
| TBE042_bdI_only | 50 | 3.06 | 0.96 |
| TBE042_bdI_only | 80 | 2.52 | 0.12 |
| TBE042_bdI_only | 100 | 2.32 | 0.05 |
| TBE042_bdI_only | 150 | 1.86 | 0.07 |

**Supplementary Table 15.** Summary of the Anand et al. unbranched-ETS extracellular phenotype analysis.

| **architecture** | **state** | **growth_mean** | **growth_sd** | **acetate_carbon_fraction_mean** | **acetate_carbon_fraction_sd** | **aerobicity_rank** |
| --- | --- | --- | --- | --- | --- | --- |
| 1H | evolved | 0.8475 | 0.03105 | 0.318 | 0.004818 | 1 |
| 1H | unevolved | 0.775 | 0.02121 | 0.3139 | 0.0005333 | 1 |
| 2H | evolved | 0.8938 | 0.01506 | 0.221 | 0.007509 | 3 |
| 2H | unevolved | 0.645 | 0.007071 | 0.2218 | 0.004111 | 3 |
| 3H | evolved | 0.8337 | 0.01302 | 0.296 | 0.006174 | 2 |
| 3H | unevolved | 0.425 | 0.007071 | 0.28 | 0.01338 | 2 |
| 4H | evolved | 0.8337 | 0.03021 | 0.1807 | 0.02914 | 4 |
| 4H | unevolved | 0.525 | 0.007071 | 0.1603 | 0.00718 | 4 |

**Supplementary Table 16.** Steinsiek respiratory-architecture predictive-transfer points. Acetate values are mmol gDW^-1^ h^-1^. The 0% and 150% states define each target-specific scale. The 12 intermediate target states are held out.

| **Source architecture** | **Target architecture** | **Aerobiosis (%)** | **Observed acetate** | **Predicted acetate** | **Role** |
| --- | --- | --- | --- | --- | --- |
| TBE029 | TBE031_bo_only | 0 | 7.650 | 7.650 | endpoint used for scaling |
| TBE029 | TBE031_bo_only | 20 | 4.170 | 4.541 | held-out prediction |
| TBE029 | TBE031_bo_only | 50 | 1.890 | 2.965 | held-out prediction |
| TBE029 | TBE031_bo_only | 80 | 0.930 | 0.868 | held-out prediction |
| TBE029 | TBE031_bo_only | 100 | 0.610 | 0.513 | held-out prediction |
| TBE029 | TBE031_bo_only | 150 | 0.440 | 0.440 | endpoint used for scaling |
| TBE029 | TBE032_bdII_only | 0 | 6.900 | 6.900 | endpoint used for scaling |
| TBE029 | TBE032_bdII_only | 20 | 5.840 | 5.416 | held-out prediction |
| TBE029 | TBE032_bdII_only | 50 | 4.210 | 4.665 | held-out prediction |
| TBE029 | TBE032_bdII_only | 80 | 3.690 | 3.664 | held-out prediction |
| TBE029 | TBE032_bdII_only | 100 | 3.270 | 3.495 | held-out prediction |
| TBE029 | TBE032_bdII_only | 150 | 3.460 | 3.460 | endpoint used for scaling |
| TBE029 | TBE042_bdI_only | 0 | 7.740 | 7.740 | endpoint used for scaling |
| TBE029 | TBE042_bdI_only | 20 | 4.990 | 5.204 | held-out prediction |
| TBE029 | TBE042_bdI_only | 50 | 3.060 | 3.919 | held-out prediction |
| TBE029 | TBE042_bdI_only | 80 | 2.520 | 2.209 | held-out prediction |
| TBE029 | TBE042_bdI_only | 100 | 2.320 | 1.920 | held-out prediction |
| TBE029 | TBE042_bdI_only | 150 | 1.860 | 1.860 | endpoint used for scaling |

**Supplementary Table 17.** Low-oxygen central carbon source attribution. Dominant sources are identified from each model state rather than fitted to a common endpoint.

| **Organism** | **Dominant selected source(s)** | **Reaction IDs** | **e^-^ / glucose** | **Fraction of net source flux (%)** | **Total net e^-^ flux** |
| --- | --- | --- | --- | --- | --- |
| *E. coli* | GAPD | GAPD | 3.915 | 95.1 | 61.330 |
| *S. enterica* | GAPD | GAPD | 3.903 | 94.9 | 41.147 |
| *B. subtilis* | GAPD | GAPD | 3.968 | 96.7 | 41.039 |
| *S. cerevisiae* | GAPD + oxidative PPP | r_0486;r_0466;r_0889 | 3.740 | 94.3 | 39.659 |

**Supplementary Table 18.** Independent *E. coli* reconstruction robustness. iML1515 and iJO1366 are compared under the same standardized glucose/O_2_ constraints.

| **Model** | **Condition** | **Growth** | **O_2_ uptake** | **Cumulative e^-^ / glucose** | **Net e^-^ / glucose** | **Transfer depth** |
| --- | --- | --- | --- | --- | --- | --- |
| iML1515 | aerobic | 0.822 | 20.00 | 29.591 | 11.110 | 2.663 |
| iML1515 | anaerobic | 0.158 | 0.00 | 8.640 | 4.103 | 2.106 |
| iJO1366 | aerobic | 0.982 | 17.58 | 28.065 | 10.749 | 2.611 |
| iJO1366 | anaerobic | 0.242 | 0.00 | 8.982 | 4.158 | 2.160 |

**Supplementary Table 19.** Systematic terminal-acceptor capability screen across five reconstructions. Carbon-bearing fumarate is retained for completeness but excluded from the non-carbon pooled correlations.

| **Organism** | **Acceptor** | **Status** | **Carbon-bearing** | **Actual uptake** | **Growth** | **Net e^-^/glucose** | **Δe^-^/glucose** |
| --- | --- | --- | --- | --- | --- | --- | --- |
| *E. coli* | none | baseline | no | 0.000 | 0.158 | 4.103 | — |
| *E. coli* | oxygen | feasible_with_uptake | no | 20.000 | 0.822 | 11.110 | 7.007 |
| *E. coli* | nitrate | feasible_with_uptake | no | 20.000 | 0.684 | 14.771 | 10.667 |
| *E. coli* | nitrite | feasible_with_uptake | no | 20.000 | 0.600 | 14.269 | 10.165 |
| *E. coli* | TMAO | feasible_with_uptake | no | 20.000 | 0.363 | 5.370 | 1.267 |
| *E. coli* | DMSO | feasible_with_uptake | no | 20.000 | 0.363 | 5.370 | 1.267 |
| *E. coli* | fumarate | feasible_with_uptake | yes | 20.000 | 0.380 | 4.854 | 0.751 |
| *E. coli* | thiosulfate | available_but_unused | no | 0.000 | 0.158 | 4.103 | 0.000 |
| *B. subtilis* | none | baseline | no | 0.000 | 0.052 | 4.104 | — |
| *B. subtilis* | oxygen | feasible_with_uptake | no | 20.000 | 0.792 | 10.213 | 6.109 |
| *B. subtilis* | nitrate | feasible_with_uptake | no | 20.000 | 0.338 | 16.945 | 12.841 |
| *B. subtilis* | nitrite | feasible_with_uptake | no | 11.630 | 0.266 | 7.721 | 3.617 |
| *B. subtilis* | fumarate | infeasible_or_failed | yes | — | — | — | — |
| *S. enterica* | none | baseline | no | 0.000 | 0.202 | 4.115 | — |
| *S. enterica* | oxygen | feasible_with_uptake | no | 20.000 | 0.964 | 11.066 | 6.951 |
| *S. enterica* | nitrate | feasible_with_uptake | no | 20.000 | 0.789 | 14.614 | 10.500 |
| *S. enterica* | nitrite | feasible_with_uptake | no | 20.000 | 0.691 | 14.198 | 10.084 |
| *S. enterica* | TMAO | feasible_with_uptake | no | 20.000 | 0.434 | 5.377 | 1.263 |
| *S. enterica* | DMSO | feasible_with_uptake | no | 20.000 | 0.434 | 5.377 | 1.263 |
| *S. enterica* | fumarate | feasible_with_uptake | yes | 20.000 | 0.443 | 5.217 | 1.103 |
| *S. enterica* | tetrathionate | feasible_with_uptake | no | 20.000 | 0.504 | 13.580 | 9.465 |
| *S. enterica* | thiosulfate | feasible_with_uptake | no | 20.000 | 0.331 | 5.148 | 1.033 |
| *K. phaffii* | none | baseline | no | 0.000 | 0.033 | 4.614 | — |
| *K. phaffii* | oxygen | feasible_with_uptake | no | 18.604 | 1.127 | 12.699 | 8.085 |
| *K. phaffii* | nitrate | available_but_unused | no | 0.000 | 0.033 | 4.614 | 0.000 |
| *K. phaffii* | fumarate | feasible_with_uptake | yes | 1.708 | 0.036 | 4.898 | 0.283 |
| *S. cerevisiae* | none | baseline | no | 0.000 | 0.162 | 3.966 | — |
| *S. cerevisiae* | oxygen | feasible_with_uptake | no | 20.000 | 0.769 | 10.806 | 6.840 |
| *S. cerevisiae* | fumarate | feasible_with_uptake | yes | 20.000 | 0.323 | 7.842 | 3.876 |

**Supplementary Table 20.** Descriptive statistics for the extended non-carbon terminal-acceptor analysis.

| **Statistic** | **Value** |
| --- | --- |
| Candidate organism × acceptor capabilities attempted | 24 |
| Feasible capabilities with uptake, total | 21 |
| Feasible non-carbon capabilities with uptake | 17 |
| Pooled Δelectron flux vs nominal terminal e^-^ capacity, Pearson r | 0.350 |
| Pooled Δelectron flux vs ΔCO_2_, Pearson r | 0.858 |
| Pooled Δelectron flux vs ΔCO_2_, Spearman ρ | 0.848 |
| Scaled multi-organism states | 15 |
| Scaled expansion vs scaled oxidation, Pearson r | 0.995 |
| Scaled expansion vs scaled oxidation, Spearman ρ | 0.957 |

**Supplementary Table 21.** Extended central carbon-source coverage using acetate, pyruvate and succinate across the five reconstructions.

| **Organism** | **Substrate** | **Entry point** | **Low feasible** | **High feasible** | **High growth** | **Low growth** | **High net e^-^/substrate** |
| --- | --- | --- | --- | --- | --- | --- | --- |
| *B. subtilis* | pyruvate | pyruvate node | no | yes | 0.3129 | — | 6.314 |
| *B. subtilis* | succinate | TCA-cycle intermediate | no | yes | 0.4556 | — | 8.633 |
| *E. coli* | acetate | acetyl-CoA / two-carbon oxidation | no | yes | 0.2101 | — | 5.393 |
| *E. coli* | pyruvate | pyruvate node | yes | yes | 0.3139 | 0.0050 | 5.913 |
| *E. coli* | succinate | TCA-cycle intermediate | no | yes | 0.4501 | — | 8.146 |
| *K. phaffii* | acetate | acetyl-CoA / two-carbon oxidation | no | yes | 0.1940 | — | 13.240 |
| *K. phaffii* | succinate | TCA-cycle intermediate | no | yes | 0.5284 | — | 13.439 |
| *S. cerevisiae* | acetate | acetyl-CoA / two-carbon oxidation | no | yes | 0.1327 | — | 7.985 |
| *S. cerevisiae* | pyruvate | pyruvate node | no | yes | 0.3045 | — | 5.840 |
| *S. cerevisiae* | succinate | TCA-cycle intermediate | no | yes | 0.4521 | — | 7.907 |
| *S. enterica* | acetate | acetyl-CoA / two-carbon oxidation | no | yes | 0.2343 | — | 5.324 |
| *S. enterica* | pyruvate | pyruvate node | no | yes | 0.3549 | — | 5.940 |
| *S. enterica* | succinate | TCA-cycle intermediate | no | yes | 0.5056 | — | 8.216 |

**Supplementary Table 22a.** Endpoints of the six-substrate × seven-level O_2_ capacity factorial response surface.

| **substrate** | **oxygen_cap** | **growth** | **actual_o2** | **co2** | **net_per_substrate** | **effective_transfer_depth** | **net_per_C** | **net_fraction_gamma** |
| --- | --- | --- | --- | --- | --- | --- | --- | --- |
| glucose | 0 | 0.1575 | 0 | -0.05785 | 4.103 | 2.106 | 0.6839 | 0.171 |
| glucose | 20 | 0.8218 | 20 | 21.76 | 11.11 | 2.663 | 1.852 | 0.4629 |
| fructose | 0 | 0.1575 | 0 | -0.05785 | 4.103 | 2.106 | 0.6839 | 0.171 |
| fructose | 20 | 0.8218 | 20 | 21.76 | 11.11 | 2.663 | 1.852 | 0.4629 |
| galactose | 0 | 0.1358 | 0 | -0.04988 | 4.089 | 2.091 | 0.6815 | 0.1704 |
| galactose | 20 | 0.8019 | 20 | 21.71 | 11.03 | 2.669 | 1.839 | 0.4598 |
| xylose | 0 | 0.09964 | 0 | -0.03659 | 3.399 | 2.081 | 0.6797 | 0.1699 |
| xylose | 20 | 0.7171 | 19.03 | 20.57 | 10.33 | 2.657 | 2.065 | 0.5164 |
| gluconate | 0 | 0.1151 | 0 | 1.366 | 2.545 | 2.014 | 0.4242 | 0.1157 |
| gluconate | 20 | 0.7825 | 20 | 26.67 | 10.98 | 2.683 | 1.83 | 0.499 |
| glycerol | 0 | 0.02796 | 0 | -9.644 | 4.018 | 1.523 | 1.339 | 0.287 |
| glycerol | 20 | 0.4947 | 13.64 | 9.694 | 7.313 | 2.697 | 2.438 | 0.5224 |

**Supplementary Table 22b.** Factorial separability statistics.

| **Statistic** | **Value** |
| --- | --- |
| Rank-one fraction of expansion variance (%) | 99.51 |
| Minimum pairwise correlation of scaled expansion curves | 0.956 |
| Mean pairwise correlation of scaled expansion curves | 0.9857 |
| Raw interaction variance (%) | 4.632 |
| Interaction variance after normalization per substrate carbon (%) | 1.356 |
| Interaction variance after normalization by substrate degree of reduction (%) | 2.181 |

**Supplementary Table 23.** Electron-flux observables extracted from the electron flux maps calculated from Gonzalez et al. ^13^C-flux data. Lower/upper envelopes independently propagate reported marginal reaction-flux limits and are not joint confidence intervals. Flux data are expressed as mmol gDW^-1^ h^-1^ and electron fluxes are expressed as mmol e^-^ gDW^-1^ h^-1^.

| **Condition** | **Substrate uptake** | **Net e^-^/substrate** | **Conservative lower envelope** | **Conservative upper envelope** | **Cumulative e^-^/substrate** | **Transfer depth** | **Absolute net e^-^ flux** | **Matched pFBA e^-^/substrate** |
| --- | --- | --- | --- | --- | --- | --- | --- | --- |
| Aerobic glucose | 8.8 | 8.534 | 8.095 | 8.912 | 18.32 | 2.147 | 75.1 | 11.11 |
| Anaerobic glucose | 13.1 | 4.376 | 4.088 | 5.155 | 10.15 | 2.32 | 57.33 | 4.103 |
| Aerobic xylose | 9.5 | 7.634 | 6.617 | 8.327 | 16.24 | 2.127 | 72.52 | 10.33 |
| Anaerobic xylose | 10.8 | 4.347 | 3.442 | 5.793 | 8.885 | 2.044 | 46.95 | 3.399 |

**Supplementary Table 24a.** Direct like-for-like comparison of electron generation derived from Jouhten et al. *S. cerevisiae* ^13^C-MFA maps with matched yeast-GEM predictions. Experimental values are means ± sample s.d. across two independent cultivations. The model is constrained by measured mean glucose uptake and published mean O_2_ uptake only.

| **Inlet O_2_ (%)** | **Experimental electron generation mean** | **Experimental s.d.** | **Mean glucose uptake** | **Published mean O_2_ uptake** | **Model electron generation** | **Absolute error** | **Relative error (%)** |
| --- | --- | --- | --- | --- | --- | --- | --- |
| 20.9 | 10.28 | 0.255 | 1.075 | 2.70 | 11.98 | 1.699 | 16.5 |
| 2.8 | 10.10 | 0.396 | 1.320 | 2.50 | 10.44 | 0.338 | 3.3 |
| 1 | 6.61 | 0.071 | 2.045 | 1.70 | 6.84 | 0.234 | 3.5 |
| 0.5 | 4.96 | 0.424 | 2.490 | 1.20 | 5.66 | 0.703 | 14.2 |
| 0 | 3.64 | 0.000 | 6.440 | 0.00 | 3.97 | 0.327 | 9.0 |

**Supplementary Table 24b.** Electron generation derived from Baumann et al. *K. phaffii* ^13^C-MFA oxygen-gradient data. Error terms are independently propagated standard deviations.

| **Strain** | **Inlet O_2_ (%)** | **Glucose uptake**  **(mmol gDCW^-1^ h^-1^)** | **Electron generation (mmol e^-^ gDCW^-1^ h^-1^)** | **Electron generation per glucose**  **(e^-^/glucose)** | **S.d.** |
| --- | --- | --- | --- | --- | --- |
| Control | 21 | 0.980 | 8.126 | 8.295 | 0.475 |
| Control | 11 | 1.266 | 9.574 | 7.562 | 0.345 |
| Control | 8 | 1.847 | 9.452 | 5.118 | 0.448 |
| Fab-expressing | 21 | 0.944 | 7.729 | 8.187 | 0.489 |
| Fab-expressing | 11 | 1.312 | 9.727 | 7.412 | 0.482 |
| Fab-expressing | 8 | 1.686 | 8.334 | 4.944 | 0.303 |

**Supplementary Table 25.** Fromanger et al. *C. shehatae* glucose/xylose oxygen-series measurements and calculated whole-cell electron-equivalent partition. Carbon yields are Cmol product per Cmol substrate. OUR values for oxygen-limited phases are reported mean ± s.d. Aerobic values are the reported maximum specific OUR and are shown as point estimates. The electron-to-O_2_ fraction is obtained from product/biomass degree-of-reduction balance and is a whole-cell endpoint-level quantity.

| **Substrate** | **State** | **OUR** | **Ethanol** | **Total polyols** | **Biomass** | **Organic acids** | **e^-^ to O_2_ (%)** |
| --- | --- | --- | --- | --- | --- | --- | --- |
| Xylose | Xylose-1 | 0.80 ± 0.35 | 0.359 | 0.316 | 0.032 | 0.009 | 7.08 |
| Xylose | Xylose-2 | 1.19 ± 0.09 | 0.426 | 0.169 | 0.074 | 0.016 | 8.04 |
| Xylose | Xylose-3 | 2.48 ± 0.77 | 0.322 | 0.150 | 0.094 | 0.018 | 23.39 |
| Glucose | Glucose-1 | 0.30 ± 0.18 | 0.535 | 0.058 | 0.083 | 0.026 | 2.02 |
| Glucose | Glucose-2 | 1.77 ± 0.16 | 0.435 | 0.088 | 0.093 | 0.016 | 13.43 |
| Glucose | Glucose-3 | 2.46 ± 0.91 | 0.394 | 0.156 | 0.048 | 0.031 | 15.91 |
| Xylose | Aerobic ref. | 9.13 (max) | 0.023 | 0.032 | 0.524 | 0.000 | 36.06 |
| Glucose | Aerobic ref. | 7.46 (max) | 0.003 | 0.050 | 0.564 | 0.000 | 32.70 |

**Supplementary Table 26.** Weusthuis et al. *S. cerevisiae* oxygen-series data used for predictive transfer. Ethanol production is calculated from culture and feed ethanol concentrations, biomass and dilution rate. Glucose states define the monotonic response. The maltose zero-O_2_ and 30.8 mmol l^-1^ h^-1^ states define the two endpoint conditions used for scaling. Five intermediate maltose states are held out. Glucose <1 mM ethanol non-detections are shown as “<1 (0 primary)” and treated as 0 mM in the primary analysis.

| **Substrate** | **O_2_ feed** | **Culture ethanol (mM)** | **Biomass (g l^-1^)** | **qethanol (mmol gDW^-1^ h^-1^)** | **Role** | **Predicted q_ethanol_** |
| --- | --- | --- | --- | --- | --- | --- |
| Glucose | 0 | 242 | 2.52 | 9.127 | response inference | — |
| Glucose | 0.7 | 201 | 2.64 | 7.159 | response inference | — |
| Glucose | 0.93 | 240 | 2.70 | 8.444 | response inference | — |
| Glucose | 4.7 | 172 | 2.56 | 6.250 | response inference | — |
| Glucose | 7.3 | 99 | 1.99 | 4.372 | response inference | — |
| Glucose | 13.5 | 58 | 2.07 | 2.222 | response inference | — |
| Glucose | 17.5 | 41 | 2.15 | 1.349 | response inference | — |
| Glucose | 24.2 | 14 | 2.67 | 0.075 | response inference | — |
| Glucose | 33.1 | <1 (0 primary) | 3.18 | -0.377 | response inference | — |
| Glucose | 46.6 | <1 (0 primary) | 2.79 | -0.430 | response inference | — |
| Maltose | 0 | 259 | 2.16 | 11.435 | low O_2_ endpoint replicate | — |
| Maltose | 0 | 259 | 2.13 | 11.596 | low O_2_ endpoint replicate | — |
| Maltose | 5.4 | 211 | 2.40 | 8.292 | held out | 7.710 |
| Maltose | 10.8 | 121 | 2.55 | 4.275 | held out | 4.801 |
| Maltose | 18.7 | 80 | 2.36 | 2.881 | held out | 2.508 |
| Maltose | 22.3 | 41 | 2.51 | 1.155 | held out | 1.738 |
| Maltose | 24.8 | 26 | 2.68 | 0.522 | held out | 1.297 |
| Maltose | 30.8 | 35 | 2.41 | 0.954 | high O_2_ endpoint | — |
