## Supplementary material for "Organizing principles of the cellular electron economy": README.md

# Baumann et al. (2010) *Komagataella phaffii* oxygen-gradient 13C-MFA validation

Source publication:

Baumann K, Carnicer M, Dragosits M, et al. **A multi-level study of recombinant Pichia pastoris in different oxygen conditions.** BMC Systems Biology 4, 141 (2010). DOI: 10.1186/1752-0509-4-141.

Article: https://doi.org/10.1186/1752-0509-4-141

Primary supplementary sources used to calculate electron generation:

- Additional file 5 (published net metabolic fluxes and standard deviations): https://media.springernature.com/original/springer-static/esm/art%3A10.1186%2F1752-0509-4-141/MediaObjects/12918_2010_552_MOESM5_ESM.XLS
- Additional file 8 (stoichiometric model and cofactor stoichiometry): https://media.springernature.com/original/springer-static/esm/art%3A10.1186%2F1752-0509-4-141/MediaObjects/12918_2010_552_MOESM8_ESM.DOC

The article and supplementary material are distributed under the article's CC BY 2.0 licence. The accompanying CSV files provide the machine-readable condition metrics and electron-accounting rules used for the reported analysis.

## Experimental design

The study analysed glucose-limited *Pichia pastoris* (now *Komagataella phaffii*) X-33 control and Fab-expressing strains in chemostats (D = 0.1 h^-1) at 21%, 11% and 8% inlet O2. The 8% condition was described by the authors as hypoxic/pseudo-steady-state.

## Observable boundary

The original 13C-MFA formulation quantifies central-carbon fluxes independently of NADH, NADPH, ATP, O2 and CO2 balances. The repository therefore calculates **electron generation** from the electron-generating reactions resolved by these metabolic flux distributions.

Reduced-cofactor production/consumption is converted using two electron equivalents per NADH, NADPH or FADH2. The source observable is the sum of positive reaction-level electron contributions. Uncertainties are first-order propagated from reported marginal flux standard deviations under an independence assumption.

`baumann2010_condition_metrics.csv` records the condition-level outputs used in the manuscript and Supplementary Figure S14. `baumann2010_electron_accounting_rules.csv` records the accounting conventions and condition-specific biomass coefficients used to calculate electron generation from the published metabolic flux distributions.
