## Supplementary material for "Organizing principles of the cellular electron economy": README.md

# Denby et al. 2015 TMAO validation

Primary source: Denby KJ, Rolfe MD, Crick E, Sanguinetti G, Poole RK, Green J. *Adaptation of anaerobic cultures of Escherichia coli K-12 in response to environmental trimethylamine-N-oxide.* Environ Microbiol. 2015, 17:2477-2491. DOI: 10.1111/1462-2920.12726.

The study used glucose-limited anaerobic chemostats of E. coli K-12 MG1655 with 20 mM glucose at a dilution rate of 0.2 h^-1 and perturbed the fermentative steady state with TMAO. `denby2015_table1_steady_states.csv` transcribes the initial and 1440-min steady-state measurements from Table 1 (means +/- SD, n=3). `denby2015_fig1_redox_balance.csv` transcribes the reducing-equivalent accounting stated in Fig. 1 and its legend.

All publication-facing derived quantities are generated by `scripts/run_acceptor_experimental_validation.py` from the documented inputs.
