## Supplementary material for "Organizing principles of the cellular electron economy": README.md

# Fromanger et al. 2010 native-yeast carbon-entry and oxygen-response validation

Primary source: Fromanger R, Guillouet SE, Uribelarrea JL, Molina-Jouve C, Cameleyre X. *Effect of controlled oxygen limitation on Candida shehatae physiology for ethanol production from xylose and glucose.* Journal of Industrial Microbiology & Biotechnology 37, 437–445 (2010). DOI: 10.1007/s10295-009-0688-7.

`fromanger2010_carbon_partition.csv` transcribes the publication's Tables 1 and 2. Oxygen-limited rows contain the reported mean specific oxygen-uptake rate (OUR) ± SD and carbon yields (Cmol product per Cmol substrate) for ethanol, xylitol, arabitol, ribitol, glycerol, biomass and pooled organic acids (pyruvate + succinate + fumarate). Aerobic-reference rows contain the Table 1 carbon yields, maximum specific OUR, CO2 yield and average respiratory quotient.

The authors report that carbon and degree-of-reduction balances closed within 6% in aerobic growth and within 10% during oxygen limitation. The derived electron-equivalent partition provides a whole-cell endpoint-level validation with the reported experimental closure.

`scripts/run_fromanger2010_electron_balance.py` derives the publication-facing electron partition from these transcribed values. Glucose and xylose both have degree of reduction 4 e- per carbon. Product degree-of-reduction values per carbon are ethanol 6, xylitol/arabitol/ribitol 4.4, glycerol 14/3 and biomass 4.34, using the measured biomass formula C1H2.08O0.66N0.14 reported by Fromanger et al. The pooled organic-acid term is evaluated at gamma/C=3.25 and bracketed from 3.0 (fumarate) to 3.5 (succinate). This changes inferred oxygen delivery by <0.4 percentage points. Aerobic product-balance estimates are independently cross-checked against CO2/RQ.
