## Supplementary material for "Organizing principles of the cellular electron economy": README.md

# Gonzalez et al. (2017) 13C-MFA validation data

This directory contains the numerical inputs used to calculate electron fluxes from the integrated 13C-MFA flux distributions reported in:

Gonzalez JE, Long CP, Antoniewicz MR. *Comprehensive analysis of glucose and xylose metabolism in Escherichia coli under aerobic and anaerobic conditions by 13C metabolic flux analysis.* **Metabolic Engineering 39**, 9–18 (2017). DOI: 10.1016/j.ymben.2016.11.003.

Files used by the analysis:

- `reaction_fluxes.csv`: best-fit reaction fluxes and reported marginal 95% limits for the four glucose/xylose × aerobic/anaerobic conditions. Fluxes are normalized to 100 mol substrate in the source workbook.
- `condition_specific_biomass_reactions.csv`: the four biomass reactions used in the authors' condition-specific 13C-MFA models.
- `condition_metadata.csv`: measured substrate uptake and growth rates transcribed from Table 1 of the paper. Substrate-uptake standard deviations are retained for the decoupling figure.

The processed tables were re-audited against the publisher PDF and original supplementary files used in this analysis. For provenance, the source-file SHA256 digests are:

- publisher PDF `1-s2.0-S1096717616302154-main.pdf`: `496ce590fce146a5ed48d2f2f159f2ae7642b13a970a5ddeab6fb31d48fe18e5`
- `1-s2.0-S1096717616302154-mmc1.xlsx`: `0e370cb5785548e05a15d304080387e70aa6818c2cbc0aa8a9a910849db86f33`
- `1-s2.0-S1096717616302154-mmc2.docx`: `788936ee8a00b2b229dc09c3c81588e4ed2838e91ee54cdc043f045aae5f64a7`
- `1-s2.0-S1096717616302154-mmc3.xlsx`: `2683612002507ef03358210561488846a317922c147a28687a536b8546ee189d`
- `1-s2.0-S1096717616302154-mmc4.xlsx`: `3012277ec569e1bc970a5ee5dc8ec0c703a7b5ce8b56dc1db0c431607991c982`
- `1-s2.0-S1096717616302154-mmc5.xlsx`: `9eb0e031c275bf4308e5bd53d7cf93d78e63eb553d8798898150750d0d66a565`

The derived numerical inputs support the scripted electron-flux calculation. SHA256 hashes above identify the corresponding publisher PDF and original supplementary files for independent transcription checks.
