## Supplementary material for "Organizing principles of the cellular electron economy": README.md

# Jouhten et al. (2008) S. cerevisiae oxygen-gradient 13C-MFA inputs

Primary source: Jouhten P. et al., *BMC Systems Biology* **2**, 60 (2008), DOI 10.1186/1752-0509-2-60.

The open-access source files bundled here are (SHA256 values are recorded in `checksums.sha256`):

- `Jouhten2008_original_Figure2.pdf`: authors' original Figure 2, which reports the two replicate net-flux distributions normalized to 100 glucose and replicate-specific glucose uptake rates.
- `Jouhten2008_Additional_file_4.doc`: published stoichiometric central-carbon model.

`jouhten2008_source_reaction_fluxes.csv` is a direct transcription of the seven electron-generating reaction fluxes used to calculate electron generation. Values are the integer relative fluxes shown in the authors' original Figure 2 (glucose uptake = 100 for each replicate). `jouhten2008_condition_metadata.csv` transcribes the replicate-specific glucose uptake values and reported errors shown in the same figure.

`jouhten2008_reaction_electron_stoichiometry.csv` records the reduced-cofactor electron-source coefficient used for each reaction. The x13 2-oxoglutarate-to-oxaloacetate branch is oriented in the publication's respiratory carbon-flow/electron-donor direction: two NADH plus one FADH2 equivalent are generated (6 e- per turnover). This explicit orientation is required because the compact reaction text in Additional file 4 writes the reduced cofactors on the reactant side, opposite to the respiratory TCA direction shown and discussed in the article.

The analysis script sums positive reduced-equivalent generation for each replicate and reports the condition mean and sample standard deviation (n=2). The resulting means are 10.28, 10.10, 6.61, 4.96 and 3.64 e- per glucose at 20.9, 2.8, 1.0, 0.5 and 0% inlet O2, respectively.
