## Supplementary material for "Organizing principles of the cellular electron economy": README.md

# Perrenoud & Sauer 2005 nitrate/DMSO routing validation

Primary source: Perrenoud A, Sauer U. *Impact of Global Transcriptional Regulation by ArcA, ArcB, Cra, Crp, Cya, Fnr, and Mlc on Glucose Catabolism in Escherichia coli.* J Bacteriol. 2005, 187:3171-3179. DOI: 10.1128/JB.187.9.3171-3179.2005.

`perrenoud2005_acceptor_routing_summary.csv` records the qualitative acceptor comparison described around Fig. 3. Under strict anoxia and with 40 mM DMSO, the TCA cycle remains bifurcated. With 40 mM nitrate, a low but significant respiratory cyclic TCA flux is observed (~0.2 mmol gDW^-1 h^-1).

The experimental panel reports external acceptor concentrations. The standardized model panel applies a common uptake upper bound to compare the qualitative acceptor-specific ordering.
