## Supplementary material for "Organizing principles of the cellular electron economy": run_ecom4la_state_constrained.py

```
"""State-constrained electron-economy analysis of Portnoy et al. 2010 ECOM4LA.

Published external physiological measurements define the WT and ECOM4LA oxic
states and constrain model-derived metabolic flux distributions from which electron fluxes are calculated. The measured
phenotype captures the combined effects of multiple terminal-oxidase deletions,
ygiN deletion and adaptive evolution. Published 13C tracing provides an
independent internal-flux check.
"""
import os, sys, pandas as pd
from pathlib import Path
ROOT=Path(__file__).resolve().parents[3]
sys.path.insert(0,str(ROOT))
from cobra.io import read_sbml_model
from cobra.flux_analysis import pfba
from etn.matching import build_metabolite_table
from etn.electron_flow_analysis import signed_donor_acceptor_edges, RESPIRATORY_ACCEPTORS, FERMENTATION_PRODUCTS
from etn.flow_decomposition import decompose_source_sink_paths

MODEL=str(ROOT/'data/iML1515.xml')
OUT=os.path.dirname(__file__)
PUB=ROOT/'results'/'publication'; PUB.mkdir(parents=True,exist_ok=True)
BIOM='BIOMASS_Ec_iML1515_core_75p37M'
STATES={
 'WT_oxic': dict(qGLC=9.02,qO2=16.49,lactate=0.0,acetate=3.37,measured_growth=0.71),
 'ECOM4LA_oxic': dict(qGLC=26.4,qO2=0.21,lactate=48.6,acetate=0.0,measured_growth=0.32),
}

def solve_state(vals, strict_products=True):
    m=read_sbml_model(MODEL)
    m.reactions.get_by_id('EX_glc__D_e').bounds=(-vals['qGLC'],-vals['qGLC'])
    m.reactions.get_by_id('EX_o2_e').bounds=(-vals['qO2'],-vals['qO2'])
    m.reactions.get_by_id('EX_lac__D_e').bounds=(vals['lactate'],vals['lactate'])
    m.reactions.get_by_id('EX_lac__L_e').bounds=(0,0)
    m.reactions.get_by_id('EX_ac_e').bounds=(vals['acetate'],vals['acetate'])
    if strict_products:
        for rid in ['EX_etoh_e','EX_for_e','EX_succ_e']:
            m.reactions.get_by_id(rid).bounds=(0,0)
    m.objective=BIOM
    opt=m.optimize()
    if opt.status!='optimal': raise RuntimeError(opt.status)
    return m,pfba(m,1.0)

base=read_sbml_model(MODEL); mt=build_metabolite_table(base)
rows=[]; sink_rows=[]
for name,vals in STATES.items():
    m,sol=solve_state(vals,True)
    edges=signed_donor_acceptor_edges(m,mt,sol)
    paths,summ,stats=decompose_source_sink_paths(edges)
    idx=stats.set_index('node')
    resp=ferm=other=unresolved=0.0
    for r in stats.itertuples():
        if r.balance < -1e-9:
            f=-r.balance
            if r.node in RESPIRATORY_ACCEPTORS: resp+=f; fate='respiration'
            elif r.node in FERMENTATION_PRODUCTS or str(r.node)=='unresolved:ACALD': ferm+=f; fate='fermentation_associated'
            elif str(r.node).startswith('unresolved:'): unresolved+=f; fate='unresolved_other'
            else: other+=f; fate='biosynthesis_or_other'
            sink_rows.append(dict(condition=name,sink=r.node,fate=fate,electron_flux=f))
    rows.append(dict(
        condition=name, qGLC=vals['qGLC'], qO2=vals['qO2'], measured_growth=vals['measured_growth'],
        predicted_growth=float(sol.fluxes[BIOM]), lactate=vals['lactate'], acetate=vals['acetate'],
        net_source_to_sink_e_flux=summ['net_source_flux'], e_per_glucose=summ['net_source_flux']/vals['qGLC'],
        cumulative_e_transfer=summ['total_edge_activity'], effective_transfer_depth=summ['effective_transfer_depth'],
        respiratory_delivery=resp, fermentation_associated_delivery=ferm,
        biosynthesis_or_other_delivery=other, unresolved_other_delivery=unresolved,
        respiratory_fraction=resp/summ['net_sink_flux'], fermentation_fraction=ferm/summ['net_sink_flux'],
        nadh_relay=float(idx.loc['nadh_c','relay_throughflow']) if 'nadh_c' in idx.index else 0.0,
        nadph_relay=float(idx.loc['nadph_c','relay_throughflow']) if 'nadph_c' in idx.index else 0.0,
        q8h2_relay=float(idx.loc['q8h2_c','relay_throughflow']) if 'q8h2_c' in idx.index else 0.0,
    ))
    edges.to_csv(os.path.join(OUT,f'{name}_electron_edges.csv'),index=False)
    paths.to_csv(os.path.join(OUT,f'{name}_illustrative_paths.csv'),index=False)

summary=pd.DataFrame(rows)
summary.to_csv(os.path.join(OUT,'ecom4la_state_constrained_summary.csv'),index=False)
pd.DataFrame(sink_rows).to_csv(os.path.join(OUT,'ecom4la_state_constrained_sinks.csv'),index=False)

# Direct experimental endpoint accounting (not equated with balanced genome-scale flux).
endpoint=[]
for name,v in STATES.items():
    o2_e=4*v['qO2']
    lac_e=2*v['lactate']
    endpoint.append(dict(condition=name,qGLC=v['qGLC'],oxygen_terminal_delivery=o2_e,
                         lactate_terminal_delivery=lac_e,
                         measured_O2_plus_lactate_delivery=o2_e+lac_e,
                         measured_delivery_per_glucose=(o2_e+lac_e)/v['qGLC']))
pd.DataFrame(endpoint).to_csv(os.path.join(OUT,'ecom4la_measured_endpoint_electron_delivery.csv'),index=False)

# Published 13C-tracing quantities transcribed from Portnoy et al. 2010 Table 3/text.
pd.DataFrame([
 {'metric':'oxidative_PPP_absolute','unit':'mmol gDW^-1 h^-1','WT_oxic':1.4,'ECOM4LA_oxic':0.5},
 {'metric':'TCA_contribution_to_OAA','unit':'percent','WT_oxic':37.6,'ECOM4LA_oxic':0.5},
 {'metric':'anaplerotic_contribution_to_OAA','unit':'percent','WT_oxic':62.4,'ECOM4LA_oxic':99.5},
 {'metric':'oxidative_PPP_relative_to_glycolysis','unit':'percent','WT_oxic':15.1,'ECOM4LA_oxic':1.7},
]).to_csv(os.path.join(OUT,'ecom4la_published_13C_checks.csv'),index=False)
# Write publication-facing canonical tables.
for fn in ['ecom4la_state_constrained_summary.csv','ecom4la_state_constrained_sinks.csv','ecom4la_measured_endpoint_electron_delivery.csv','ecom4la_published_13C_checks.csv']:
    import shutil; shutil.copy2(os.path.join(OUT,fn), PUB/fn)
print(summary.to_string(index=False))
```
