## Supplementary material for "Organizing principles of the cellular electron economy": README.md

# Steinsiek et al. 2014 respiratory-architecture response data

`steinsiek2014_byproducts_means.csv` contains the mean acetate and ethanol production values transcribed from the published aerobiosis series for the parental TBE029 strain and three single-oxidase architectures.

These physiological measurements test transfer of a response shape across respiratory architectures using acetate formation as the response variable. Electron-flux interpretation follows the framework in the accompanying manuscript.

The source provides condition means. The predictive-transfer workflow reports point estimates together with source-choice, leave-one-architecture-out and source-level-omission sensitivity checks.
