## Supplementary material for "Organizing principles of the cellular electron economy": README.md

# Toya et al. 2012 nitrate 13C-flux validation

Primary source: Toya Y, Nakahigashi K, Tomita M, Shimizu K. *Metabolic regulation analysis of wild-type and arcA mutant Escherichia coli under nitrate conditions using different levels of omics data.* Mol BioSyst. 2012, 8:2593-2604. DOI: 10.1039/C2MB25069A.

`toya2012_table_SII_redox_subset.csv` and `toya2012_table_SII_reduced_product_sinks.csv` are direct transcriptions of the relevant rows of Supplementary Table SII supplied with the study. The first WT flux column uses the publisher’s oxygen-absent notation (rendered as **Ø2** in extracted text). The main article identifies the corresponding flux map as **WT anaerobic**. The repository uses that unambiguous condition name. The strongly fermentative pattern (large pyruvate-formate lyase and ethanol flux, reverse succinate-dehydrogenase direction) is consistent with that assignment. Fluxes are normalized to glucose uptake = 100 and marginal confidence intervals are those reported by Toya et al.

The three publisher supplementary PDFs used during transcription are identified by SHA256 checksums in `source_checksums.sha256` and are available from the article supplementary-material page.

The derived quantity is **electron generation** calculated from the electron-generating reactions in the reported central-carbon metabolic flux distributions.
