## Supplementary material for "Organizing principles of the cellular electron economy": README.md

# Weusthuis et al. (1994) *Saccharomyces cerevisiae* oxygen-series data

Curated values from Weusthuis RA, Visser W, Pronk JT, Scheffers WA & van Dijken JP. Effects of oxygen limitation on sugar metabolism in yeasts: a continuous-culture study of the Kluyver effect. *Microbiology* 140, 703–715 (1994). DOI: 10.1099/00221287-140-4-703.

`s_cerevisiae_oxygen_series.csv` contains the glucose and maltose chemostat series used for the predictive-transfer analysis. Ethanol values reported as `<1 mM` remain encoded as censored values in the input. The primary analysis treats glucose non-detections as 0 mM culture ethanol and reports sensitivity across the 0–1 mM censoring interval.
