## Supplementary material for "Organizing principles of the cellular electron economy": README.md

# Figure Source Data

Generated by `scripts/make_source_data.py` from the repository analysis outputs. Figure 1 is conceptual and has no numerical Source Data. Reconstruction diagnostics are documented in Supplementary Fig. S1.

## Main figures
- Fig. 2: `Fig2*` files.
- Fig. 3: `Fig3*` files. Factorial statistics are provided with Supplementary Fig. S12.
- Fig. 4A-D: cross-species and Jouhten files `Fig4a_*` to `Fig4d_*`.
- Fig. 4E: `Fig4e_Steinsiek2014_predictive_transfer.csv`.
- Fig. 4F: `Fig4f_Weusthuis1994_predictive_transfer.csv` and `Fig4f_Weusthuis1994_prediction_curve.csv`.

## Supplementary figures
Supplementary Fig. S2 presents the reconstructed iML1515 electron transfer network and is supported by node and edge Source Data tables. Quantitative source files are named `SuppFigS#_*`. Fromanger whole-cell electron-equivalent balances support Supplementary Fig. S18. Respiratory-architecture transfer is documented in Supplementary Fig. S19. Weusthuis carbon-source transfer is documented in Supplementary Fig. S20.

## Observable boundaries
Gonzalez intervals are conservative marginal-limit envelopes. Jouhten, Toya and Baumann quantify electron generation calculated from published metabolic-flux maps. Fromanger provides a whole-cell degree-of-reduction endpoint balance. The Steinsiek and Weusthuis analyses test response transfer using measured acetate and ethanol production as physiological readouts. ECOM4LA calculations complete measured physiological states in the genome-scale model. The Steinsiek analysis uses published condition means and reports point estimates together with source-choice and omission sensitivity tests.
