## Supplementary material for "Organizing principles of the cellular electron economy": README.md

# Cellular electron economy — Supplementary Data

This package contains the machine-readable inputs and outputs supporting the manuscript and Supplementary Information.

## Contents

- `canonical_results/`: analysis outputs used for reported tables, statistics and figures.
- `experimental_inputs/`: curated experimental source material, transcriptions, provenance notes and checksums.
- `figure_source_data/`: the 72 figure-facing CSV files for main and supplementary quantitative panels.
- `FIGURE_SOURCE_DATA_MANIFEST.csv`: panel-to-source-data mapping.
- `INPUT_PROVENANCE_SHA256.csv`: repository input provenance and checksum registry for primary inputs.
- `RESULTS_MANIFEST.md`: guide to canonical result files.
- `REPRODUCIBILITY_WORKFLOW.md`: commands and environment for complete reproduction.

The repository is the executable companion to this data package. Run `python scripts/run_all.py` from the repository root to generate the canonical outputs and figures, then run `pytest -q` to execute the automated tests.

The complete systematic five-organism by six-carbon-source screen retains all 30 attempted combinations. Twenty-two pairs support both low- and high-disposal states. The expansion range across those pairs is 3.24–8.76 e⁻ per substrate.
