## Supplementary material for "Organizing principles of the cellular electron economy": REPRODUCIBILITY_WORKFLOW.md

# Reproducibility workflow

Run from the repository root in the environment specified by `requirements.txt`.

## Core reconstruction and E. coli analyses

```bash
python scripts/validate_reduction_degree.py
python scripts/matching_validation.py
python scripts/run_validation.py
python scripts/validation_battery.py
python scripts/run_analysis.py
python scripts/coverage_analysis.py
python scripts/run_electron_flow_analyses.py
python scripts/run_electron_path_decomposition.py
python scripts/compare_networks.py
python scripts/analyse_community_resolution_sensitivity.py
python scripts/analyse_community_robustness.py
python scripts/analyse_network_comparison_sensitivity.py
python scripts/analyse_flux_robustness.py
```

## Experimental and perturbation analyses

```bash
python scripts/run_gonzalez_13c_mfa_validation.py
python scripts/run_jouhten_13c_mfa_validation.py
python scripts/run_jouhten_matched_model_validation.py
python scripts/run_acceptor_experimental_validation.py
python scripts/run_mechanistic_analysis.py
python data/experimental/portnoy2010_ecom4la/run_ecom4la_state_constrained.py
python scripts/run_respiratory_architecture_support.py
python scripts/run_factorial_carbon_disposal_analysis.py
python scripts/run_fromanger2010_electron_balance.py
python scripts/run_baumann2010_13c_mfa_validation.py
python scripts/run_steinsiek2014_predictive_transfer.py
python scripts/run_weusthuis1994_predictive_transfer.py
```

## Cross-species analyses

```bash
python scripts/run_cross_species_analysis.py
python scripts/run_cross_species_principle_analysis.py --organism "B. subtilis"
python scripts/run_cross_species_principle_analysis.py --organism "S. enterica"
python scripts/run_cross_species_principle_analysis.py --organism "K. phaffii"
python scripts/run_cross_species_principle_analysis.py --merge
python scripts/run_extended_acceptor_screen.py
python scripts/run_extended_carbon_entry_screen.py
python scripts/run_cross_species_routing_redistribution.py
```

## Source Data, figures and tests

```bash
python scripts/make_excel_table.py
python scripts/make_source_data.py
python scripts/make_figures.py
pytest -q
```

`make_figures.py` reproduces computational main Figs. 2–4 through `scripts/main_figures.py` and generates Supplementary Figs. S1–S20. Supplementary Fig. S2 presents the iML1515 electron transfer network generated by `scripts/make_ecoli_etn_network_figure.py`. Main Fig. 4e and Supplementary Fig. S19 use the Steinsiek respiratory-architecture predictive transfer. Main Fig. 4f and Supplementary Fig. S20 use the Weusthuis carbon-source predictive transfer. Fromanger whole-cell balances generate Supplementary Fig. S18.
