## Supplementary material for "Organizing principles of the cellular electron economy": RESULTS_MANIFEST.md

# Results manifest

This manifest groups the principal repository outputs supporting **Organizing principles of the cellular electron economy**. Panel-level mappings are provided in `FIGURE_SOURCE_DATA_MANIFEST.csv`.

- Core reconstruction, validation and E. coli outputs: `results/`.
- Publication analyses: `results/publication/`.
- Cross-species outputs: `results/cross_species/` and `results/publication/cross_species_*`.
- Fromanger whole-cell balance: `results/publication/fromanger2010_electron_balance.csv` and summary JSON.
- Baumann K. phaffii electron-generation validation: `results/publication/baumann2010_13c_mfa_*`.
- Steinsiek respiratory-architecture predictive transfer: `results/publication/steinsiek2014_*`.
- Weusthuis predictive transfer: `results/publication/weusthuis1994_*`.
- Factorial carbon-entry × electron-disposal analysis: `results/publication/factorial_*`.
- Extended terminal-acceptor and carbon-entry screens: `results/publication/extended_*`.
- Whole-network routing redistribution: `results/publication/cross_species_routing_redistribution*`.
- Figure-facing Source Data: `source_data/publication/`.
- Manuscript figures: `figures/publication/Fig2_*` to `Fig4_*`.
- Reconstructed iML1515 electron transfer network: `figures/publication/SuppFigS2_iML1515_ETN.png`, `.pdf` and `.svg`.
- Supplementary Fig. S2 generation script: `scripts/make_ecoli_etn_network_figure.py`.
- Supplementary Fig. S2 node and edge tables: `source_data/publication/SuppFigS2_iML1515_ETN_nodes.csv` and `source_data/publication/SuppFigS2_iML1515_ETN_edges.csv`.
- Boundary-node network representation: `results/iML1515_ETN_boundary_nodes.xml`.
- Quantitative supplementary figures: `figures/publication/SuppFigS1_*` to `SuppFigS20_*`.
