## Supplementary figures and images for "Organizing principles of the cellular electron economy"

### Jouhten2008_original_Figure2.pdf

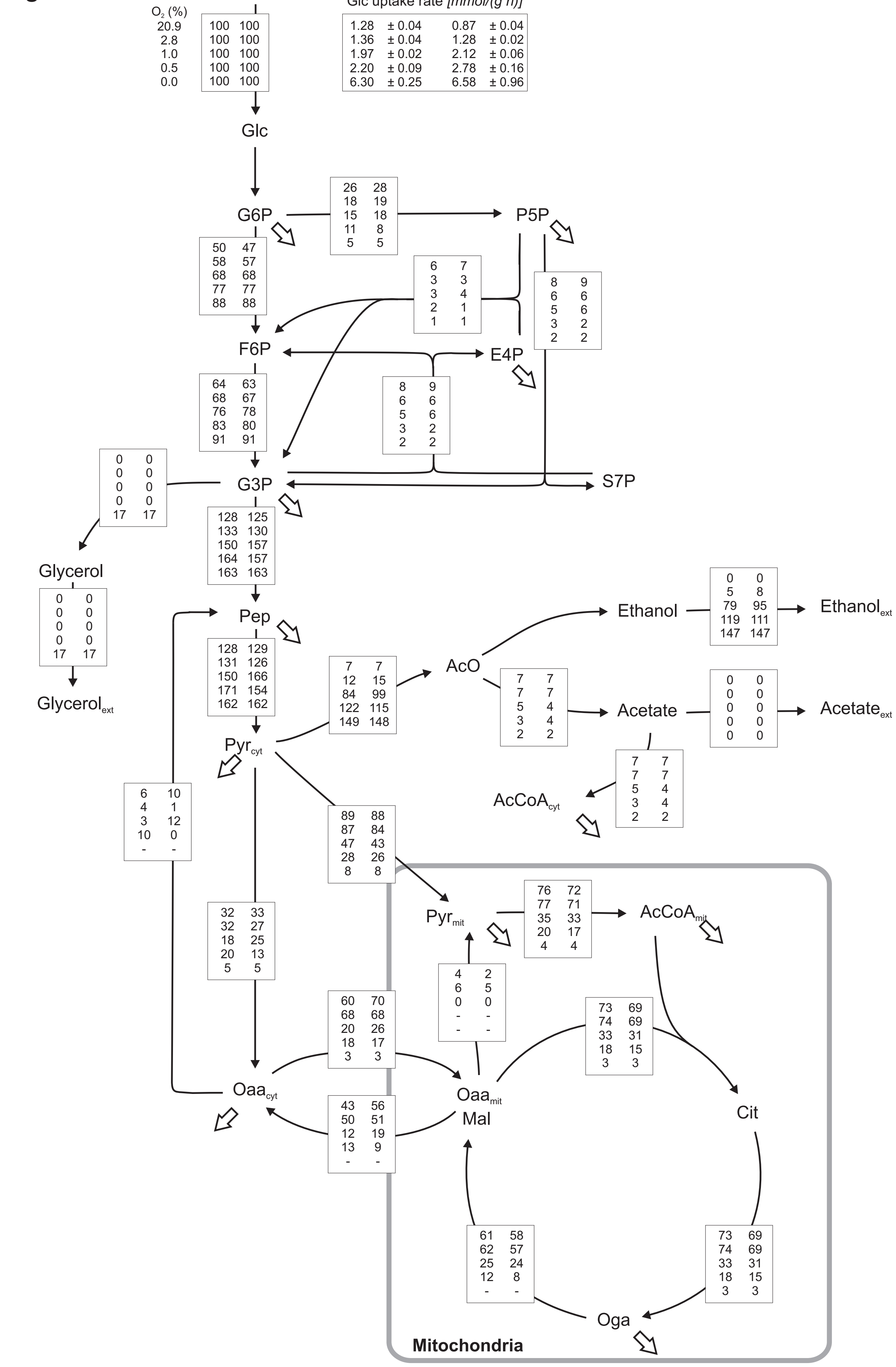
